# TOGGLE Unveils Functional Heterogeneity and Epigenetic Memory in Single Cells

**DOI:** 10.1101/2025.01.01.631041

**Authors:** Junpeng Chen, Tianda Sun, Tian Song, Zhouweiyu Chen, Haowei Xu, Zixuan Guo, Eric Jiang, Yibing Nong, Tao Yuan, Charles C. Dai, Yexing Yan, Jinwen Ge, Haihui Wu, Tong Yang, Shanshan Wang, Zixiang Su, Park Tian, Xiao Yang, Ahmed Abdelbsset-Ismail, You Li, Changping Li, Richa A. Singhal, Kailin Yang, Lu Cai, Zixin Wu, Alex P. Carll

## Abstract

Cells that appear transcriptionally identical can maintain vastly different functions or fates—an enduring blind spot in single-cell transcriptomics. Here we introduce TOGGLE, a self-supervised graph diffusion framework that delineates fine-grained functional heterogeneity within phenotypically stable cell populations. By combining deep diffusion learning with reinforcement-guided clustering and a BERT-inspired masking strategy, TOGGLE reconstructs hidden trajectories of cell fate without prior labels or temporal information. Applied across multiple single-cell RNA-seq datasets, TOGGLE achieves up to 90 % accuracy in unsupervised fate prediction, surpassing existing trajectory algorithms such as WOT and Cospar. It distinguishes ferroptotic, apoptotic, and intermediate neuronal states in ischemic stroke, with predictions experimentally validated in animal models. In neural stem cells TOGGLE reveals epigenetic memory of metabolic activity, linking local DNA demethylation and chromatin accessibility to mitochondrial RNA expression. We further introduce the Graph Diffusion Functional Map, which isolates subtle RNA functional groupings otherwise obscured by conventional dimensionality reduction. TOGGLE thus establishes a generalizable framework for mapping functional identity and epigenetic dynamics at single-cell resolution, providing new insights into cellular memory, regeneration, and disease mechanisms.

**Biographical Note:** Chen’s guidance comes from Dr. Alex P. Carll. Another mentor is Dr. Lu Cai, one of the contributors to the standardization of autophagy detection methods, whose research on criteria for identifying diabetes induced apoptosis has been included in numerous clinical guidelines. Professor Jinwen Ge is a State Council Special Allowance Expert of China. He has been involved in formulating medical standards for integrated traditional Chinese and Western medicine in China and has participated in developing retrieval/indexing norms for journals in the field of integrated traditional Chinese and Western medicine.

**Code and Data availability:** The raw data for the hematopoiesis dataset can be accessed at the Gene Expression Omnibus database with accession number GSE140802, the reprogramming dataset with accession number GSE99915, the e-cigarette aerosol inhalation on mouse cardiac tissue with accession number GSE183614, the rat myocardial infarction dataset with accession number GSE253768, the ischemic stroke dataset with accession number GSE232429, and the adult NSC lineage dataset with accession number GSE209656 and GSE211786.

All codes and guide page & analyzed data are full open available at: https://github.com/FullBlackWolf/TOGGLE?tab=readme-ov-file https://fullblackwolf.github.io/TOGGLE/

**Key points:**

(1) Previous studies have primarily focused on distinguishing cell types and different cellular subpopulations, whereas our tool is designed to identify subtle functional differences among highly similar cell states. For example, in Test2, it first isolates neurons undergoing cell death, then further differentiates early, intermediate, and late death stages, and separates ferroptosis from apoptosis as two major categories. Notably, ferroptosis and apoptosis share a large number of molecular pathways, making this task highly challenging. In addition, in Test4, our method discriminates fibroblasts with distinct immune-communication patterns, even though they do not show prominent separation or temporal progression on conventional UMAP plots.
(2) We developed an innovative Graph Diffusion Functional Map that significantly reduces noise, resulting in clearer visualization of RNA functional groupings and enabling the detection of subtle functional differences in high-dimensional data. This provides a novel tool for pathway discovery. For instance, in Test4, we found that the KEGG inflammatory response pathway was most strongly activated in young mice, suggesting that e-cigarette exposure exerts more severe effects on immature cardiac tissue.
(3) We implemented a novel trajectory-tracking strategy capable of distinguishing early, intermediate, and late stages of cellular programming, which we refer to as functional lineage tracing. In Test3, we successfully separated nearly indistinguishable aNSC and qNSC populations and further uncovered their methylation-regulated transcriptional programs, demonstrating that local DNA demethylation activates genes involved in cell proliferation and energy metabolism, thereby regulating the activation of quiescent neurons.
(4) In the testing dataset, it was found that local DNA demethylation activates the expression of genes associated with cell proliferation and energy metabolism, thereby regulating the activation of quiescent neurons. Neuronal ferroptosis in ischemic stroke may be linked to the high expression of ferroptosis driver genes such as Ctsb, Mtdh, Ndrg1, Smad7, Cd82, and Acsl1.

## 1. Introduction

RNA plays a central regulatory role throughout the life cycle of a cell—from formation and development to maturation, functional execution, aging, and pathological transformation. Dynamic changes in RNA expression precisely orchestrate cellular function at each stage. To capture these transitions, gene expression profiles across distinct cellular populations are typically compared. However, even within a single cell type, functional divergence or fate transitions may occur without significant transcriptomic differences, leading to a critical challenge: phenotypically similar cells may exist in vastly different functional states that are indistinguishable using standard expression-based methods. A salient example arises in ischemic stroke, where current single-cell RNA sequencing (scRNA-seq) approaches can distinguish broad neuronal subtypes (e.g., glutamatergic neurons) but fail to resolve neurons undergoing distinct stages of programmed cell death^1^. Similarly, neural stem cells exhibit marked DNA methylation shifts upon activation, yet their transcriptomic profiles remain nearly identical before and after transition, eluding traditional RNA-based classification^2^.

We refer to this integrative methodology as Fate Tracing—a conceptual and computational framework for reconstructing the historical trajectories of functional transitions and programmed cellular processes. It is necessary to use an unsupervised algorithm because the outcomes of costly cellular experiments are typically unknown in advance—otherwise, the algorithm would lose its purpose. Although numerous computational tools have been developed for scRNA-seq analysis, most are designed to model developmental trajectories or lineage differentiation, and offer limited resolution for identifying functional transitions within terminally differentiated cells, such as neurons. Current algorithms are primarily represented by WOT^3^ and Cospar^4^, Weighted Optimal Transport (WOT) relies on temporal information to infer cell trajectories but often confounds similar transitional paths. Cospar constructs state-transition models based on diffusion principles, yet lacks the capacity to resolve multi-stage classifications or fine-grained transitions within phenotypically stable populations. A shared limitation among these methods is their inability to distinguish between functionally distinct yet transcriptionally similar mature cell states.

Current trajectory inference tools are typically represented by WOT and Cospar. Tools such as WOT (Weighted Optimal Transport) and Cospar, which rely on diffusion or optimal transport principles, primarily depend on pronounced transcriptional differences or well-defined time-series information. The WOT algorithm requires predefined cell type labels as input and cannot separately perform subpopulation-type analysis (for example, distinguishing CD4 T cells from CD8 T cells) or functional analysis (such as determining whether fibroblasts activate immune-related functions or secretory functions). Cospar, in its current implementation, is limited to binary classification tasks. For instance, during programmed cell death, different death modalities (such as ferroptosis and apoptosis) often share highly overlapping marker genes and pathways, and transcriptional changes tend to occur in a sudden, burst-like manner rather than gradually. As a result, these algorithms frequently confuse similar transition paths, struggle to resolve multi-stage fine-grained transitions, and have difficulty distinguishing the dominant mode of death within phenotypically stable populations of dying cells.

Similar limitations appear in other cell dynamics tools. Conventional clustering methods, such as those implemented in Seurat, and pseudotime analysis tools, such as Monocle, excel at broad cell type classification and reconstruction of linear trajectories. However, in scenarios involving programmed cell death, they typically capture only coarse binary states, such as “surviving versus dying” or “activated versus resting.” These approaches struggle to identify overlapping biological programs—for example, the coexistence and competition between ferroptosis and apoptosis pathways—when the overall transcriptional profile remains largely unchanged. They are also unable to resolve tree-like functional hierarchies, particularly failing to further subdivide dying cell populations into ferroptosis-dominant and apoptosis-dominant subgroups. This limitation arises because such methods rely heavily on pronounced transcriptome differences or smooth pseudotime gradients. In contrast, programmed cell death processes frequently feature an early “transcriptionally silent” phase followed by highly overlapping signals in later stages, resulting in severely reduced resolution within dying cells using traditional tools. At present, there is also a lack of methods capable of distinguishing stages of cellular programmed functions—for instance, identifying cells approaching death and then pinpointing late-stage ferroptosis subtypes among them.

To address these limitations, we developed TOGGLE, a computational framework tailored to delineate functional heterogeneity within mature cell populations. We conceptualize cellular function as a hierarchical system: core biological processes, such as metabolism and material transport, form the “trunk,” while specialized functions emerge as branching “limbs.” TOGGLE captures this hierarchy by applying a graph diffusion algorithm to construct multi-level representations of RNA regulatory networks. Coupled with a self-supervised learning scheme, the framework autonomously detects critical functional boundaries and enables fine-grained partitioning of cell populations. The resulting diffusion-derived modules reveal transcriptional specialization that traditional clustering methods fail to capture. TOGGLE thereby establishes a high-resolution functional hierarchy that facilitates discrimination not only between broad cell classes but also among subtle functional subtypes. It further disentangles overlapping biological programs that obscure classification boundaries, enabling precise identification of key transitional states along cell fate trajectories. To operationalize this approach, we constructed an RNA-based functional graph model for visualizing and quantifying transcriptional function.

## 2. Methods

### 2.1 Issues related to the definition of the new concept of Fate Tracing

Fate Tracing builds upon and extends the principles of Lineage Tracing. Traditional Lineage Tracing focuses on tracking cell lineages and identifying broad cell subtypes, such as distinguishing different cell types or branching points in developmental trajectories. In contrast, Fate Tracing refines the analysis to the level of functional micro-tuning within a single cell type, enabling a detailed characterization of functional heterogeneity. We introduce TOGGLE as a novel tool to establish the concept of Fate Tracing, addressing and resolving several newly emerging challenges:

(1) Endpoint loss. In previous tools, both starting and endpoint information for cell fates must be provided simultaneously. For example, this requires progenitor cells—such as granulocyte-monocyte progenitor cells (GMP)—that can generate descendant clones, along with samples capturing those descendant clones, such as neutrophils and monocytes. However, in certain diseases, samples of cells that have reached their final endpoint cannot be obtained. For instance, in cerebral infarction, neurons that have completed programmed cell death cannot be captured.
(2) Temporal loss. The aforementioned algorithms require sampling at multiple distinct time points to reconstruct continuous and gradual cellular changes, thereby inferring transitional trajectories during progenitor cell differentiation. However, in clinical practice, obtaining samples at scheduled time intervals is often challenging—for example, postoperative tumor tissue, hematoma following cerebral hemorrhage, or samples from deceased individuals or aborted embryos. Furthermore, ethical constraints severely limit the number of repeated surgical samplings from the same human subject at different time points. As a result, collecting samples across multiple time points becomes infeasible. This necessitates increasing the analytical depth of single-time-point samples. Achieving this requires first distinguishing cells at different stages of fate progression with high classification accuracy, and then using these precise classifications to predict cellular fate.
(3) Sample imbalance. When samples are collected at multiple time points, it becomes possible to capture the starting point, endpoint, and intermediate stages of cell fate, along with dynamic information on changes in their proportions. At a single time point, however, it is difficult to achieve a balanced representation of cells across different stages of fate progression, resulting in data scarcity and uneven sample distribution for certain stages. This challenge requires a fundamental shift in algorithmic design: moving away from constructing state transitions across multiple time points and instead reconstructing continuous trajectory information from a single time point alone.
(4) Absence of clear fate boundaries. Between progenitor cells and their differentiated descendants—such as neutrophils and monocytes—there exist well-defined fate boundaries. The distinct fate directions taken by progenitor cells lead to clear transcriptional differences. In contrast, the process of ferroptosis represents a unidirectional and gradual fate progression that lacks any distinct boundary. As a result, cells undergoing ferroptosis become intermixed with healthy cells and with other cells exhibiting varying expression levels. This intermingling makes the construction of fate trajectories substantially more challenging.

During the testing process, we did not utilize the aforementioned known information, even though such information was present in the original dataset.

Task 1: Stage-specific classification. This task simultaneously incorporates both progenitor cells and their differentiated descendants from the dataset. A single progenitor cell type can differentiate into multiple distinct lineages. For example, hematopoietic progenitor cells may develop into either monocytes or neutrophils. Accordingly, neutrophil progenitor cells represent an early stage of progression toward neutrophil differentiation, while monocyte progenitor cells represent an early stage of progression toward monocyte differentiation. Fully differentiated neutrophils and monocytes constitute the late stages in which the respective differentiation functions have been completed. This task also serves to evaluate the algorithm’s performance in multi-class classification.

Task 2: Lineage tracing classification. Under the condition that Task 1 is correct, it is necessary to maintain high original performance on the lineage tracing task—that is, to separately isolate progenitor cells with different developmental tendencies with high accuracy. This means that the performance of Task 1 is built upon a correct foundation. At this point, progenitor cells with different developmental tendencies imply that the progenitor state can be defined as a functional stage, allowing internal functions to be distinguished within the same selected cell stage.

In the tasks described above, subtle differences observed in single-cell RNA sequencing primarily depend on the tendencies of fate-related functions, rather than on cell type definitions. This characteristic also renders it a challenging classification problem. By employing similar tasks to evaluate the algorithm’s accuracy, we adapt the fate tendencies of developing cells to the functional tendencies of terminally differentiated cells. This adaptation facilitates the migration of the lineage tracing concept to the new definition embodied in Fate Tracing. After defining this problem, we compare it with previous lineage tracing methods. Lineage tracing can be viewed as functional tracing confined to developmental scenarios. However, our algorithm differs by emphasizing the inclusion of cell analysis during non-differentiated periods, as well as performing internal distinctions within early, middle, and late stages. We therefore select TOGGLE, Cospar, Super OT, WOT, GAN-Based OT, and Deep-Lineage (Supervised) as comparison methods. These methods represent the complete technological evolution in this field in recent years, spanning from the traditional optimal transport (Optimal Transport, OT) framework to supervised enhancement, generative adversarial approaches, and deep learning end-to-end prediction. By conducting systematic comparisons on the same datasets, it becomes possible to comprehensively evaluate the relative advantages and limitations of the proposed methods—such as TOGGLE or Deep-Lineage—when addressing real biological challenges.

These include early fate bias detection, multi-stage classification, fate subdivision in static mature cells, robustness to clone sparsity or downsampling, and fine distinctions of functions or fate programs within the same cell type.

Specifically, WOT serves as a classic unsupervised optimal transport benchmark from its peak period, representing the early mainstream approach that relies on aligning time-series distributions. Cospar—one of the currently recognized state-of-the-art methods—significantly enhances the accuracy and robustness of integrating lineage barcodes with transcriptome information through the introduction of coherence and sparsity optimizations, and has become a core reference that most new methods must surpass. Super OT and GAN-Based OT embody improvement directions that combine the optimal transport framework with supervised learning or generative adversarial mechanisms, aimed at alleviating issues related to nonlinear correspondences and distribution drifts. TOGGLE is specifically optimized for multi-programmed stages within the same mature cell type and for static differentiation scenarios, filling gaps left by previous methods in handling non-unique correspondences or mature cell classification. All algorithms employ the initial parameters provided by their original authors.

### 2.2 Method for distinguishing cellular functions and functional phases

The model consists of three progressively layered modules, each exhibiting distinct differences in functional positioning and analytical objectives. For ease of understanding, the following illustrates this with a hypothetical scenario. Suppose there are a total of 10,000 cells, all originating from the unactivated progenitor cell A. Upon activation, progenitor cell A transforms into secretory cell type A1 and conductive cell type A2, though the proportions among the three remain unknown. Furthermore, due to underlying unknown pathological mechanisms, the appropriate sampling time window is also indeterminable, with only the knowledge that A, A1, and A2 coexist within the same sample. At the same time, the functional states of these cells also vary; for example, some cells are undergoing autophagy, others are engaged in sugar metabolism, and still others are prepared to initiate functional remodeling.

The first step involves the data preprocessing phase. The raw input provided by the database consists of complete RNA sequencing data, such as the GAPDH sequence AGGTUGTA, among others.

A typical RNA-seq experiment usually contains billions of reads, each 150 base pairs in length. Even after encoding, the data dimensions remain extremely high, making direct effective computation challenging. To address this, the present study employs commonly used processing strategies in the field. It begins by utilizing preset alignment and counting workflows in R language toolkits to compress the raw sequence data into a gene-level expression count matrix. Subsequently, a graph diffusion formula is used to generate an asymmetric cell similarity matrix. The choice of the graph diffusion formula over traditional metrics, such as the Pearson correlation coefficient, stems from the fact that the latter creates excessively large invalid symmetric regions in the data space. This leads to an overconcentration of effective information, significantly reducing the algorithm’s stability and generalization capability. In summary, Section 2.2.1 describes the graph diffusion module. This module is responsible for converting raw sequencing data into an expression matrix and calculating pairwise similarities between cells, thereby establishing a data foundation for downstream classification and clustering analyses.

The second step encompasses the cell network reprogramming and grouping phase, which addresses the network rewiring problem^29^. Following the completion of the first step, the algorithm produces an unordered matrix M1 of size 10,000 x 10,000, where the arrangement of cells is disorganized and randomly distributed. At this juncture, cells with similar functional states must be preferentially consolidated into adjacent positions—for instance, separately aggregating cells that have just initiated secretion, those already in the secretion process, and those in late-stage secretion. These clustered units correspond to distinct stages within the functional continuum. The output of this step is a sorted matrix M2 of size 10,000 x 10,000, accompanied by the identification of cluster boundaries. For example, cells numbered 1 to 500 are assigned to the early secretory precursor stage of cell A, cells 600 to 800 to the secretion initiation stage of cell A2, and cells 5300 to 6700 to the mid-secretion stage of cell A1. Assume there are 30 functional combinations that predominate in the data. The output from this stage cannot yet be directly applied to subsequent analyses. Although the algorithm has successfully grouped cells according to function, the cell groups representing the 30 functions remain in an unordered arrangement within matrix M2. This stems from the inability at this point to discern whether the grouping basis is functional similarity, cell type proximity, or other underlying factors. In other words, Section 2.2.2 introduces a self-supervised learning module that integrates reinforcement learning principles. This module allows the model to automatically classify cells without predefined labels and to optimize clustering results through an iterative mechanism, ultimately producing clustered units composed of functionally similar cells. This accomplishes only the functional grouping of cells, without yet specifying the precise types and functional attributes of cells within each group.

The third step involves the sorting and hierarchical organization phase for functional groups. This step aggregates the 10,000 x 10,000 cell similarity matrix M2 into a 30 x 30 group similarity matrix M3. Due to significant differences in the numerical analysis resolution required for M2 and M3, a specialized processing module is needed to handle the unique structure of M3. Section 2.2.3 introduces the mask optimization module, which is used to reorder M3. This algorithm first sorts M3 according to the most dominant difference factor in the data. This dominant factor can be cell type, cell functional state, or other features, determined by the inherent difference characteristics of the data. In functional remodeling samples, the factor occupying the most dominant position in transcriptional differences is the functional differentiation type, so cells will be assigned to A, A1, and A2 according to functional differentiation type, thereby generating the sorted 30 x 30 matrix M4. At this point, the algorithm has provided cluster boundaries for each type, for example, groups 1 to 8 belong to A, groups 9 to 15 to A1, and groups 16 to 30 to A2. Since the sorting process is progressively based on the similarity of cells in each group, taking cell A as an example, the data within groups 1 to 8 have been sorted according to the second most dominant factor. Similarly, the dominant factor at any position can be obtained. Therefore, assuming group 1 is the starting end of the state corresponding to cell A, corresponding to the conventional metabolic functional state; then the terminal end is group 8, corresponding to cells fully prepared to initiate functional differentiation expression. This involves a resolution parameter, which allows users to independently determine the size of the matrix generated when aggregating M2 into M3, without affecting the algorithm’s cluster boundaries. For example, cell A can be subdivided into 8 progressive levels, or divided into any number of progressive levels according to needs.

The data presentation is illustrated in Figure 1. The functionality of the algorithm is depicted in Figure 5.

**Figure 1-1.**
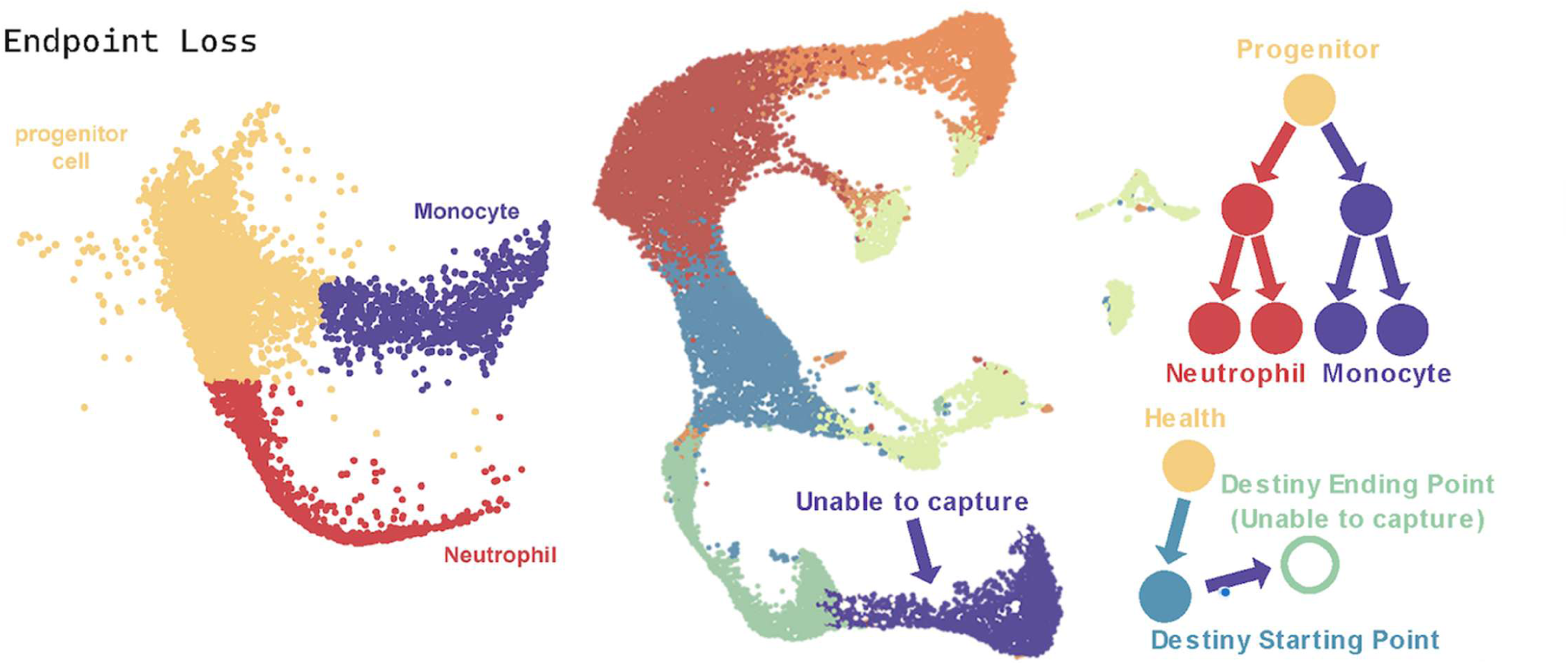
GMP cells can differentiate into monocytes and neutrophils, providing cells with clear, capturable endpoints in the context of developmental differentiation. In contrast, when defining boundaries based on cellular function, functional processes often occur in a overlapping and concurrent manner. Unlike developmental trajectories, functional states do not possess a discrete endpoint analogous to terminal differentiation.

**Figure 1-2.**
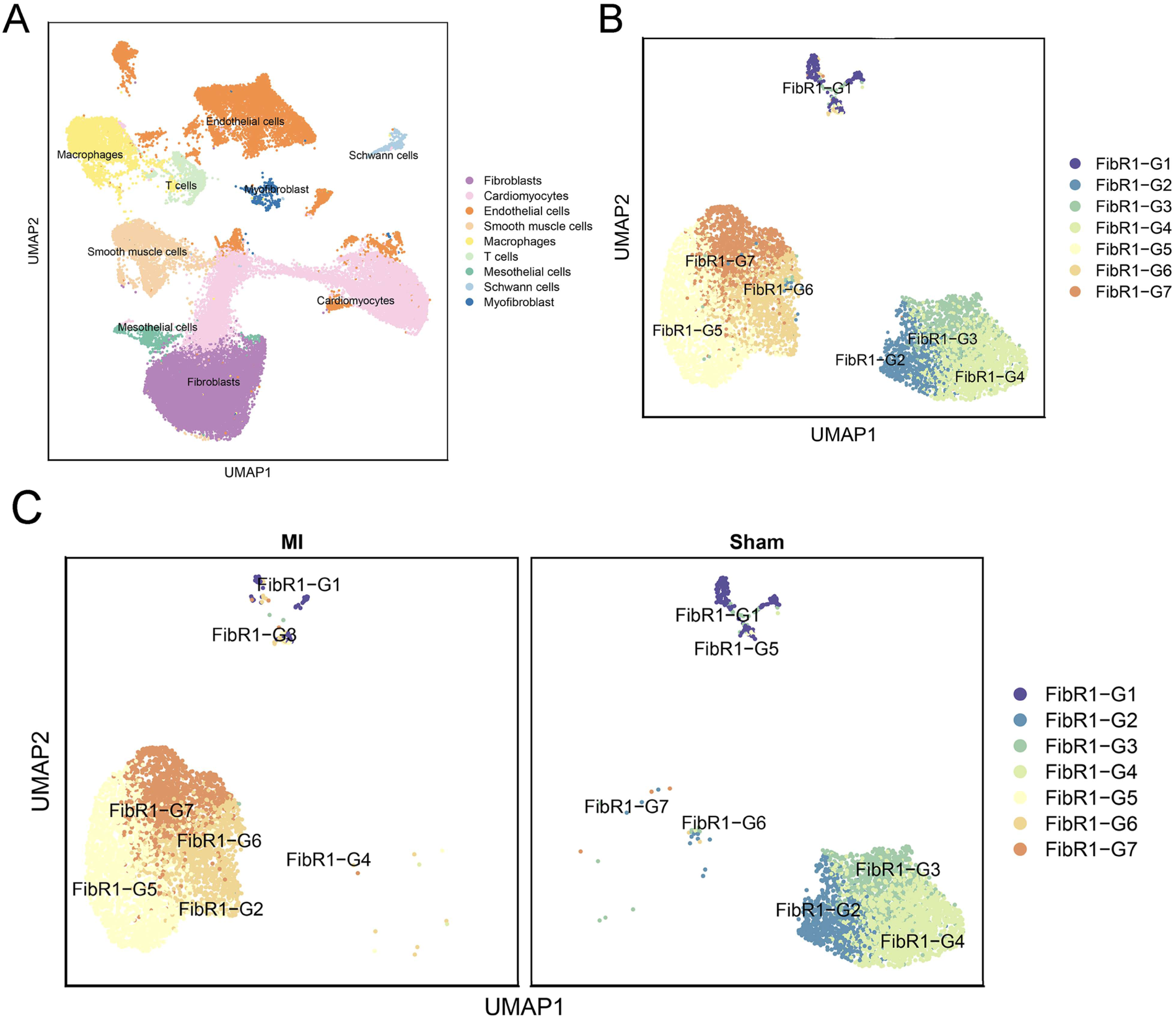
A: Cells in Background。 B: Fibroblasts and myofibroblasts were divided into 7 groups after TOGGLE analysis; C: Distribution of 7 cell types in MI and Sham groups after TOGGLE analysis; Before applying TOGGLE, the functional distinctions of cells lack color coding or group labeling (G1–G7). It is evident that, even when cells exhibit distinct functions, they do not necessarily display prominent trajectories.

**Figure 2.**
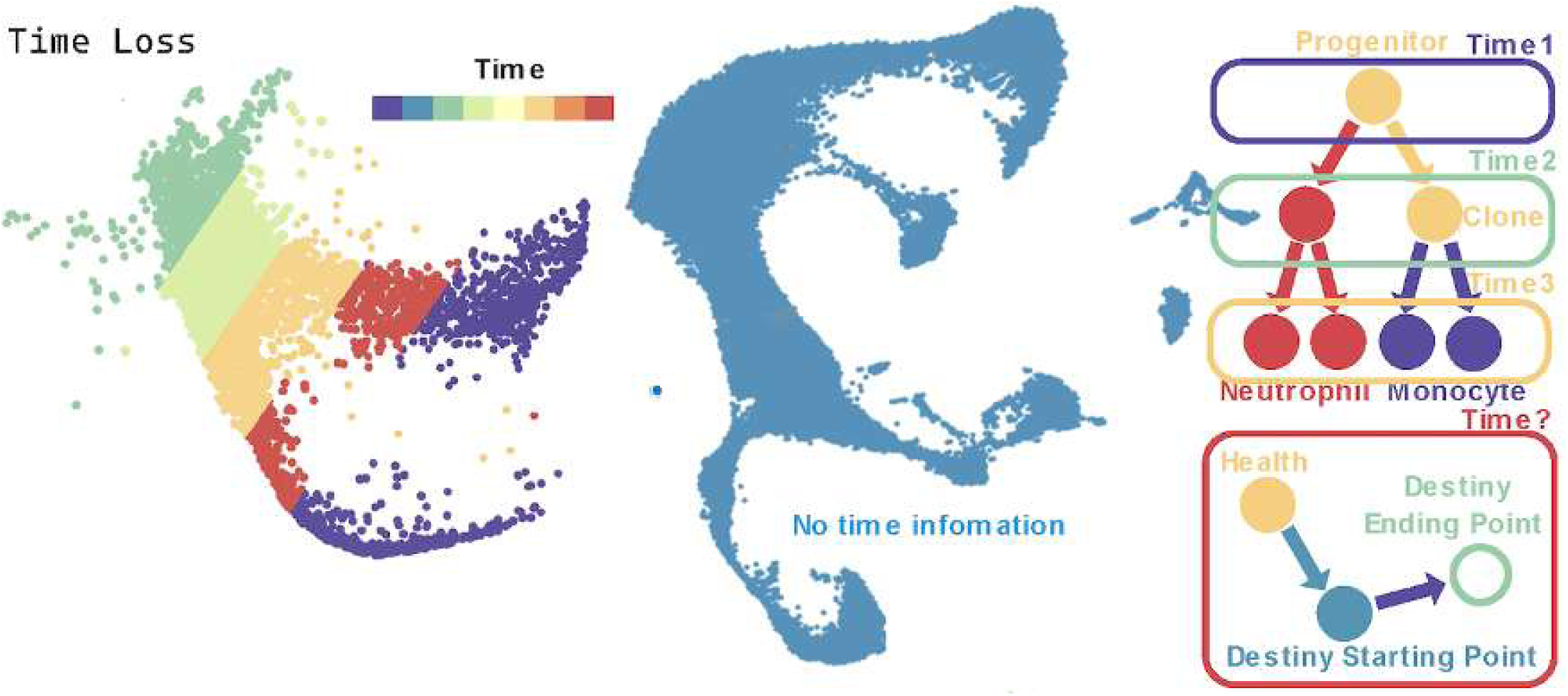
In developmental processes, cells exhibit an ideal single temporal order. In contrast, the execution of cellular functions occurs in an overlapping and concurrent manner, often involving parallel performance of multiple functions or transitions between functions. As a result, functional states do not possess a single, sequential temporal progression. This characteristic requires that our testing approach does not rely on time-related information.

**Figure 3.**
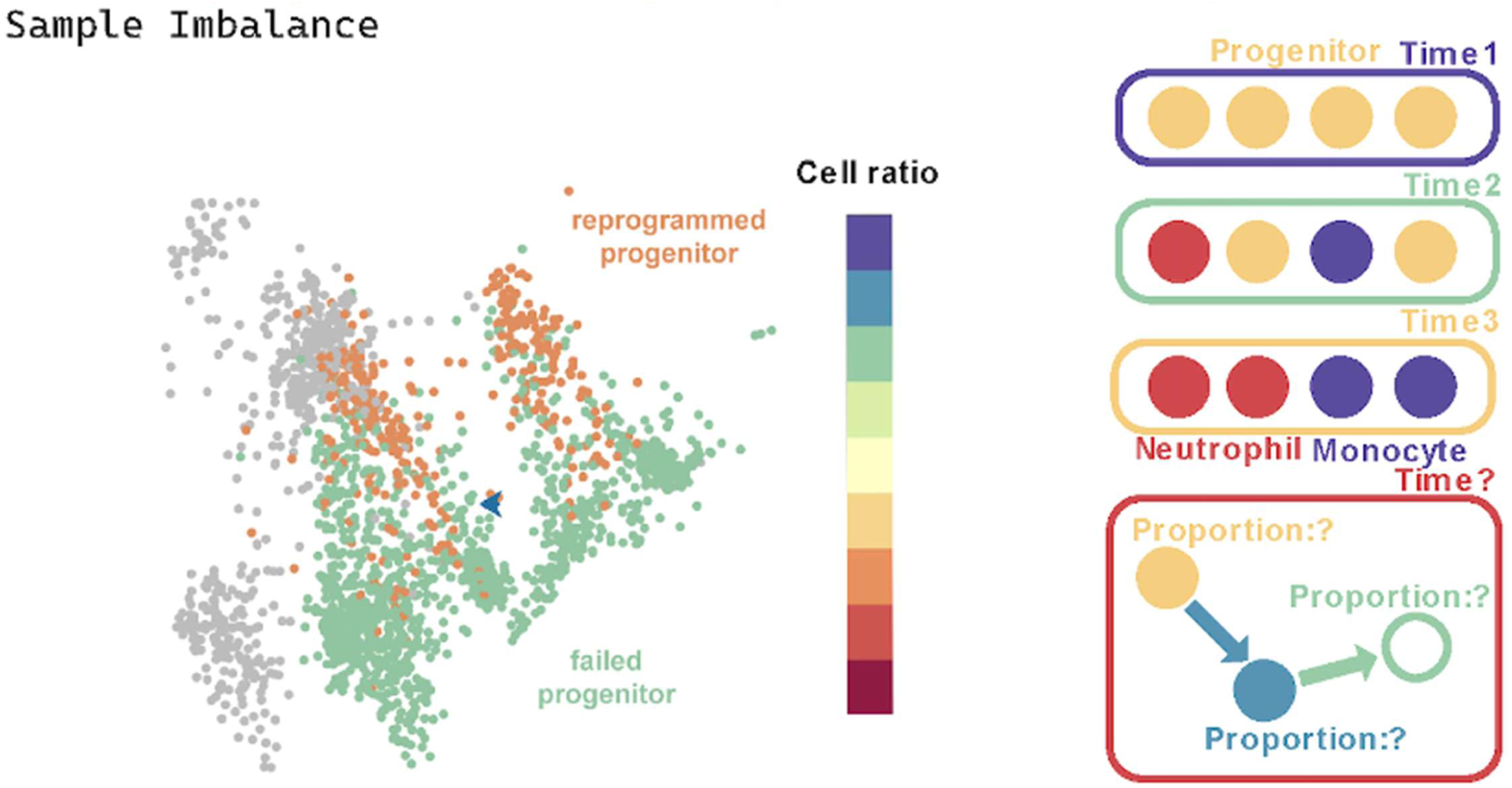
In developmental processes, cells can be tracked using lineage tracing techniques, allowing samples to be collected at appropriate time points to analyze optimal critical states of differentiation. In contrast, cells actively performing functions are often at or near terminal differentiated states. This requires that our testing approach does not rely on balanced sample information across different stages.

#### 2.2.1 Step 1: Preprocessing, Graph Construction and Asymmetric Similarity Computation

To construct the cell-cell similarity graph within the TOGGLE framework, we first calculate pairwise distances between all cells using Euclidean distance based on their gene expression profiles. We then convert these distances into similarity scores with a Gaussian kernel function that features an adaptive bandwidth. Here, the temperature parameter β = 0.1 controls the decay rate, assigning higher scores to nearby cells while still providing moderate weights to more distant ones. To balance noise reduction with the preservation of information, we apply row normalization to the weighted adjacency matrix, transforming it into a row-stochastic transition matrix. We also perform a single round of graph diffusion to introduce mild smoothing without altering the original local structural features. Finally, we sparsify the resulting similarity matrix by eliminating very small values, which reduces the computational load. All parameters—such as the number of neighbors set to the total number of cells, diffusion rounds equal to 1, β equal to 0.1, and sparsification threshold of 0.001—remain constant across all datasets and are not adjusted for specific datasets, ensuring the method’s consistency and reproducibility.

To construct the cell-cell similarity graph that serves as the foundation of the TOGGLE framework, we first build a k-nearest neighbor (kNN) graph based on the gene expression matrix. Let *X* ∈ ℝ^n×p^ denote the preprocessed gene expression matrix, which includes n cells and p genes (following standardization, logarithmic transformation, and selection of highly variable genes, as described previously). We employ Euclidean distance to compute the distance matrix between cells:

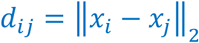

Based on this distance matrix, we constructed an undirected k-nearest neighbor graph, where V represents the set of n cells. The value of k is set to the maximum possible value in the dataset, namely n-1 (that is, considering all other cells as potential neighbors for each cell), thereby constructing an approximately fully connected initial adjacency relationship (with actual edge weights still determined by the subsequent kernel function). We did not enforce the requirement that neighbors be mutual in the k-nearest neighbor sense.

The edge weights are defined using a Gaussian kernel function, with a temperature parameter introduced for regulation:

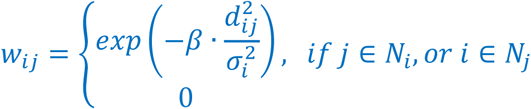

Where *α*_*i*_ denotes the distance from cell *i* to its A-th nearest neighbor (serving as an adaptive bandwidth), and *β* = 0.1 is a hyperparameter that controls the decay speed of the kernel function (a smaller value results in a smoother kernel function, with slower weight decay for distant cells).

In the present study, we employ a minimally intensive diffusion strategy, conducting only a single diffusion iteration (round_of_smooth = 1), to preserve the local features of the original k-nearest neighbor structure while incorporating a modest amount of long-range information through exponentiation of the transition matrix:

1. Normalize the adjacency matrix into a row-stochastic transition matrix *P*:

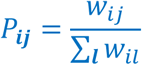
2. Perform a single-step diffusion:

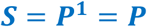 Since the number of diffusion steps is strictly limited to 1, and the number of neighbors is set to the maximum possible value (*k* = *n*), the resulting similarity matrix largely preserves the global information of the original distance structure, while achieving a balance between smoothing and noise suppression through the parameter and mild diffusion.
3. To reduce storage requirements and computational costs, perform sparsification on the resulting similarity matrix *S*.

##### Parameter Description

In all experiments conducted in this study, the key parameters in the graph construction and asymmetric similarity calculation processes remain fixed and unchanged to ensure the method’s consistency and reproducibility. Specifically, the number of neighbors is uniformly set to the total number of cells in the dataset (*neighbor*_*N* = *n*, that is, taking the maximum number of cells available), the diffusion strength is fixed to a single iteration (*round*_*of*_*smooth* = 1), the Gaussian kernel temperature parameter is always *beta* = 0.1, and the sparsification threshold is uniformly adopted as *truncation*_*threshold* = 0.001. These parameters were not individually tuned for different datasets or experiments, but were applied completely consistently to all analysis samples (including Hematopoiesis, Reprogramming, and any additional validation datasets).

The core processing principle of this method is to employ the most primitive form of diffusion: by constructing a nearly fully connected graph (k = n) and applying a single-step diffusion (t = 1), under the control of a mild temperature parameter (β = 0.1), the transition matrix P is made to approximate a smoothed version of the original similarity structure. This approach uses the inherent geometric smoothness of the data itself as a benchmark, thereby fully presenting the global overview of all potential relationships between cells, while ensuring low sensitivity to parameters. During our testing, we found that, excluding extreme parameter cases, the calculations in subsequent sections are almost insensitive to parameters, unless the user intentionally employs extreme parameter situations.

- **Diffusion steps round_of_smooth = 1:** Employing a single-step diffusion—that is, directly using the row-normalized transition matrix P as the similarity matrix—avoids the risk of excessive smoothing that arises from multi-step exponentiation operations. In single-cell RNA sequencing data, literature indicates that multi-step diffusion, such as when t exceeds 3 to 5, frequently results in the loss of biological signals. In contrast, single-step or very mild diffusion effectively preserves the balance between maintaining local structures and suppressing noise, and shows relative insensitivity to the selection of t, particularly when k is large.
- **Number of neighbors neighbor_N = n (taking the maximum number of cells):** Setting k to fully connected (or nearly fully connected) makes the initial graph approach a complete graph structure, avoiding the sensitive impact of k value selection on local connectivity and global topology in traditional k-nearest neighbor methods. In diffusion maps or similar methods, when k is sufficiently large (approaching fully connected), the steady-state distribution of the transition matrix and the diffusion distance exhibit strong robustness to further increases in k, because information propagation has covered almost all paths between cells.
- **Gaussian kernel temperature parameter β = 0.1:** The β value controls the decay rate of the kernel function. A smaller β, such as 0.1, renders the kernel function smoother, enabling distant cells to retain certain weights. This further reduces reliance on precise bandwidth or k values in fully connected configurations. Similar temperature parameters frequently demonstrate stability across a broad range in graph diffusion and affinity matrix construction, particularly when integrated with adaptive bandwidth and row normalization.

#### 2.2.2 Step 2: Identify functionally similar cell clusters using a self-supervised learning algorithm

We propose a self-supervised algorithm—a type of unsupervised algorithm in which the method generates its own conclusions for self-reference—designed for unsupervised grouping of samples based on a precomputed matrix of similarities between cells. This approach falls within the broader category of unsupervised learning. Specifically, it generates internal structural signals through repeated transformations of the similarity matrix, which in turn guide the recursive division and ordering of samples.

The algorithm does not depend on external labels. Instead, it uses the matrix’s own correlation patterns as substitute guidance signals to identify consistent subgroups. Each iteration refines the matrix representation and bases subsequent recursive steps on prior outcomes, thereby establishing a layered, self-improving process of discovery. Let *S* ∈ ℝ^n×n^ represent the input similarity matrix, where *n* denotes the number of samples (such as cells), and *S*_*ij*_ quantifies the pairwise similarity between sample *i* and *j*. The algorithm consists of two primary phases: (1) recursive binary sorting to produce ordering vectors and division points, and (2) removal of duplicates and establishment of boundaries to define grouped blocks.

(1) Recursive binary sorting to generate permutation vectors and split points:

The core process is a recursive function that iteratively transforms *S* to extract dominant structural patterns. Based on these patterns, it sorts the samples and bipartitions the set until stopping criteria are met. Define the transformation function ∅: ℝ^n×n^ → ℝ^n×n^ as the operator that computes the Pearson correlation coefficient matrix for the row vectors of the input matrix. Specifically, for the input matrix M, first center each row by subtracting the row mean. Then, compute the Gram matrix (inner product matrix) of the centered matrix, and standardize it using diagonal normalization factors—the reciprocals of the square roots of the Gram matrix’s diagonal elements—to obtain the correlation coefficient matrix. For a given submatrix M ⊆ *S* of size *m* × *m*, initially M = S, *m* = *n*, the process is as follows:

1. Iterative transformation: For the iteration count *I*—after which performance stabilizes completely once *I* is sufficiently large, thus setting it adequately high when computational resources permit—repeatedly apply ∅.

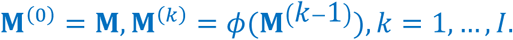 This amplifies coherent patterns within the similarity structure, serving as self-supervised signals by emphasizing the inherent consistent correlations within the data itself.
2. Thresholding and feature extraction: Binarize the transformed matrix to preserve sign information.

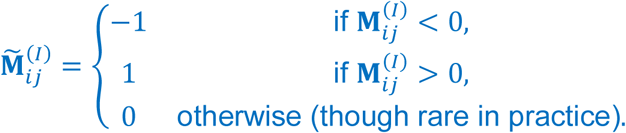 Extract the first two columns as approximate dominant features (serving as low-rank proxies for the matrix’s principal directions).

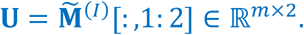

This approach can be extended to acquiring the top *k*singular vectors through sparse singular value decomposition to enhance efficiency, though when *m* is small, using columns simplifies the computation.
3. Sorting and splitting: Sort the samples based on the first feature dimension, where the sorting function returns ascending indices. Identify the split point as the transition in the sorted first dimension from negative to non-negative values.

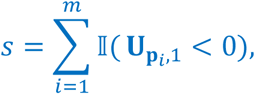 Here, 𝕀 represents the indicator function. This split point delineates two subgroups based on self-supervised sign partitioning.
4. Stopping criteria and recursion: If the subgroup size falls below the minimum length threshold *L* (for example, *m* < *L*), or if the split is unbalanced (for example, *s* < *δ* or *m* − *s* < *δ*, where *δ* represents the minimum split size), terminate the recursion and return P = [1, …, *m*] and *s* = 0. Otherwise, recursively invoke the process on the submatrix of the left subgroup, M[P_1:s_, P_1:s_] to obtain the permutation P_l_ and split point *s*_l_ ; similarly, apply the recursion to the submatrix of the right subgroup, M[P_s+1:m_, P_s+1:m_], yielding P_R_ and *s*_R_, where Δ denotes an additional asymmetry tolerance (typically set equal to *δ*). Combine the full permutation and collect the split points.

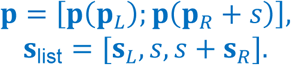

Here, each recursion is conditioned on the transformation and sorting output from the previous layer, creating a hierarchical self-supervised chain in which prior structural discoveries inform subsequent partitions. The recursion depth is implicitly constrained by the stopping criteria, ensuring termination on average at *o*(log *n*) layers. The overall permutation P reorders the samples to position similar samples consecutively, while *s*_list_ accumulates the hierarchical split points.

(2) Deduplication and boundary construction to define clustering blocks:

After recursion, deduplicate the split list to remove redundant boundaries while preserving order, yielding unique elements in the sequence of their first appearances. Construct simplified boundaries by prepending the starting index and appending the end.

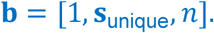

These boundaries b define *K* =∣ b ∣ −1 continuous blocks in the reordered matrix *S*[P, P], which are interpreted as unsupervised clusters. Each block *i* spans samples from b_*i*_ to b_i+1_− 1.

Parameters:

- *I*: Number of transformation iterations (for example, 5–20) to stabilize the self-supervised signal. Once *I* is sufficiently large, performance tends to become completely unchanged, so make it sufficiently large when computational power is adequate.
- *L*: The minimum subgroup size for recursion (for example, 100) to prevent overfitting to noise.
- *δ*, Δ: The minimum split size (for example, 5) to ensure balance. The self-supervised nature of this algorithm originates from using iterative matrix transformations as internal pretext tasks, in which each recursion is guided from the refined representation of the previous step. This promotes the emergence of clustering structures without requiring external supervision. The computational complexity is *o*(*In*^2^ + *n*log *n*) per recursive layer, dominated by transformations of the dense *S*, but if needed, it can be extended through sparse approximations.

The pseudocode for this subsection is as follows:

##### Correlation transformation function 𝝓 (correlation_phi)

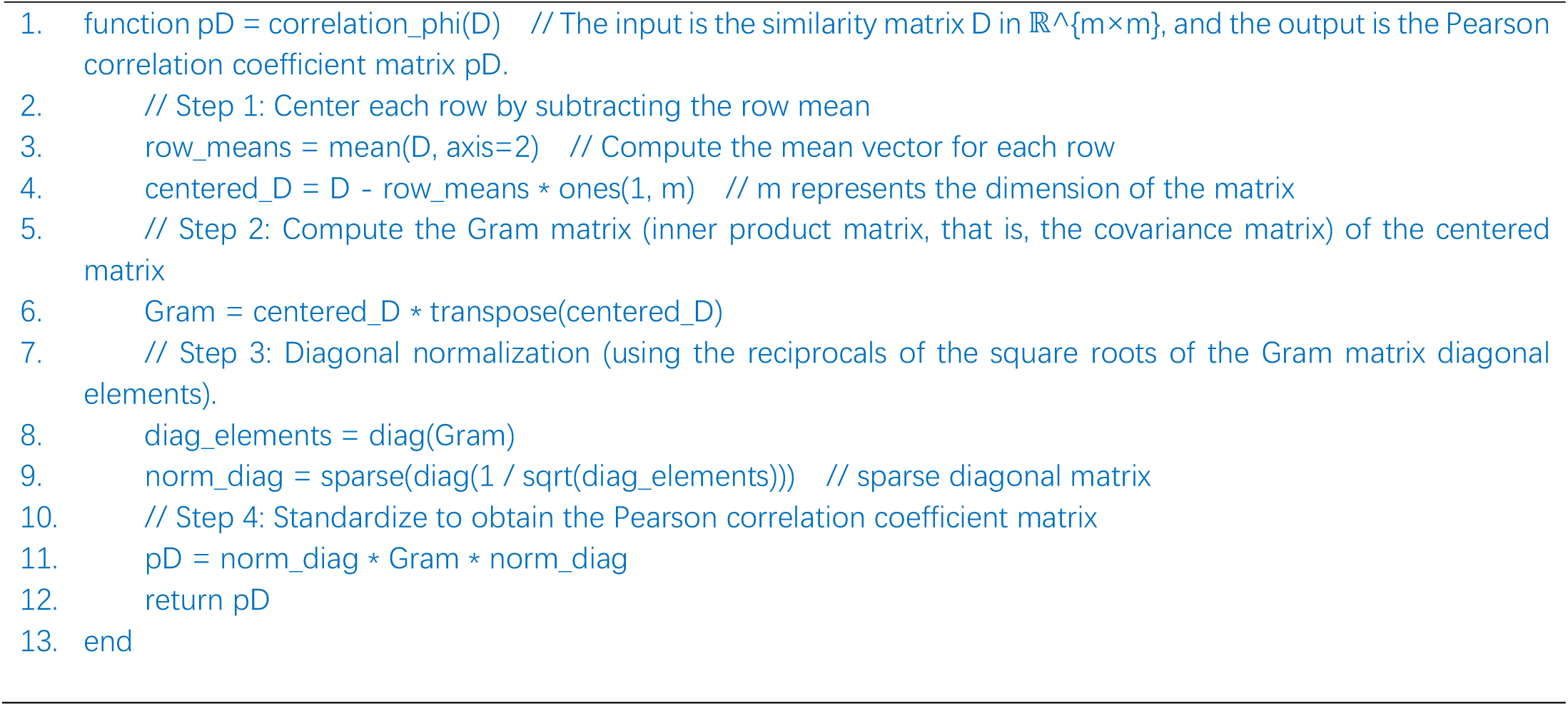

##### Recursive binary sorting main function (binary_corr_sorting)

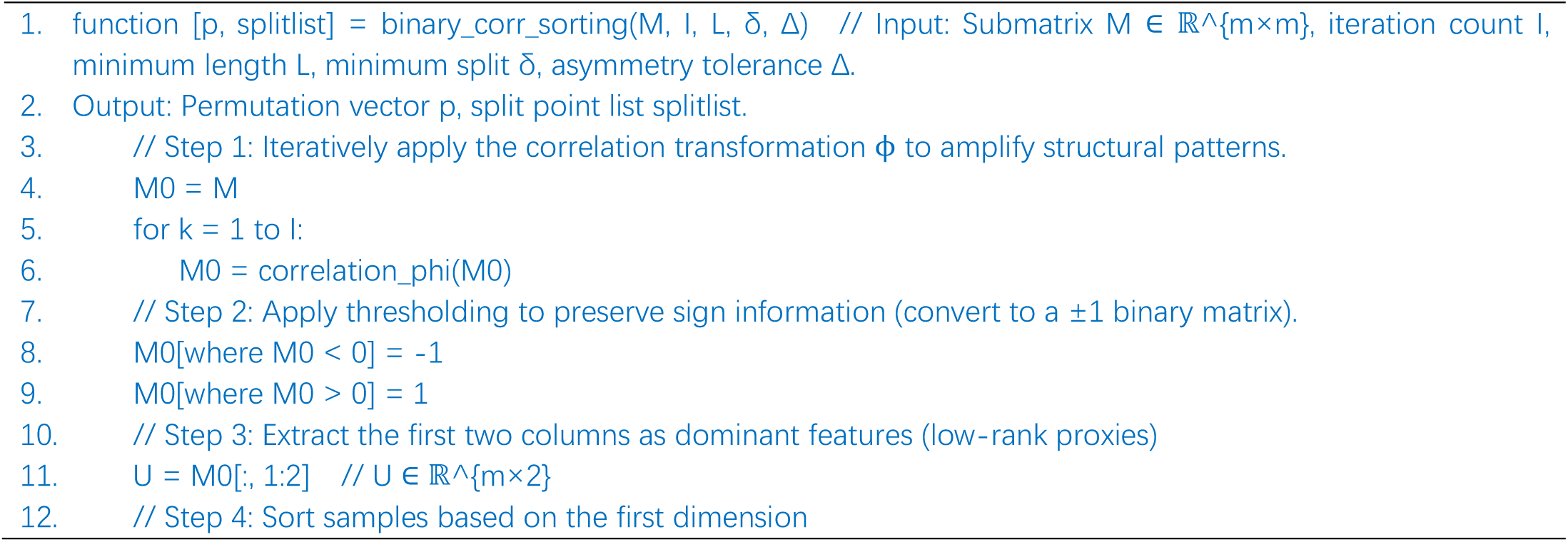
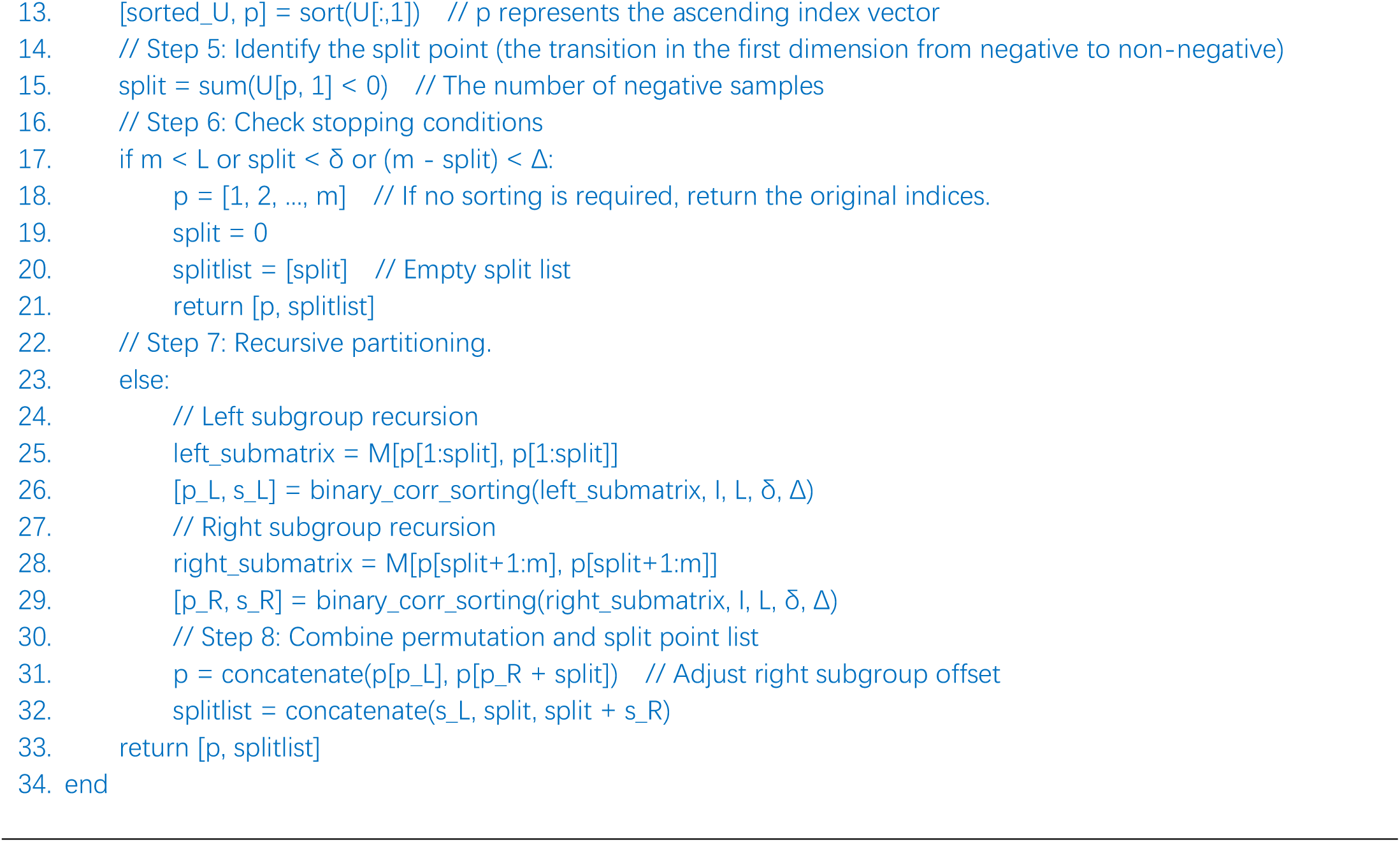

#### 2.2.3 Step 2+3: Group functionally identical cell clusters together

The correlation-based clustering from Step 2 generates boundary markers that separate different functional regions, making it straightforward to determine each cell’s relative position with respect to these markers. By selecting cells located between adjacent markers, we can identify functionally similar cell groups. However, cellular functional relationships are often nonlinear, meaning that different cell types with distinct biological functions may still exhibit similar correlation patterns in the similarity space. This poses a fundamental limitation: correlation-based grouping alone is insufficient to determine final cell type classifications, as it may merge functionally distinct cell populations that happen to share similar correlation signatures. To address this challenge, we developed a Deep Genetic Network that adapts the masking framework from BERT to an unsupervised setting. This approach captures global similarity relationships between individual cell clusters and all other clusters, enabling more precise functional classification by considering contextual information beyond local correlations. The final functional assignment of each cluster is determined through this integrated framework (Figure 4).

**Figure 4.**
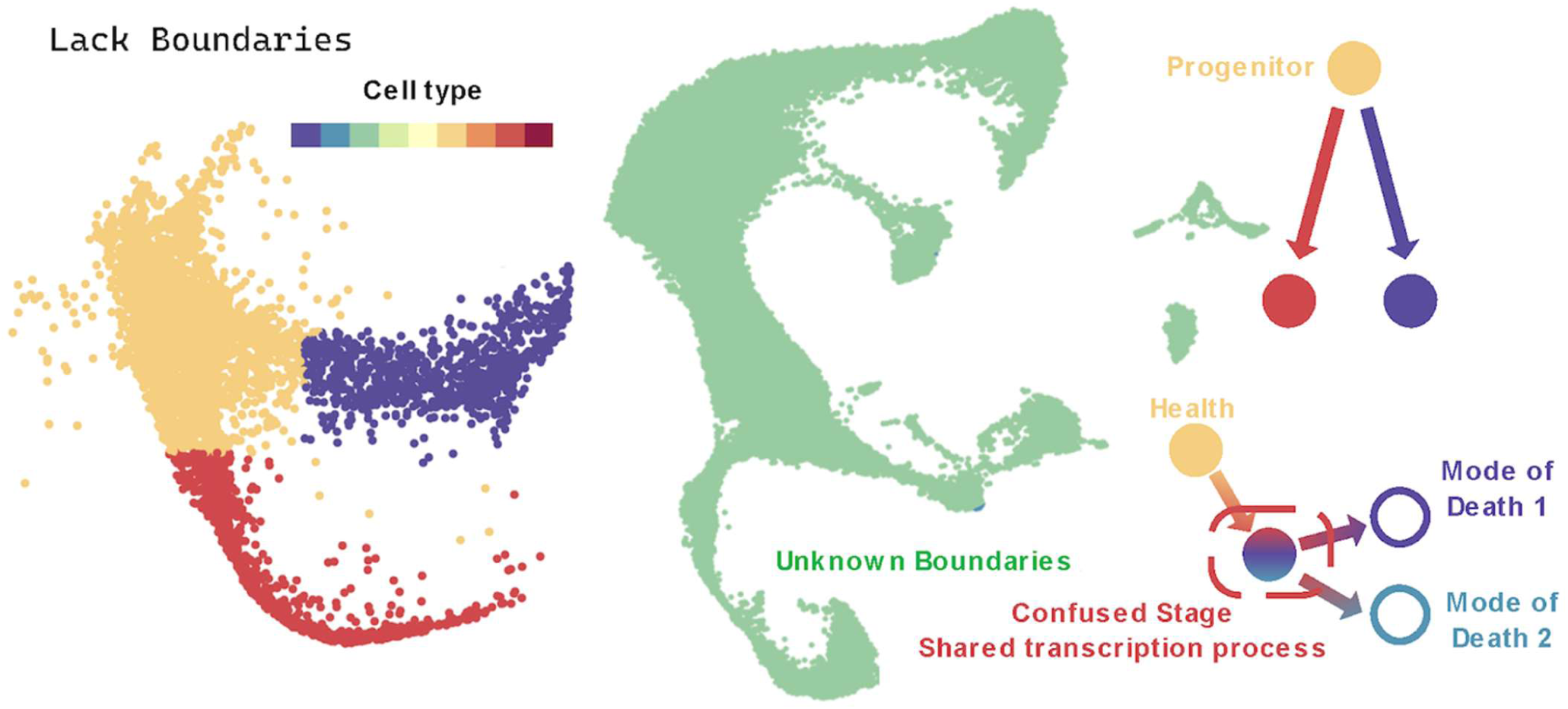
In developmental processes, cells exhibit clear and distinct cell type identities, such as GMP cells, monocytes, and neutrophils. In contrast, under functional tracing techniques, cells do not display prominent differences in cell type. At the same time, it is difficult to manually determine which functional differences should serve as marker features for separation. This requires that the algorithm be capable of distinguishing cells based on the maximum functional differences it identifies.

To enforce semantic coherence in the reordered matrix (i.e., grouping cells by shared annotations such as cell types), we employ a genetic algorithm (GA) to optimize the order of label blocks while preserving intra-block smoothness. Let L ∈ ℝ^K_1_×1^ be the aggregated labels from Step 1, and *S* ∈ ℝ^K_1_×K1^ the micro-cluster summary matrix, where

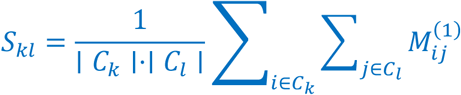

is the mean similarity between micro-clusters *C*_k_and *C*_l_. The unique labels are 𝒰 = {*u*_1_, …, *u*_K_}with *K* ≤ *K*_1_, and each *u*_*m*_corresponds to a set of micro-clusters ℐ_*m*_ = {*i*: *L*_*i*_ = *u*_*m*_}.

The GA operates on the permutation space Π_K_of label blocks, where a candidate solution *π* ∈ Π_K_encodes the sequence of blocks [*π*(1), …, *π*(*K*)]. For each block *π*(*m*), the internal micro-cluster order is fixed via average-linkage hierarchical clustering on the pairwise distance submatrix 𝐃_ℐm,ℐm_, where 𝐷_ij_ = 1 − 𝜌(*S*_*i*_, *S*_j_)and 𝜌(⋅,⋅)is the Pearson correlation (falling back to Euclidean distance ∥ *S*_*i*_ − *S*_j_ ∥_2_ if rows are constant). The linkage tree 𝐙_*m*_ is computed as 𝐙_*m*_ = linkage(squareform(𝐃_ℐm,ℐm_),’ *a*𝑣*erage* ‘), and the leaf order 𝜆_*m*_ = optimalleaforder(𝐙_*m*_,squareform(𝐃_ℐm,ℐm_))yields the intra-block permutation 𝐨_*m*_ = ℐ_*m*_[𝜆_*m*_]. The full order for *π*is then 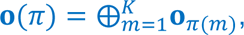 the concatenation of intra-block orders.

The fitness function *f*(*π*): Π_K_ → ℝ_+_balances smoothness and label integrity:

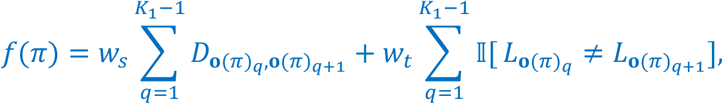

where 𝑤_s_ = 1 weights adjacent distances (normalized by the median off-diagonal 𝐷), 𝑤_t_ = 50 ⋅ median(𝐃_off-diag_) penalizes label transitions (ideally minimized to *K* − 1), and 𝕀[⋅] is the indicator function. The scale of 𝑤_t_ensures that splitting label blocks is prohibitively costly relative to smoothness gains.

For small *K* ≤ 8, we exhaustively evaluate all *K*!permutations in Π_K_to find *π*^∗^ = arg min _n∈TT_ *f*(*π*). For larger *K*, a standard GA is run for *c* = 5independent restarts, each with population size *N*_p_ = 60, offspring ratio *n*_c_/*N*_p_= 1(yielding *n*_c_ = 60children per generation), and *I*_max_ = 200generations. The population is initialized with random permutations of {1, …, *K*}. Elitism retains the top *e* = 0.2*N*_p_ individuals. Offspring are generated via order crossover (OX): for parents *π*^(1)^, *π*^(2)^, select cutpoints *i* < *j*uniformly, copy *π*^(1)^[*i*: *j*]to the child, and fill the remainder with elements from *π*^(2)^in cyclic order while preserving relative positioning. Swap mutation flips two random positions with probability 𝜇 = 0.25. The next generation is selected via fitness-proportional tournament (size 2). The global optimum is the best *π*^∗^across restarts. The final outputs are the reordered label vector L^A^ = L[𝐨(*π*^∗^)]and matrix *Ŝ* = *S*[𝐨(*π*^∗^), 𝐨(*π*^∗^)].

In each GA generation, the evaluation of a permutation *π*conceptually treats the positioning of each label block as a “random walk” candidate: the fitness aggregates contributions from all blocks, where the placement of block *m*is scored relative to its neighbors based on the precomputed inter-block distances in 𝐃and the transition penalties enforced by 𝑤_t_. This ensures that block positions are determined holistically by the global data configuration (i.e., similarities to other blocks), rather than greedily.

Boundary detection is performed on the reordered micro-cluster summary matrix *Ŝ* ∈ ℝ^K_1_×K1^ (or a progenitor-filtered submatrix for lineage tasks), employing an autoencoder-inspired ridge detection method. This approach generates binarized correlation matrices through multi-resolution thresholding iterations and extracts boundary strength signals therefrom. Specifically, the combination of the encoder ℰand decoder 𝒟produces a weighting matrix 𝐖for computing potential boundary locations.

The encoder ℰ(*Ŝ*; *u*, *l*, *r*, 𝜌)first transforms the input matrix *Ŝ*into a latent representation 𝐄 ∈ ℝ^K_1_×K1^, where *u* and *l* are the upper and lower thresholds, respectively, *r* = 10is the resolution, and 𝜌 = 3is the correlation rounding parameter. The algorithm initializes 𝐄 = 𝟎_K1×K1_ and accumulates binarized correlation matrices 𝐂^(m)^over *r* + 1iterations:

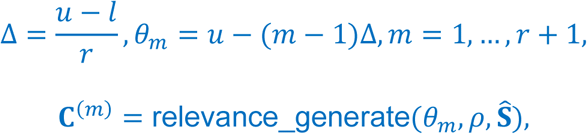

where relevance_generate(⋅) produces a binarized matrix based on threshold 𝜃_*m*_ and rounding 𝜌(specific implementation depends on correlation computation logic, not explicitly defined in the core pipeline). After accumulation, 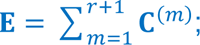 to mitigate diagonal noise, all elements of 𝐄are sorted in descending order, the median of unique values *ẽ* = median(unique(sort(𝐄(:),’descend’))) is extracted, and the diagonal is replaced: 𝐸_ii_ ← *ẽ*, ∀*i*.

The decoder 𝒟(𝐄) extracts a boundary points matrix 𝐁 ∈ {0,1}^K_1_×K1^ from 𝐄 via nested loops detecting threshold crossings in row and column directions (from > 0to = 0or vice versa):

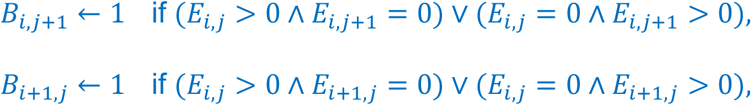

∀*i*, *j* = 1, …, *K*_1_ − 1. Subsequently, row and column boundary strength summaries are computed: 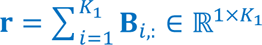 (row sums per column) and 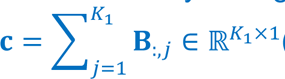(column sums per row). The initial reconstruction matrix 𝐃^(O)^ ∈ ℝ^K_1_×K1^ is populated via broadcasting: each row 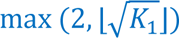 and each column 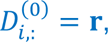 To capture the length of continuous boundary segments, the decoder further generates two auxiliary matrices 𝐑_1_, 𝐑_2_ ∈ ℝ^K_1_×K1^ for horizontal and vertical directions, respectively. For the horizontal direction (𝐑_1_), for each row *i* = 1, …, *K*_1_ − 1, a counter 𝜅 = 0is initialized, and along columns *j* = 1, …, *K*_1_ − 1, 𝜅 ← 𝜅 + 1is incremented; at $0 \to 1$ or $1 \to 0$ transitions, 𝑅_1,i,j+1_ = 𝜅is recorded and 𝜅 = 0is reset. The vertical direction (𝐑_2_) proceeds analogously but loops over rows for column transitions. The final weighting matrix is:

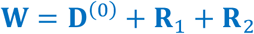

𝐑_1_and 𝐑_2_are auxiliary matrices in the decoder that quantify the persistence of boundary segments by measuring the lengths of consecutive runs of identical values (0s or 1s) in the binary boundary points matrix 𝐁. Specifically, 𝐑_1_operates in the horizontal direction: for each row *i*, it initializes a counter 𝜅 = 0and increments it along columns *j*, assigning the counter value to 𝑅_1,i,j+1_at each transition (from 0 to 1 or 1 to 0 between 𝐁_i,j_and 𝐁_i,j+1_) before resetting 𝜅 = 0, thereby encoding the length of the preceding uniform segment ending at *j* + 1. Analogously, 𝐑_2_processes the vertical direction: for each column *j*, it increments 𝜅along rows *i*and records 𝑅_2,i+1,j_ = 𝜅at transitions between 𝐁_i,j_and 𝐁_i+1,j_, capturing vertical run lengths. These matrices enhance the weighting by prioritizing longer, more coherent ridges indicative of true cluster boundaries over isolated noise points. The boundary score vector 𝜷 ∈ ℝ^K1-1^is defined as the potential ridge strength:

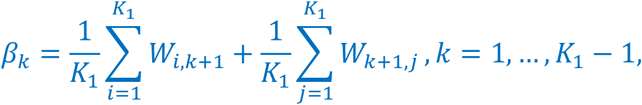

with the diagonal excluded (pre-set to NaN). For a target number of clusters *K*^∗^, the top *K*^∗^ − 1peaks in {*β*_k_}(above the empirical threshold 𝜇_p_ + 0.5*α*_p_, where 𝜇_p_, *α*_p_ are the mean and standard deviation of 𝜷; minimum block size 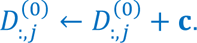 ensures spacing) yield boundary positions b = [*b*_1_ < ⋯ < *b*_K_∗_-1_]. If a ground-truth label sequence 𝚲 ∈ 𝒰^K1^ is provided, a snapping mechanism is enabled: the transition set 𝒯 = {*k*: Λ_k_ ≠ Λ_k+1_, *k* = 1, …, *K*_1_ − 1}is identified, each *b*_q_is snapped to the nearest *t* ∈ 𝒯(based on min ∣ *b*_q_ − *t* ∣), and the selection is supplemented or pruned to exactly *K*^∗^ − 1positions ranked by *β*_k_to align with label block interfaces.

##### Genetic Encoder: Label Block Optimization for Matrix Reordering and Boundary Detection

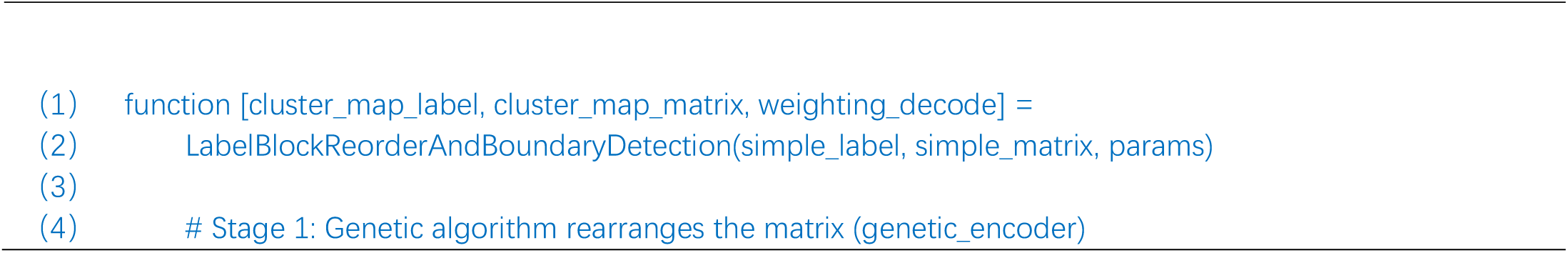
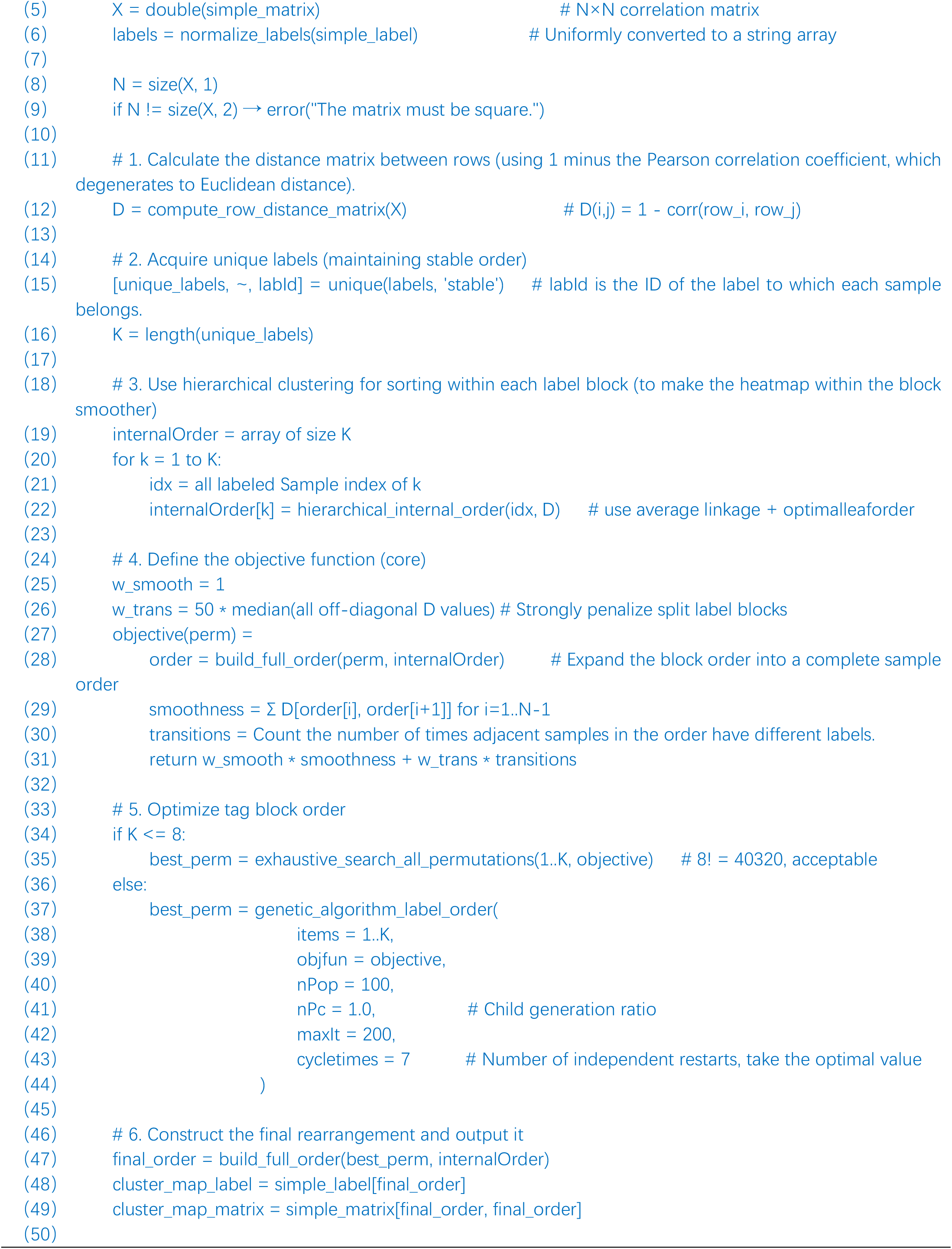
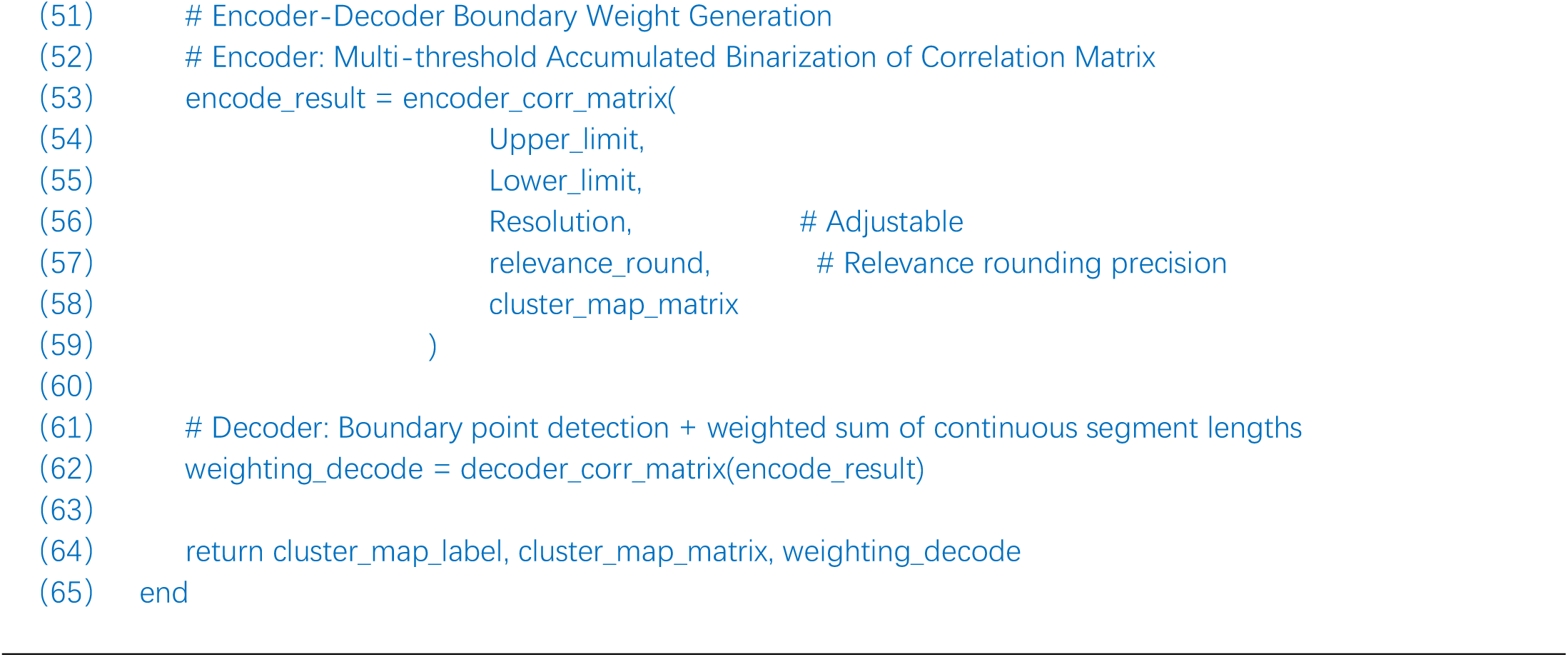

### 2.3 Graph Diffusion Functional Map

We used UMAP to visualize the functional grouping of RNA. While UMAP effectively captures overall expression trends, its dimensionality reduction process inevitably results in information loss, leading to the mixing of RNA species with distinct functional roles. This mixing effect hinders accurate discrimination of functional states and limits the depth of target identification and functional interpretation. To address this limitation, we propose a novel approach: the Graph Diffusion Functional Map. As described in Section 2.2, we replace the original cell-based graph diffusion matrix 𝑀_ij_ with a matrix derived from RNA–RNA correlations and apply the full computational framework detailed in the Methods section to construct the functional map. This strategy significantly reduces functional mixing while preserving the intrinsic correlation structure among RNAs, enabling clearer delineation of RNA functional groupings. Unlike conventional dimensionality reduction techniques, this graph diffusion-based method retains high-dimensional information and improves sensitivity to subtle functional differences. Our results demonstrate that the Graph Diffusion Functional Map not only recapitulates known functionally enriched clusters but also reveals previously undetected, highly expressed RNA groups overlooked by UMAP. These newly identified targets provide novel insights into single-cell transcriptomic data and expand opportunities for understanding disease mechanisms and identifying therapeutic targets.

### 2.4 Dataset & Protocols

#### 2.4.1 TEST 1: Performance Testing of TOGGLE Using Developmental Cells

To evaluate the accuracy of TOGGLE, it was first applied to several publicly available scRNA-seq datasets with clearly annotated ground-truth outcomes.

**(1) GSE140802**^30^: Mouse bone marrow dataset, a subset of progenitor cells differentiates into monocytes—immune cells specialized in pathogen engulfment—while others give rise to neutrophils, another distinct immune lineage. These are referred to as monocyte progenitors and neutrophil progenitors, respectively.

**(2) GSE99915**^31^: Reprogram mouse dermal cells into endoderm-like progenitors. A fraction of cells successfully acquired the target identity (reprogrammed progenitor cells), whereas others failed to transition (failed progenitor cells).

Since the final cellular fates are explicitly annotated in both datasets, they provide robust benchmarks for evaluating classification and fate-prediction performance.

#### 2.4.2 TEST 2: Distinguish the stages of ferroptosis by TOGGLE and perform biological validation

TOGGLE was then applied to a scRNA-seq dataset without cell state annotation.

**GSE232429**^1^: A dataset of neurons after ischemic stroke. It was used to identify cells from early, intermediate, and late stages of programmed cell death.

To evaluate the reliability of the prediction results, we observed significant alterations in genes involved in apoptosis and ferroptosis (two forms of programmed cell death) in the relevant cell populations, which can accelerate programmed cell death. These transcriptomic signatures were further corroborated by animal experiments, in which consistent protein expression changes were detected in the ischemic brain tissue of rats. Details of scRNA-seq data processing and experimental methods are shown in the Supplementary Appendix 6 – Animal Processing.

#### 2.4.3 TEST 3: Identifying Epigenetic Metabolic Regulation in NSCs Using Novel Functional Mapping of TOGGLE

The functional biology discovery capabilities of TOGGLE were further assessed by using the following dataset.

**GSE209656**^2,32^: Identified quiescent neural stem cells (qNSCs) embedded within a broader population of glial cells—a particularly challenging task due to the near-identical transcriptomic profiles shared by qNSCs and astrocytes.

By leveraging the Graph Diffusion Functional Map, we clustered RNAs based on functional similarity and uncovered epigenetic signatures associated with energy metabolism regulation.

#### 2.4.4 TEST 4: Verifying the Graph Diffusion Functional Map of TOGGLE with other datasets

The following datasets were used to further validate the ability of TOGGLE to make Graph Diffusion Functional Maps. These results are detailed in the Supplementary Appendix 1(**RNA Pathway Alterations in Air Pollution**) & Supplementary Appendix 2 (**Cellular functional mapping**).

**(1) GSE183614**^33^: it was used to detect RNA pathway perturbations.

**(2) GSE253768**: it was used to analyze intercellular communication.

## 3. Result

### 3.1 Test 1: Performance Testing Using Developmental Cells

To systematically evaluate TOGGLE’s performance, we benchmarked it against four widely used computational methods for lineage inference: Cospar, Super OT, WOT, Deep-Lineage, and GAN-Based OT. As an unsupervised algorithm, TOGGLE does not require prior training or labeled data, distinguishing it from supervised approaches. We assessed performance across two experimental datasets: mouse bone marrow hematopoietic progenitor cells and lineage-reprogrammed cells, evaluating accuracy both for progenitor cells alone and for combined progenitor-descendant cell populations.

The precision for class N is given by the formula:

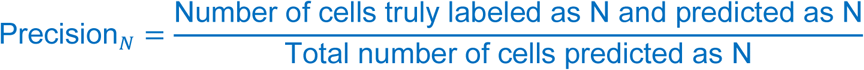

This measures the proportion of correctly predicted N cells among all predicted N cells.

**Table 1.**
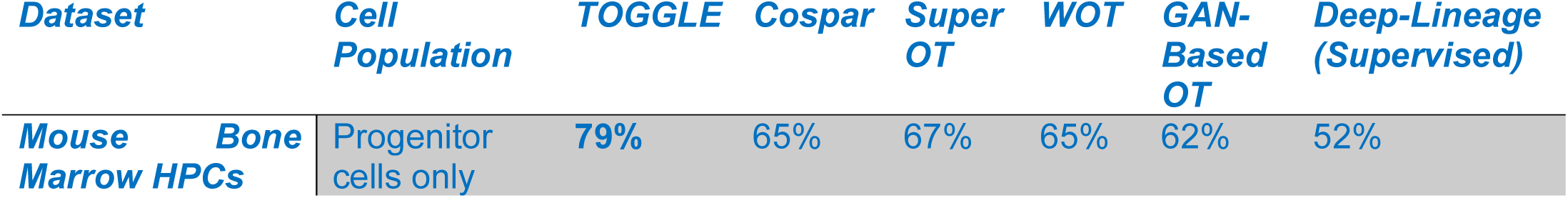
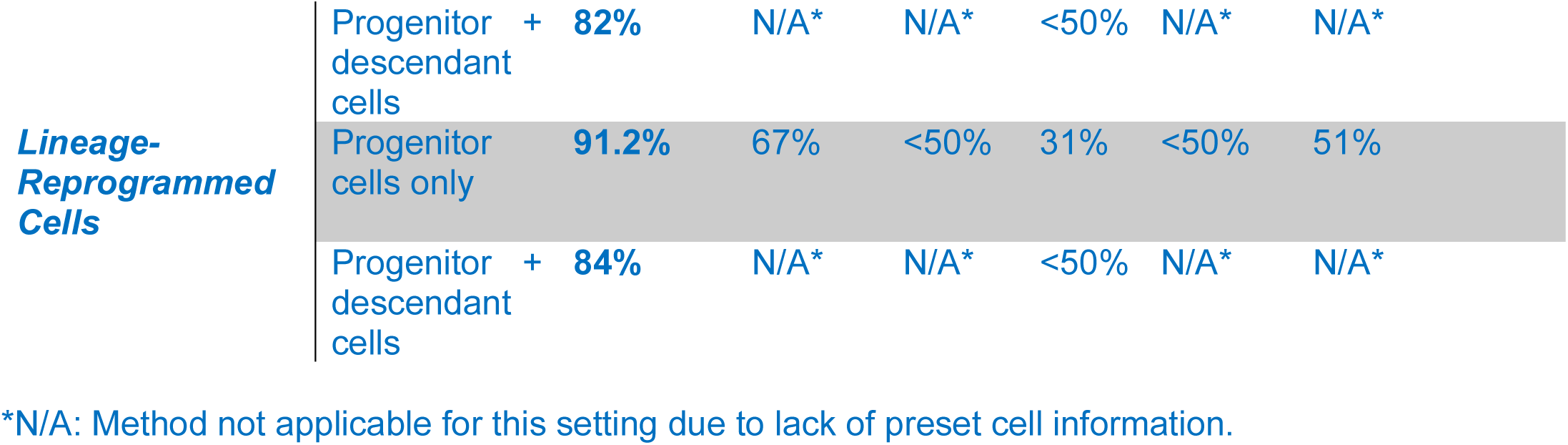
Comparative Performance of Lineage Inference Methods(Dataset 1)

TOGGLE demonstrated superior performance across all evaluated scenarios (Figure 5A-H). For mouse bone marrow hematopoietic progenitor cells, TOGGLE achieved 79% accuracy when evaluating progenitor cells alone, substantially outperforming Cospar (65%), Super OT (67%), WOT (65%), GAN-Based OT (62%), and Deep-Lineage (52%). When extending the evaluation to include both progenitor and descendant cells, TOGGLE’s accuracy increased to 82%, while WOT’s performance dropped below 50%; other methods were not applicable to this setting as they require preset cell information. For lineage-reprogrammed cells, TOGGLE’s performance advantage was even more pronounced, achieving 91.2% accuracy for progenitor cells compared to 67% for Cospar, 51% for Deep-Lineage, and only 31% for WOT. In the combined progenitor-descendant evaluation, TOGGLE maintained 84% accuracy, again substantially exceeding WOT (<50%), with other methods remaining inapplicable, write as N/A. These results demonstrate TOGGLE’s robust performance as an unsupervised method, particularly in scenarios lacking prior cell-type annotations.

**Figure 5-1.**
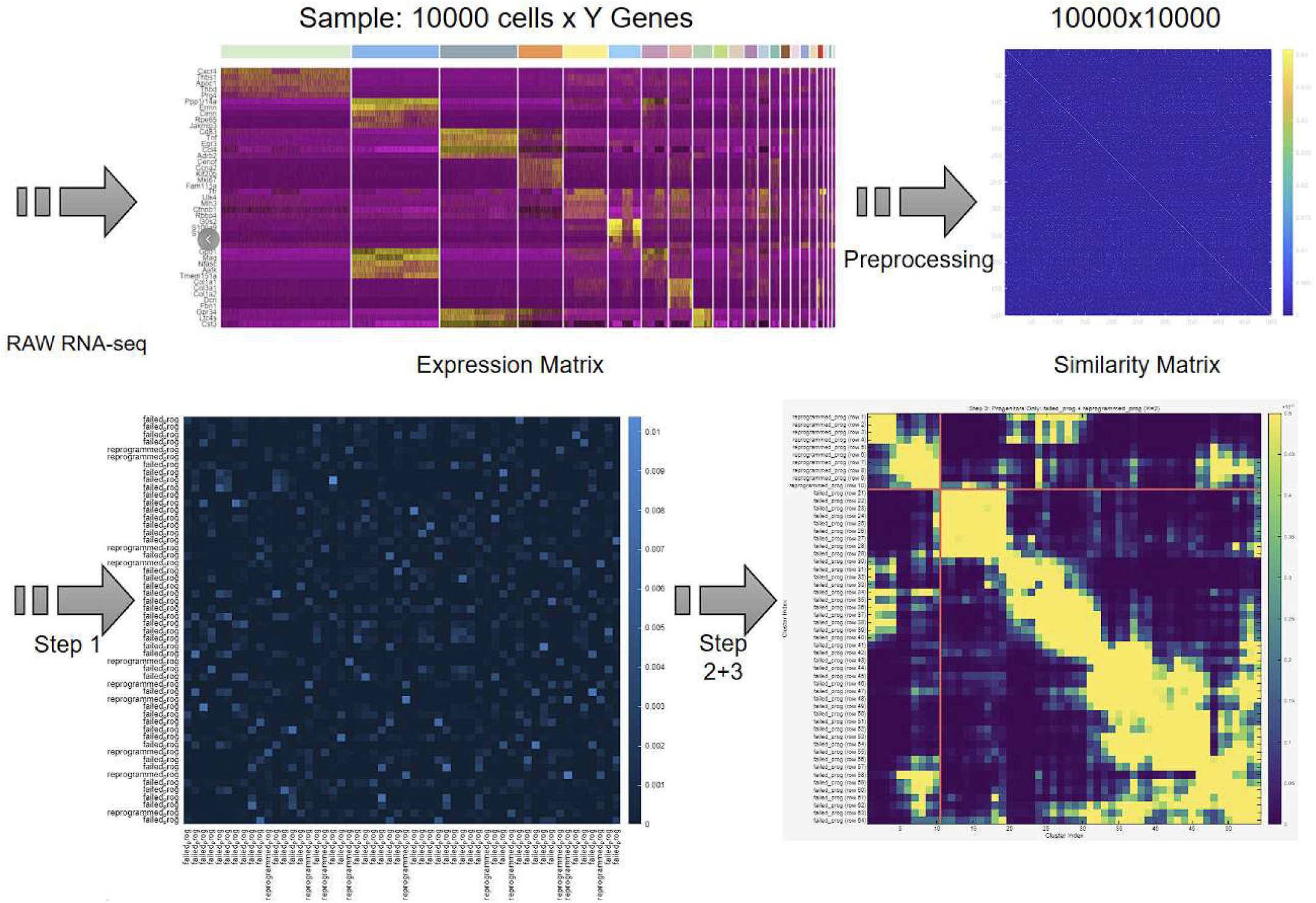
The text contains hypothetical fictional data. The image materials are derived from cell reprogramming cases and serve solely as simulated illustrations to explain the stepwise procedures. Input raw RNA sequencing data. Generate an expression matrix. After preprocessing, produce a cell similarity matrix [10,000 x 10,000]. Following Step 1, generate preliminary groupings [30 x 30]. After Steps 2 and 3, obtain the final result [30 x 30]. Scalable vector graphics for magnified viewing are provided in the submission system.

**Figure 5-2.**
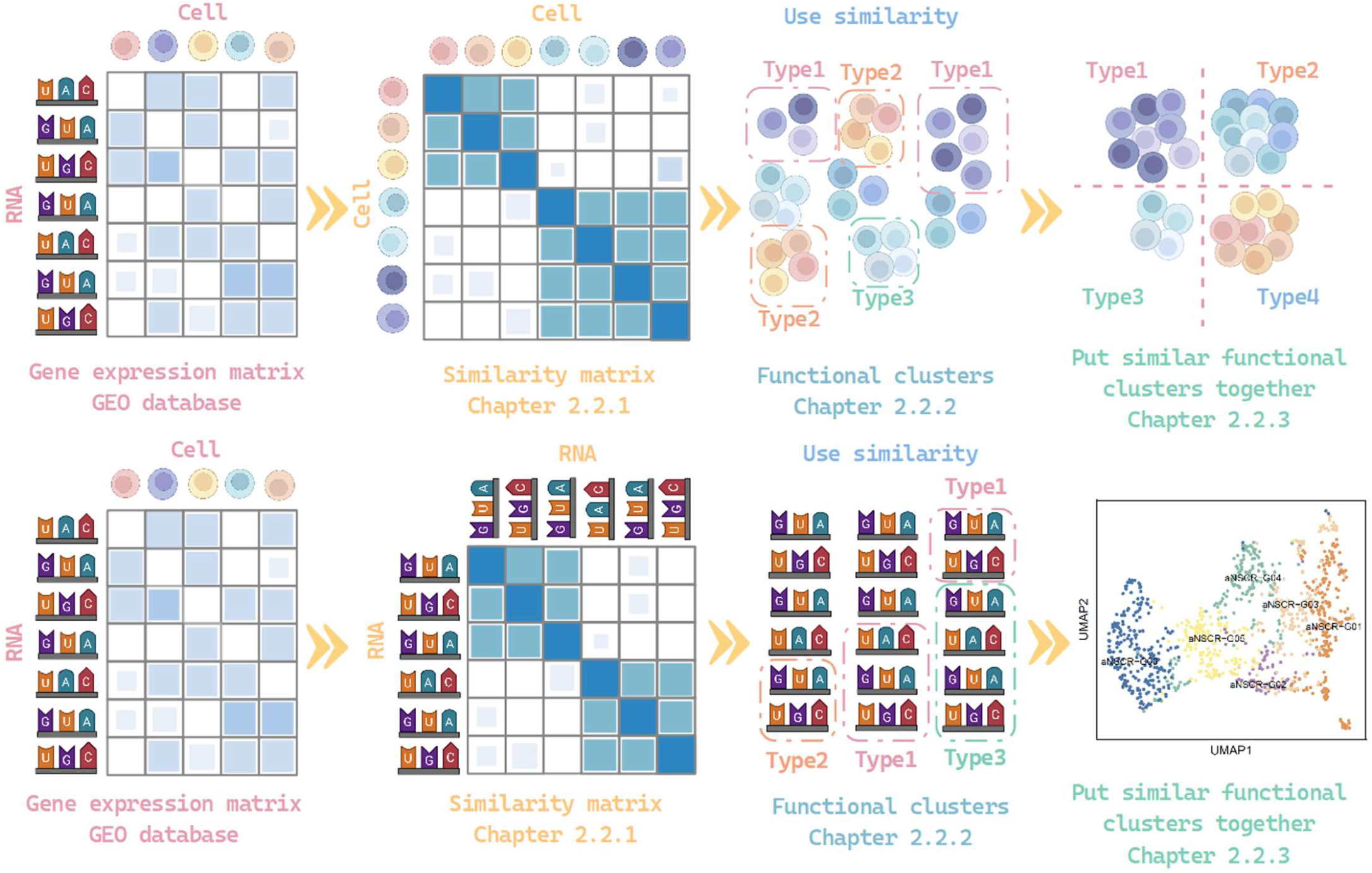
**Top**: Derivation of Cell Functions from a Gene Matrix. **Bottom**: Derivation of RNA functions from a Gene Matrix. A **gene expression matrix** is used to retrieve RNA transcript counts for individual cells from the GEO database. A **similarity matrix** calculates pairwise similarities between cells to enable classification. **Functional clusters** identify initial groups of cells with similar biological functions. And lastly, similar functional clusters are merged together.

**Figure 6.**
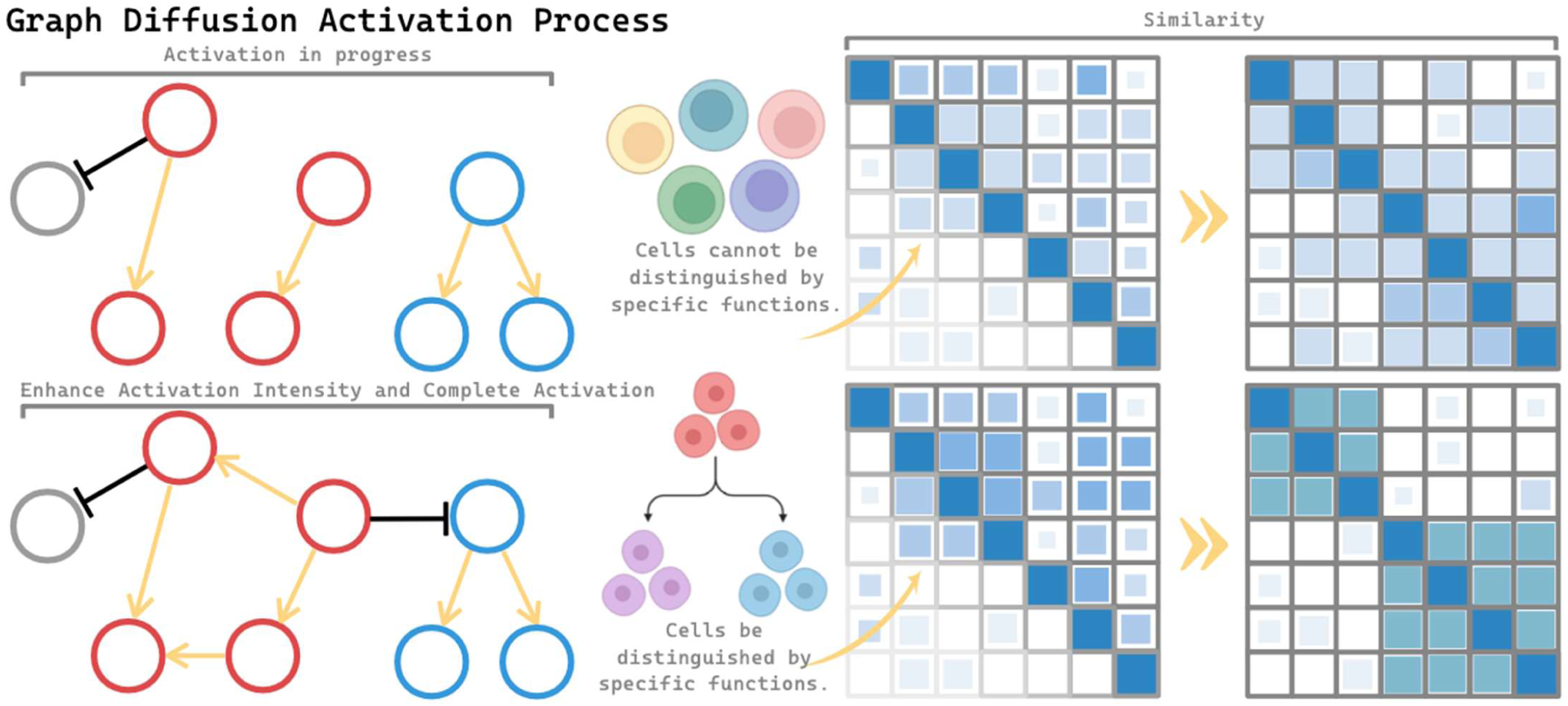
**Top panel:** In conventional models (left), the initial model activation is limited. Due to the masking effect of similar gene expression profiles (middle), functional differences between cells remain indistinct, resulting in poor separation in the data space (right). **Bottom panel:** By applying TOGGLE’s novel framework, deeper model activation is achieved (left). Cells are distinguished based on both functional identity and progression stage (middle), leading to more pronounced separation in the data space (right). The data separation is visibly more distinct.

**Figure 7.**
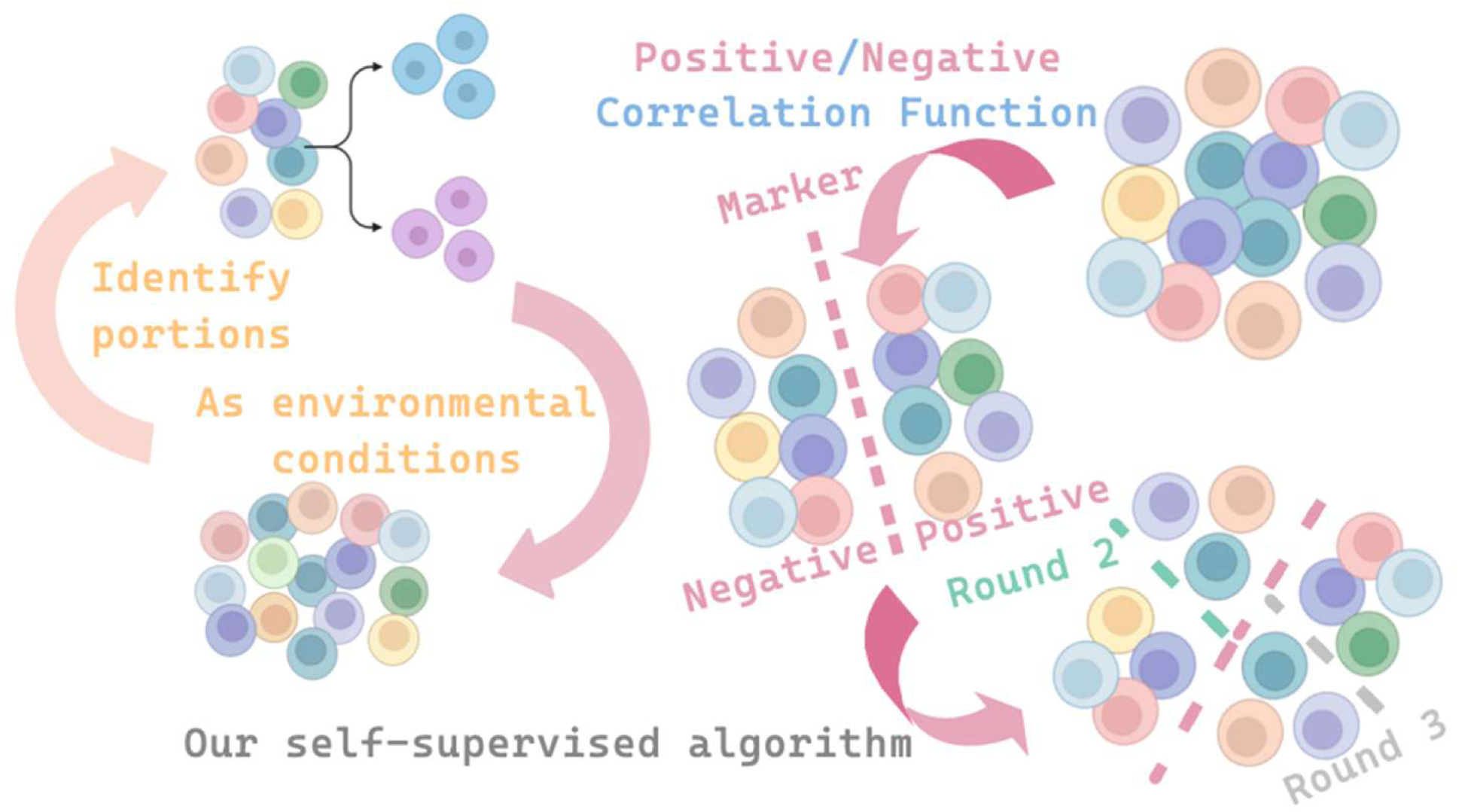
**Right:** The algorithm first generates a mathematical condition used to evaluate whether each cell is positively or negatively correlated. The boundary separating these two categories is defined as a “Marker.” Each pair of adjacent Markers delineates a cluster, comprising cells that exhibit similar functional characteristics. This process requires no manual labeling; the algorithm autonomously generates and refines Markers through iterative feedback, enabling unsupervised functional clustering. **Left:** The algorithm treats the task of identifying spatial regions of cell distribution as an output action (Action), which is then fed back as the subsequent state or environmental condition (State/Environment).

#### 3.1.2 Ablation Experiment Results

This ablation experiment aims to verify the effectiveness of each component in the TOGGLE algorithm by comparing two configurations:

- **Step 2 Only:** Using only binary correlation sorting, without genetic algorithm optimization or automatic boundary detection.
- **Full TOGGLE:** Applying the complete three-step process (Binary Correlation Sorting + Genetic Algorithm + Cluster combine), as the second and third steps are bound together.

**Table 2:** Comparison of TOGGLE Ablation Experiment Results. This table compares the performance between Step 2 Only (using only binary correlation sorting) and Full TOGGLE (the complete three-step process).

| Dataset | Task type | Metric | Step 2 Only | Full TOGGLE | Performance improvement | Conclusion |
| --- | --- | --- | --- | --- | --- | --- |
| Hematopoiesis (n=3,394) | Task 2 Cell Type | K | 77 | 4 | ↓ 94.8% | Significant improvement |
|  |  | ARI | 0.0460 | 0.5629 | ↑ 1123% | Significant enhancement |
|  |  | NMI | 0.3276 | 0.5977 | ↑ 82.4% | Significant enhancement |
|  | Task 3 Lineage | K | 77 | 2 | ↓ 97.4% | Significant improvement |
|  |  | ARI | 0.3682 | 0.3668 | ≈ comparable | Already optimal |
|  |  | NMI | 0.2846 | 0.2835 | ≈ comparable | Already optimal |
| Reprogramming (n=1,741) | Task 2 Cell Type | K | 32 | 4 | ↓ 87.5% | Significant improvement |
|  |  | ARI | 0.0546 | 0.7139 | ↑ 1207% | Significant enhancement |
|  |  | NMI | 0.3086 | 0.5777 | ↑ 87.2% | Significant enhancement |
|  | Task 3 Lineage | K | 32 | 2 | ↓ 93.8% | Significant improvement |
|  |  | ARI | 0.6637 | 0.6495 | ≈ comparable | Already optimal |
|  |  | NMI | 0.4869 | 0.4603 | ≈ comparable | Already optimal |

The full TOGGLE demonstrates superior performance in cell type identification tasks.

##### For the Hematopoiesis dataset (hematopoietic cells)

Cluster number optimization: Precisely reduced from 77 clusters to 4 (the true number of cell types), with an optimization rate of 94.8%.
ARI performance: Surged from 0.0460 to 0.5629, achieving an improvement of 1123% (12.2 times). NMI performance: Increased from 0.3276 to 0.5977, with a growth of 82.4%.

##### For the Reprogramming dataset (reprogramming cells)

Cluster number optimization: Precisely reduced from 32 clusters to 4, with an optimization rate of 87.5%.
ARI performance: Soared from 0.0546 to 0.7139, achieving an improvement of 1207% (13.1 times).
NMI performance: Increased from 0.3086 to 0.5777, with a growth of 87.2%.
In lineage tracing tasks, the first step of TOGGLE is already close to optimal.

For the coarse-grained lineage classification task (Task 3), the performance of Step 2 Only is comparable to that of Full TOGGLE. Differences in both ARI and NMI metrics are less than 6%. This indicates that binary correlation sorting alone is sufficient to capture the primary signals of lineage differentiation. The additional optimization provided by the genetic algorithm yields only limited marginal gains. The key insight is as follows: the complete TOGGLE pipeline (genetic algorithm optimization combined with automatic boundary detection) is essential for fine-grained cell type identification, where it can improve clustering accuracy by more than tenfold. In contrast, for coarse-grained lineage classification, simple binary correlation sorting proves sufficiently effective.

However, from a structural perspective, the lineage tracing task involves only the classification of progenitor cells during development. In actual experimental processes, if the algorithm can simultaneously identify progenitor cells during development and their post-development counterparts, it can greatly simplify the experimental workflow and reduce funding expenditures. This capability allows the algorithm to generalize to cell sampling at any time point, without being restricted to cells solely in development or fully developed cells. Therefore, the entire structure remains necessary.

In certain tasks, the third step’s calculations resulted in a slight decline. However, the decline was minimal (0.664 to 0.650, Δ=0.014), attributable to fundamental differences in label assignment methods between the two steps. In the “Step 2 only” results, each of the 32 clusters independently undergoes majority voting. Clusters do not need to be adjacent—for example, cluster #5 and cluster #28 are far apart in the reordered sequence, yet both can be labeled as “reprogrammed_prog”. This flexibility means that scattered cells within the “correct” majority clusters can still be correctly assigned. However, in the complete algorithm steps, a single continuous cut is performed on the progenitor cell submatrix. Everything above the boundary is assigned to Group A, and everything below the boundary to Group B. Even if only 1-2 boundary clusters are located on the wrong side of the cut line, those cells will be misclassified. A Δ of 0.014 indicates that approximately 25 cells out of 1741 progenitor cells have undergone a change in assignment strategy. This represents essentially stable performance—the structure captured by continuous boundaries is identical to that of non-continuous merging, with only a minor performance loss due to the continuity constraint. Nevertheless, this approach remains necessary, as we expect cells of the same type, such as “reprogrammed_prog”, to be merged into a single group as much as possible, rather than spread across 13 groups. This constitutes an unresolved issue in the algorithm.

#### 3.1.3 Benchmark Comparison Experiment

This comparison experiment comprehensively contrasts the full TOGGLE with three classic clustering algorithms. To ensure fairness, all baseline methods employ Oracle-K—the optimal number of clusters predetermined by TOGGLE—thereby eliminating the impact of cluster number selection and purely assessing the merits of the algorithms’ core mechanisms.

- **K-means:** A centroid-based clustering algorithm that uses Euclidean distance and optimizes the within-cluster sum of squares through iteration.
- **Spectral Clustering:** A clustering algorithm based on the eigenvalue decomposition of the graph Laplacian matrix.
- **Hierarchical Clustering:** A clustering algorithm based on distance metrics, using Ward linkage.

TOGGLE demonstrates superior performance in cell type identification tasks.

**Table 3:** Performance Comparison Between TOGGLE and Classic Clustering Algorithms. All baseline methods use Oracle-K—the optimal number of clusters predetermined by TOGGLE.

| Datasets | Tasks | Metric | Full<br>TOGGLE | K-means | Spectral | Hierarchical |
| --- | --- | --- | --- | --- | --- | --- |
| **Hematopoiesis**<br>(n=3,394) | **Task 2**<br>**Cell Type** | **K** | **4 ★** | 4 | 4 | 4 |
|  |  | **ARI** | **0.5629 ★** | 0.0099 | 0.0107 | 0.3329 |
|  |  | **NMI** | **0.5977 ★** | 0.0809 | 0.0637 | 0.4863 |
|  | **Task 3**<br>**Lineage** | **K** | **2 ★** | 2 | 2 | 2 |
|  |  | **ARI** | **0.3668 ★** | 0.0532 | -0.0004 | -0.0006 |
|  |  | **NMI** | **0.2835 ★** | 0.1225 | 0.0015 | 0.0016 |
| **Reprogramming**<br>(n=1,741) | **Task 2**<br>**Cell Type** | **K** | **4 ★** | 4 | 4 | 4 |
|  |  | **ARI** | **0.7139 ★** | 0.1060 | 0.2187 | 0.1437 |
|  |  | **NMI** | **0.5777 ★** | 0.2877 | 0.3523 | 0.2827 |
|  | **Task 3**<br>**Lineage** | **K** | **2 ★** | 2 | 2 | 2 |
|  |  | **ARI** | **0.6495 ★** | 0.0006 | -0.0629 | -0.0616 |
|  |  | **NMI** | **0.4603 ★** | 0.0002 | 0.0470 | 0.0178 |

##### For the Hematopoiesis dataset (Task 2 - Cell Type)

ARI ranking: TOGGLE (0.5629) >> Hierarchical (0.3329) >> Spectral (0.0107) ≈ K-means (0.0099). Relative improvement over the best baseline: TOGGLE outperforms Hierarchical by 69.1%.

##### For the Reprogramming dataset (Task 2 - Cell Type)

ARI ranking: TOGGLE (0.7139) >> Spectral (0.2187) > Hierarchical (0.1437) > K-means (0.1060).

Relative improvement over the best baseline: TOGGLE outperforms Spectral by 226.5% (exceeding three times the performance).

TOGGLE also maintains a leading position in lineage tracing tasks. Even in relatively simple lineage classification tasks (Task 3), TOGGLE significantly outperforms all baseline methods. Notably, Spectral and Hierarchical produce negative ARI values in certain scenarios, indicating that their clustering results are even inferior to random assignment. This fully reveals the limitations of traditional methods in processing single-cell data.

This algorithm addresses the following typical problems present in the baselines:

K-means faces a dual dilemma. It performs the worst across all eight evaluation metrics, with an average adjusted Rand index of only 0.04. The primary reasons are as follows:

① The curse of dimensionality: Single-cell data typically features thousands to tens of thousands of gene dimensions, causing Euclidean distance to fail in high-dimensional spaces.

②The spherical assumption: K-means assumes clusters have convex boundaries, making it unable to capture the nonlinear complex structures of cell states.

**Spectral clustering faces a computational challenge.** Although it can theoretically capture non-convex structures, in practice, it is constrained by the construction of similarity matrices and feature selection, resulting in unstable performance on single-cell data (with adjusted Rand index values ranging from 0.02 to 0.35).

**Hierarchical clustering exhibits strong data dependency.** It achieves relatively good results in Task 2 of the Hematopoiesis dataset (adjusted Rand index of 0.33) but completely fails in the other three tasks (adjusted Rand index below 0.15), demonstrating its severe reliance on specific dataset characteristics and lack of robustness.

**The insight from the Oracle-K advantage.** Even when all baseline methods are provided with Oracle-K—the optimal number of clusters determined by TOGGLE—as prior knowledge, their performance still lags far behind TOGGLE. This fully proves that **TOGGLE’s core strength lies in its unique three-step algorithmic mechanism, rather than simply the accurate selection of cluster numbers.**

TOGGLE achieves the best performance across all 12 evaluation dimensions, spanning two datasets, two tasks, and three metrics, demonstrating significant advantages over traditional clustering methods.

### 3.2 Test 2: Distinguish the stages of ferroptosis and perform biological validation

Detailed protocols for all animal experiments are provided and can be found in the **Supplementary Appendix 6**. We utilized the GSE232429 dataset to identify and characterize neuronal cell death associated with ischemic stroke, including both middle cerebral artery occlusion (MCAO) and healthy control (Sham) samples. Throughout the study, MCAO and Sham samples were combined and submitted to the algorithm for integrated clustering, followed by separate visualization based on their respective labels.

#### 3.2.1 Identification of Programmed Cell Death by TOGGLE

The above results confirm the reliability of TOGGLE for predicting cell fate. In cases of ischemic stroke, neurons often undergo programmed cell death, with ferroptosis playing a significant role in neuronal loss. However, ferroptosis frequently overlaps with other forms of programmed cell death, making it challenging for existing algorithms to effectively distinguish ferroptosis from other cell death mechanisms.

We first utilized the Seurat software package to construct the single-cell sequencing matrix. We performed quality control on the neurons from the GSE232429 dataset using the criteria: 500 < nFeature_RNA < 8000, mitochondrial gene expression percentage (mt_percent) < 30%, red blood cell gene proportion < 5%, and nCount_RNA < 98th percentile. After constructing the single-cell sequencing expression matrix, we applied the TOGGLE algorithm to conduct stage identification analysis on the cells.

TOGGLE ultimately divided the neurons from the GSE232429 dataset into three cell groups: Surviving Neurons C1, Surviving Neurons C2, and Stressed Neurons (Figure 10). Among these, Surviving Neurons C1 accounted for a larger proportion in the Sham cohort, while Surviving Neurons C2 accounted for a larger proportion in the MCAO cohort.

**Figure 8.**
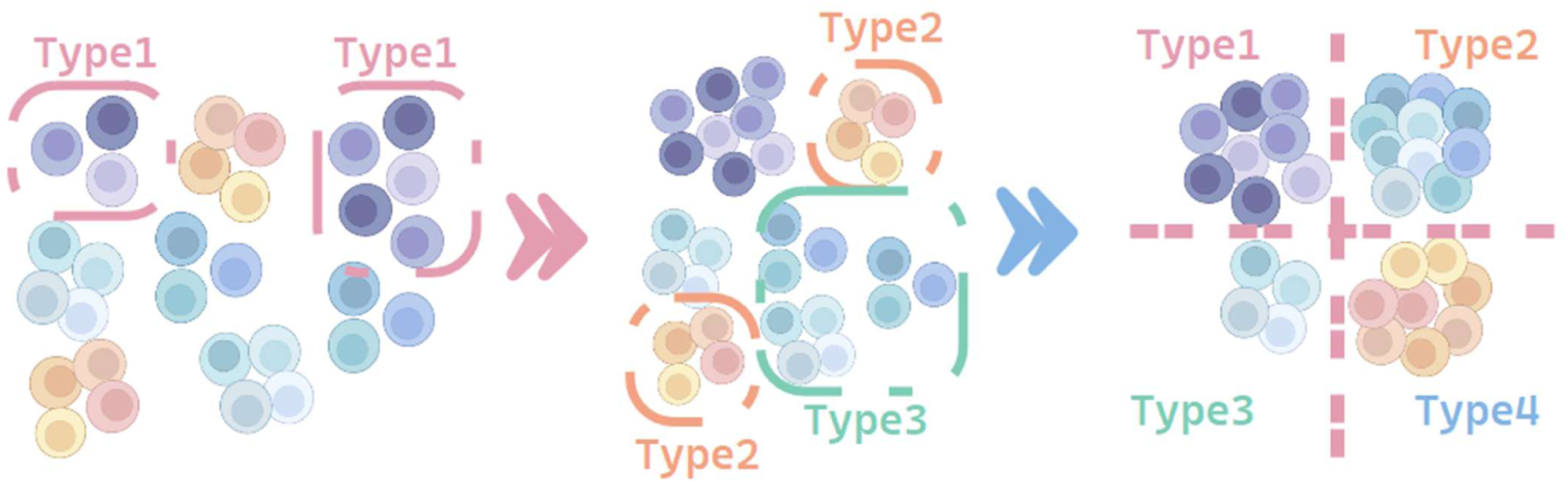
(1) In Section 2.2.2, the algorithm brings together cells with the same functional type (Type 1), even if they are in different conditions. (2) In Section 2.2.3, it further merges all cells with Functional Type 1 into a unified group. (3) Eventually, all cells with the same functional type—regardless of condition—are grouped together.

**Figure 9.**
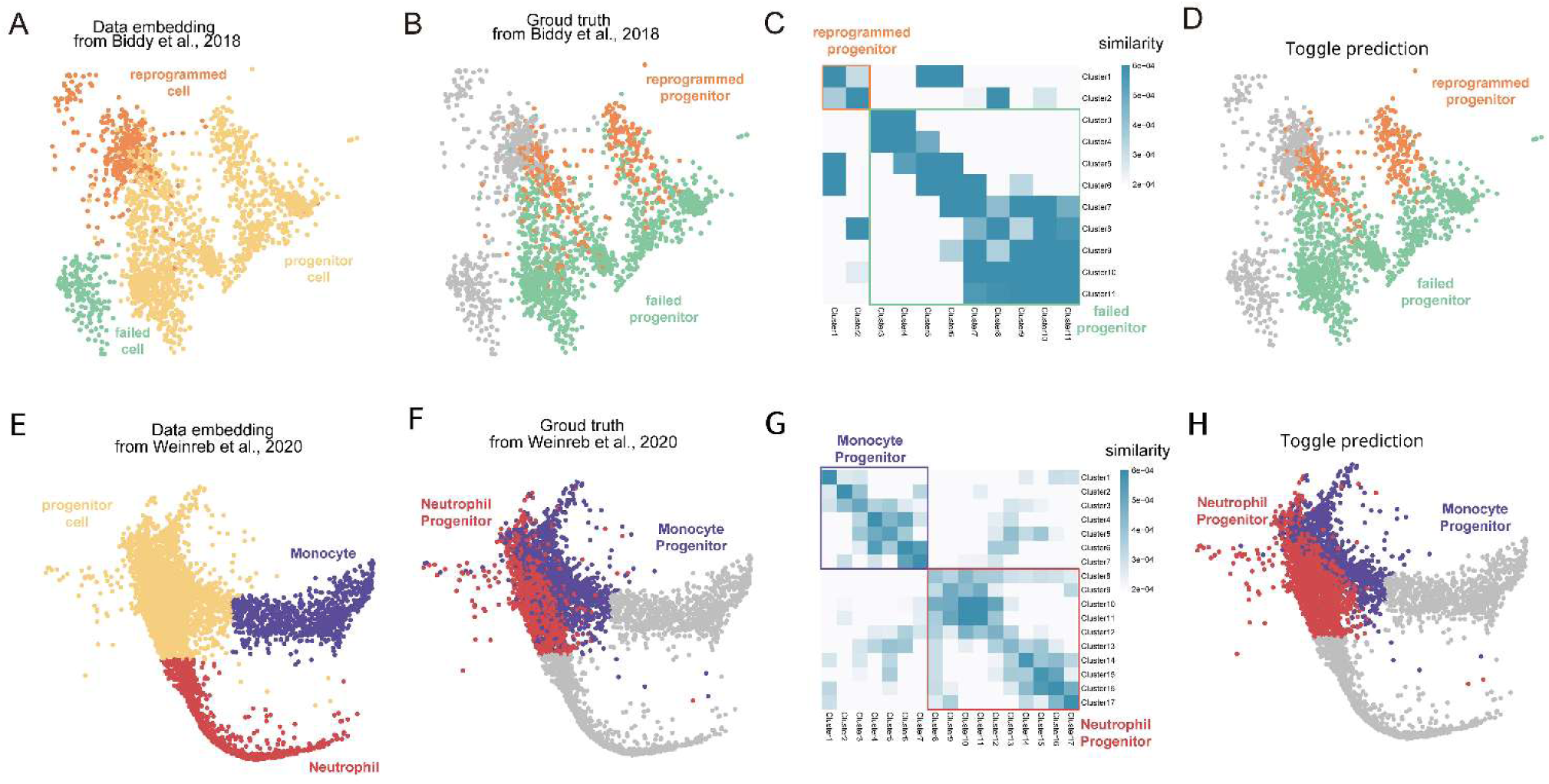
Tracing the lineage of fibroblast reprogramming⁹ and hematopoietic cell¹⁶ development. A/E: UMAP visualization of the original dataset. The Progenitor Cell population represents the targets to be predicted. **B/F:** Real label of lineage outcomes showing the eventual fate of Progenitor Cells. **C/G:** Algorithm-predicted results with output cluster boundaries. **D/H:** Final fate predictions based on the inferred trajectories.

**Figure 10.**
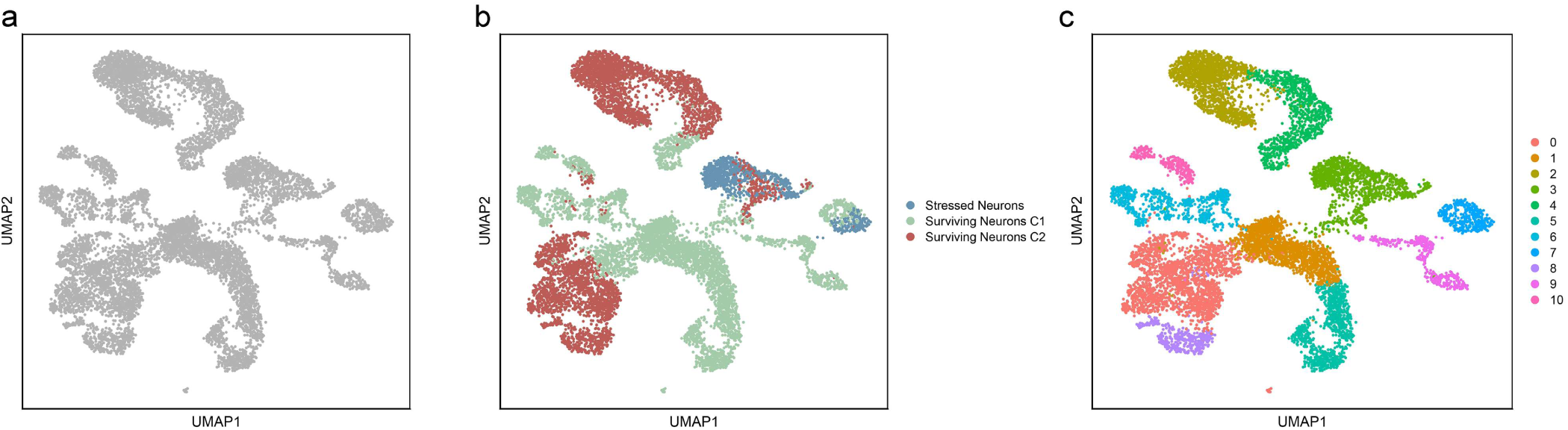
presents the UMAP visualization of the GSE232429 dataset (a: original UMAP; b: UMAP based on TOGGLE classification results; c: UMAP based on Seurat clustering). Seurat is one of the most commonly used cell classification toolkits, with its primary classification levels focusing on notable changes in gene expression patterns and differences among cell subtypes. This visualization serves to illustrate the distinctions from TOGGLE, revealing that standard classification levels cannot substitute for TOGGLE’s functional classification levels. The classification accuracy is verified in the following text.

#### 3.2.2 Accuracy Verification

Subsequently, we identified genes associated with the Surviving state, Stressed state, and Dying state based on previous literature. We then calculated the proportion of cells in each group exhibiting the corresponding state. This analysis revealed that more than 70% of cells in the Surviving Neurons C1 and Surviving Neurons C2 groups were classified as being in the Surviving state, whereas more than 70% of cells in the Stressed neurons group were classified as being in the Stressed state (Figure 11). To further validate this grouping, we performed differential gene expression analysis, enrichment analysis, and GSVA on Stressed neurons from the MCAO cohort and Surviving Neurons C1 from the Sham cohort.

**Figure 11.**
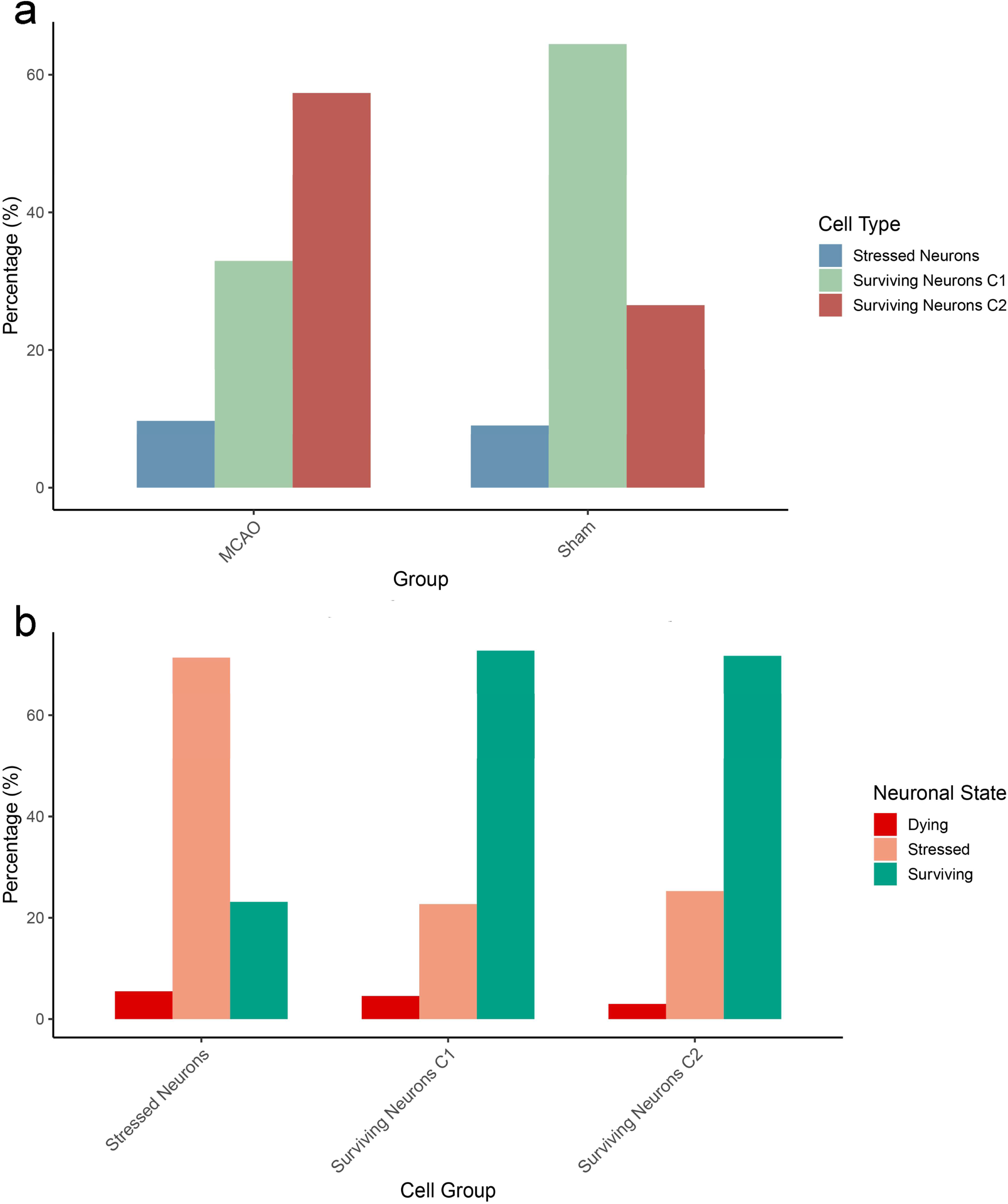
shows the proportion of cells in different categories (a: proportions of various cell types in the MCAO and Sham cohorts; b: proportions of different neuronal states within the three neuronal cell groups).

#### 3.2.3 Biological interpretation of the Stressed neuron group

Differential gene expression analysis revealed a total of 4,474 differentially expressed genes when comparing Stressed neurons from the MCAO cohort with Surviving Neurons C1 from the Sham cohort, of which 3,370 genes were upregulated and 1,104 genes were downregulated (Figure 12).

**Figure 12.**
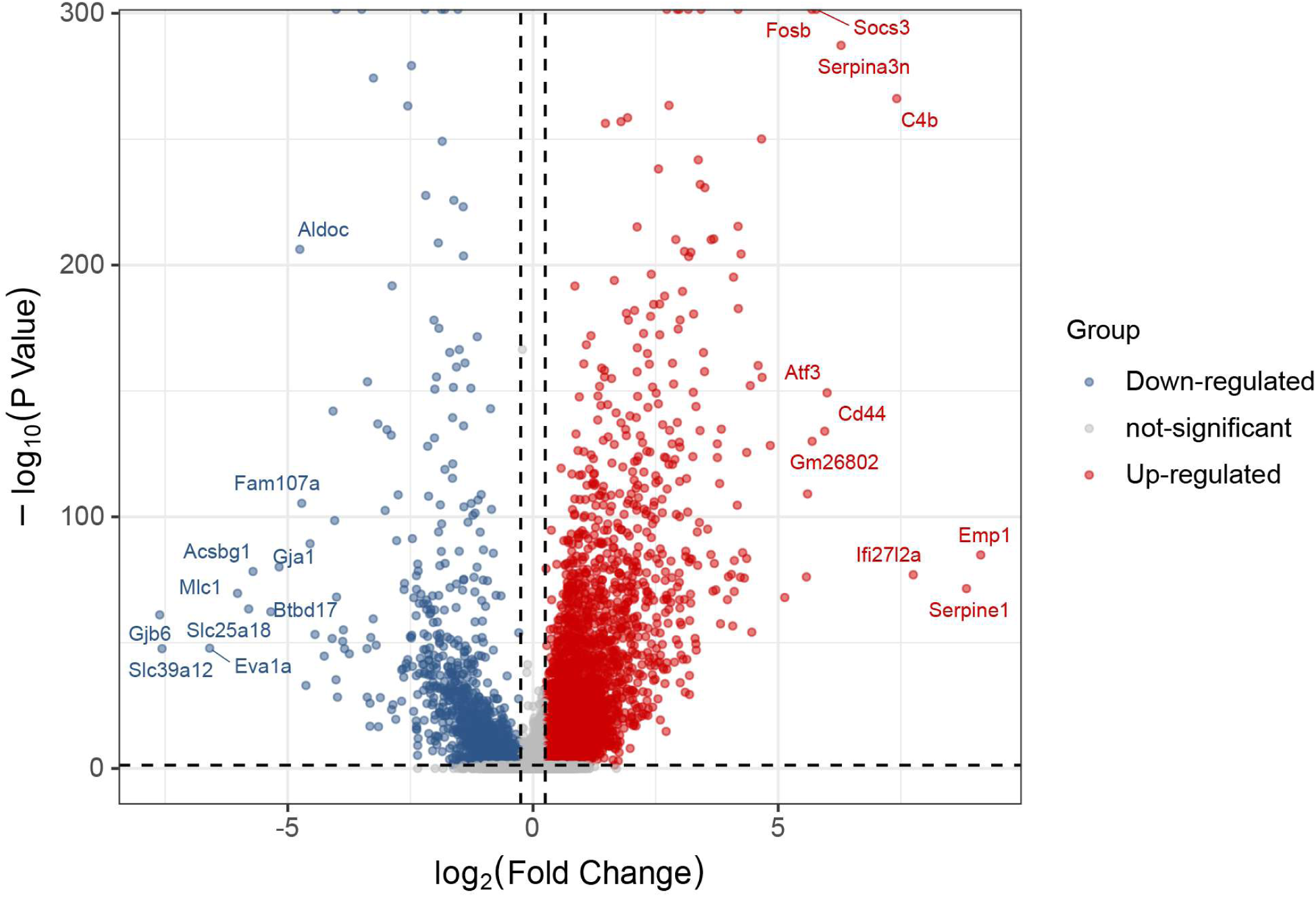
Volcano plot of differentially expressed genes. This volcano plot displays the results of differential gene expression analysis between Stressed Neurons from the MCAO cohort and Surviving Neurons C1 from the Sham cohort. Blue points represent genes significantly downregulated in Stressed Neurons, while red points represent genes significantly upregulated. These genes are involved in both pro- and anti-apoptotic signaling pathways as well as stress-related pathways, highlighting enhanced programmed cell death and stress responses in the Stressed group. This observation is further confirmed by the enrichment analysis presented later.

Enrichment analysis revealed that the biological processes, cellular components, molecular functions, and signaling pathways involved include neuronal apoptosis, iron metabolism, and ferroptosis (Figure 13). GSVA analysis further demonstrated pathway alterations in Stressed neurons (Table 4).

**Figure 13.**
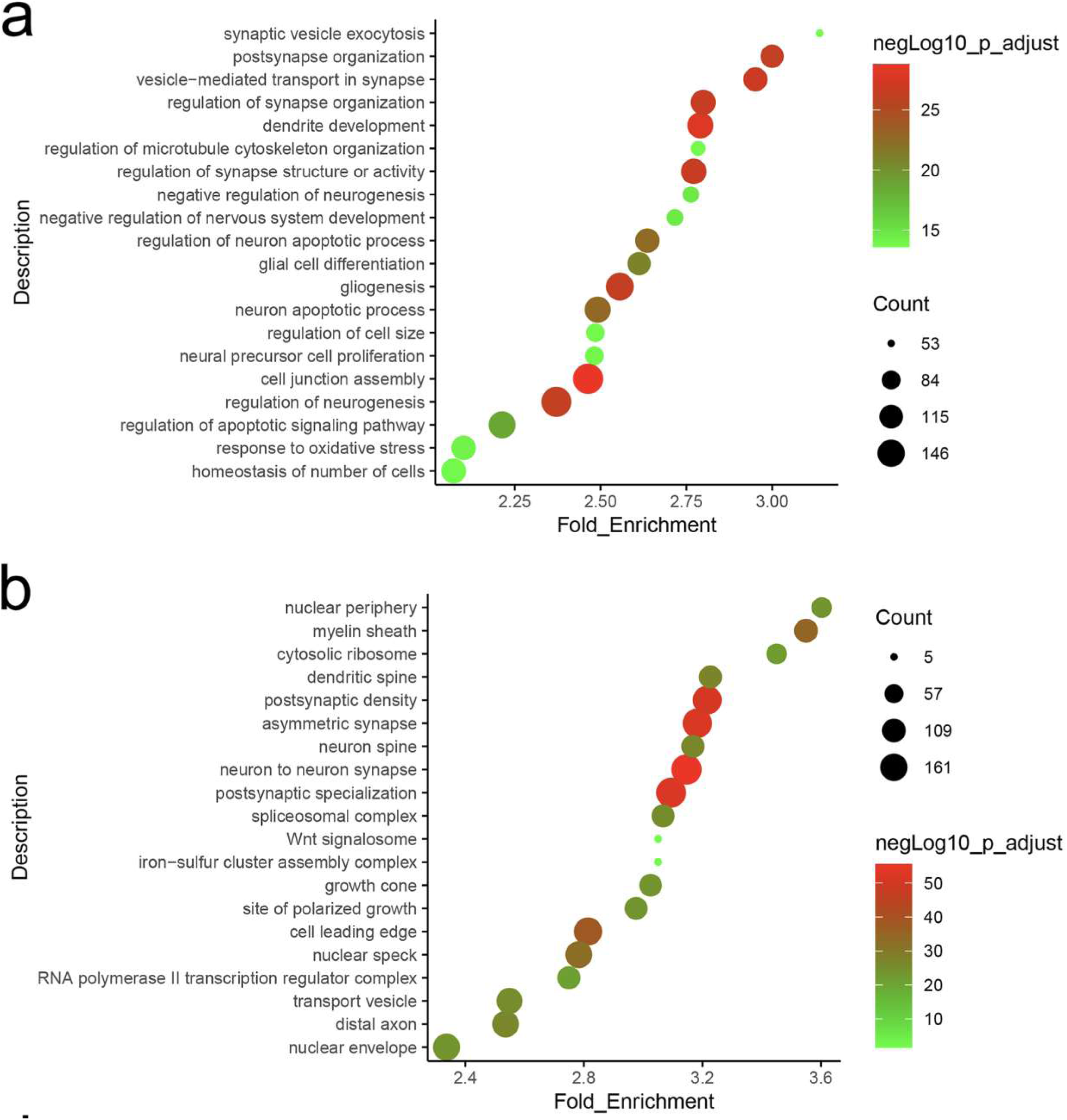
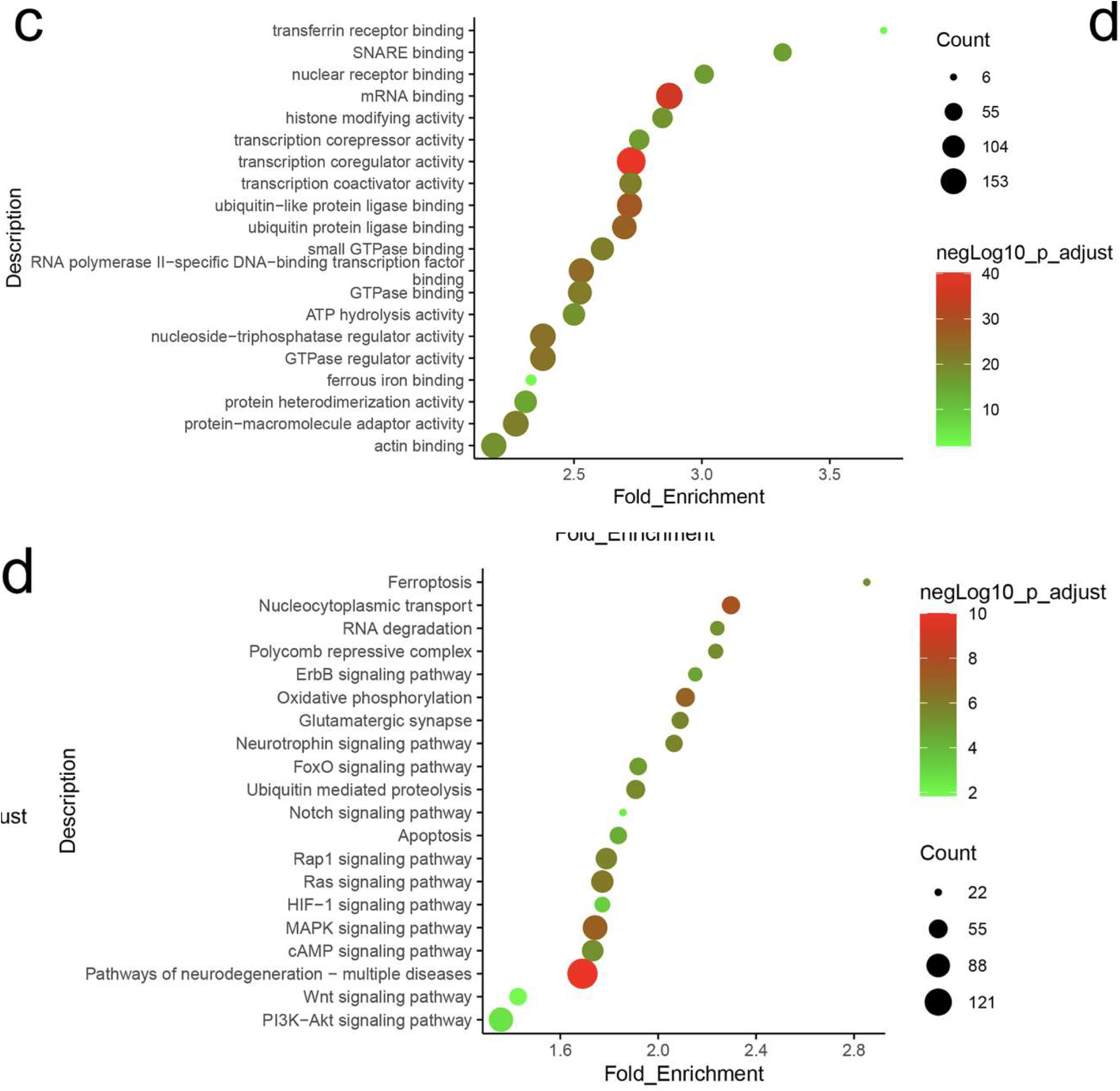
Bubble plot of enrichment results (a: BP; b: CC; c: MF; d: KEGG). This enrichment analysis figure (divided into panels a–d) reveals significant functional differences between Stressed Neurons and Surviving Neurons. The BP panel (a) highlights biological processes related to oxidative stress and neuronal apoptosis among the differentially expressed genes. The CC panel (b) emphasizes cellular components associated with the nuclear periphery and iron-sulfur clusters. The MF panel (c) focuses on transcriptional regulation and RNA-binding activity. The KEGG panel (d) shows that the differentially expressed genes are linked to signaling pathways including ferroptosis, apoptosis, and hypoxia.

**Table 4.** Pathway alterations in Stressed neurons.

| Pathway categories | Enhanced activity | Inhibited activity |
| --- | --- | --- |
| *Biological Process* | cellular response to iron ion,<br>regulation of iron ion transport,<br>multicellular organismal level iron ion homeostasis,<br>negative regulation of oxidative stress induced neuron intrinsic apoptotic signaling pathway | ferroptosis,<br>regulation of iron ion transmembrane transport,<br>iron ion import across plasma membrane,<br>negative regulation of ferroptosis |
| *Cellular Component* | death inducing signaling complex,<br>cytoplasmic stress granule,<br>nuclear stress granule,<br>HFE transferrin receptor complex | oxidoreductase complex, iron sulfur cluster assembly complex,<br>CD95 death inducing signaling complex |
| *Molecular Function* | death domain binding,<br>oxidative rna demethylase activity,<br>cysteine type endopeptidase activator activity involved in apoptotic process | death receptor activity,<br>cysteine type endopeptidase inhibitor activity involved in apoptotic process,<br>oxidative phosphorylation uncoupler activity |
| *REACTOME*<br>*Pathway Database* | apoptotic cleavage of cell adhesion proteins,<br>suppression of apoptosis,<br>apoptotic cleavage of cellular proteins, apoptotic execution phase,<br>mitochondrial fatty acid beta oxidation of unsaturated fatty acids | formation of apoptosome,<br>intrinsic pathway for apoptosis,<br>smac xiap regulated apoptotic response,<br>ros and rns production in phagocytes,<br>sulfide oxidation to sulfate |

Taken together, these results indicate that neuroprotective mechanisms—such as synaptic repair and transcriptional regulation—are activated in Stressed Neurons to counteract stress, yet dysregulation of iron metabolism may increase the risk of apoptosis.

Results are shown in Figure 14. Overall, the GSVA results simultaneously revealed processes that promote programmed cell death (PCD) and processes that inhibit PCD. This pattern is consistent with the competitive interplay between pro-PCD and anti-PCD mechanisms in cells undergoing the stressed state.

**Figure 14.**
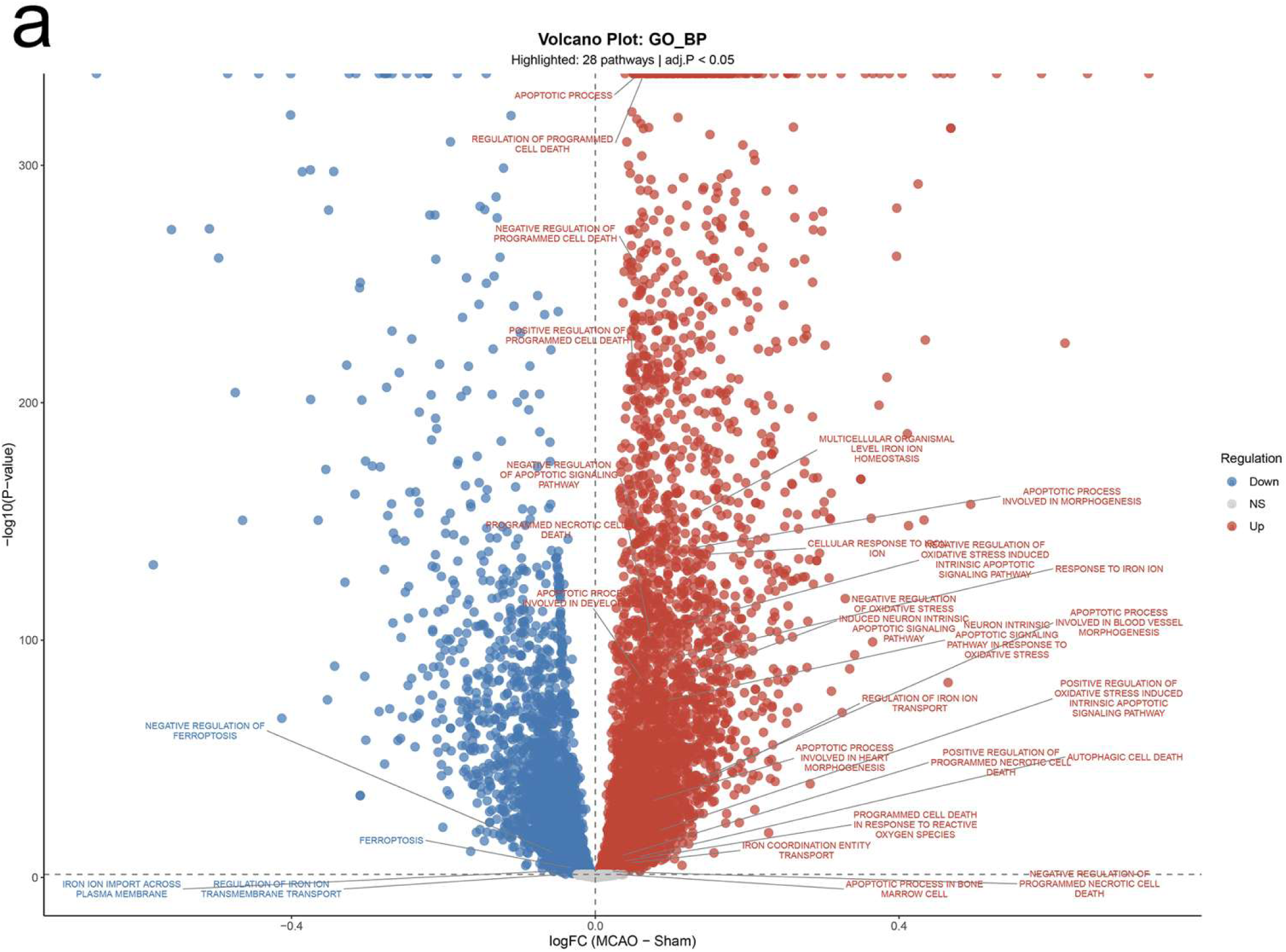
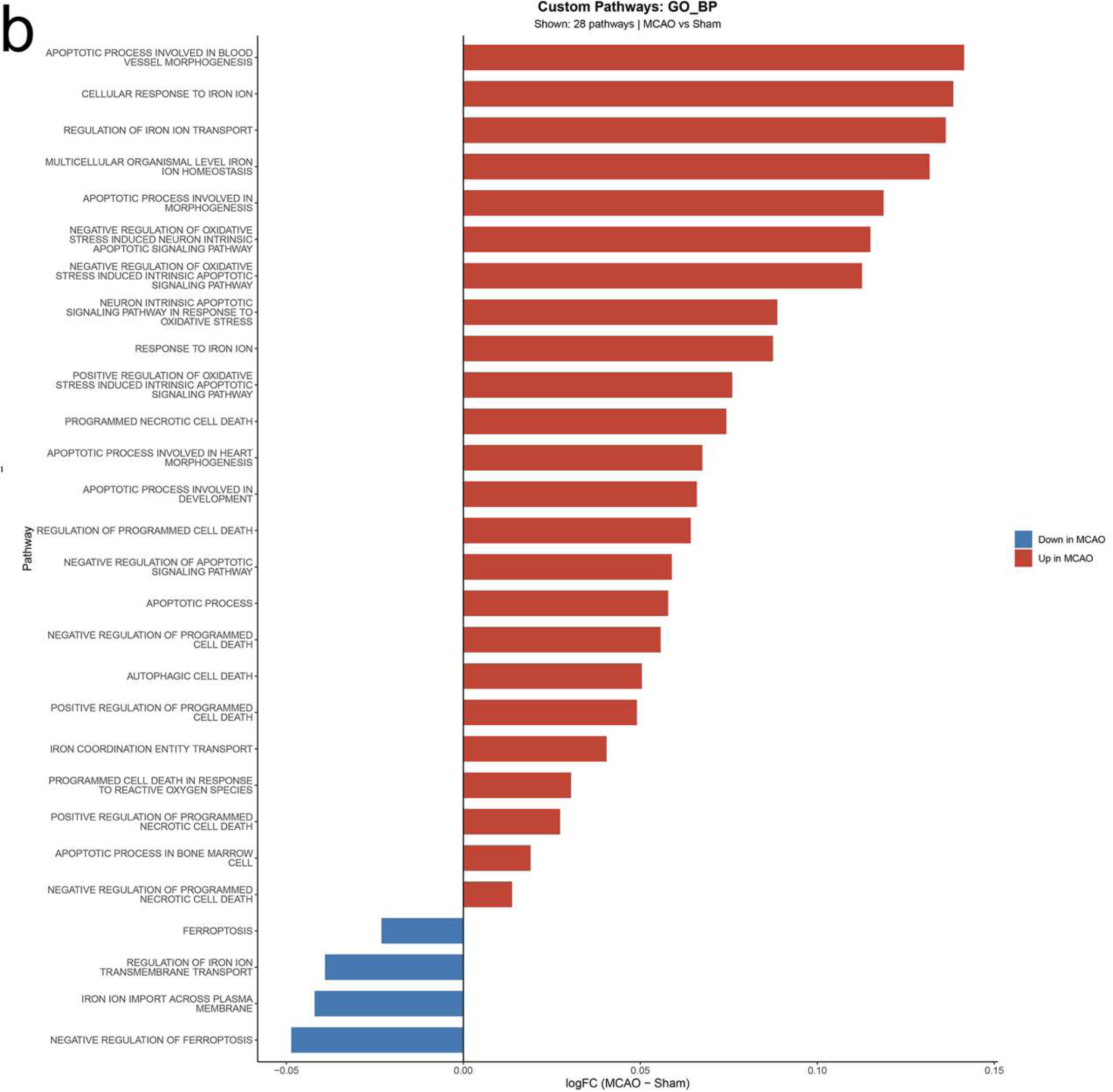
Volcano plot and bar plot of biological processes (BP) from GSVA analysis (a: volcano plot of BP; b: bar plot of BP). This figure reveals significant differences in biological processes between Stressed Neurons and Surviving Neurons. It demonstrates activation of processes related to programmed cell death, including neuronal apoptosis and iron metabolism, alongside relative suppression of processes such as ferroptosis and iron ion transport.

**Figure 15.**
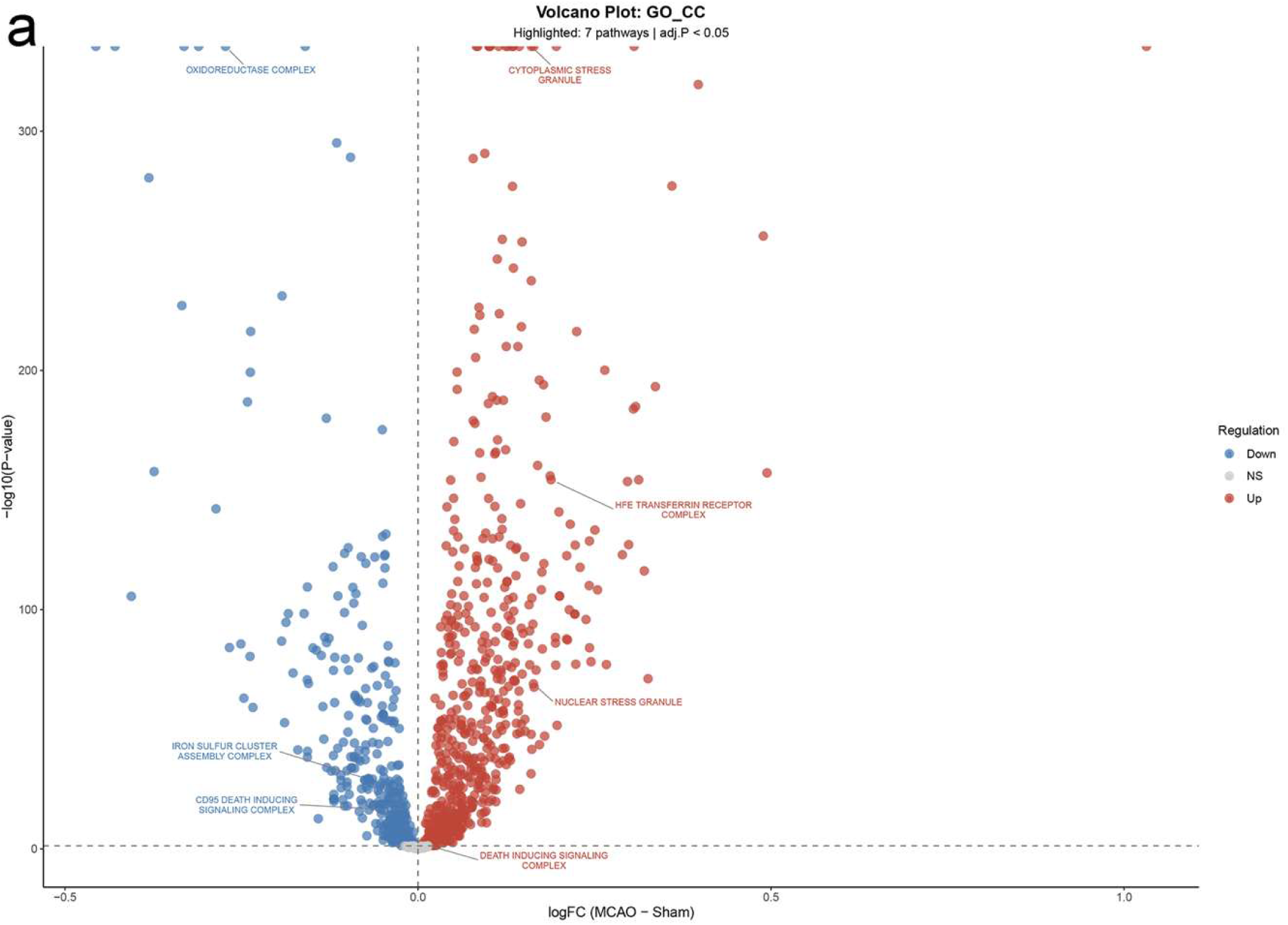
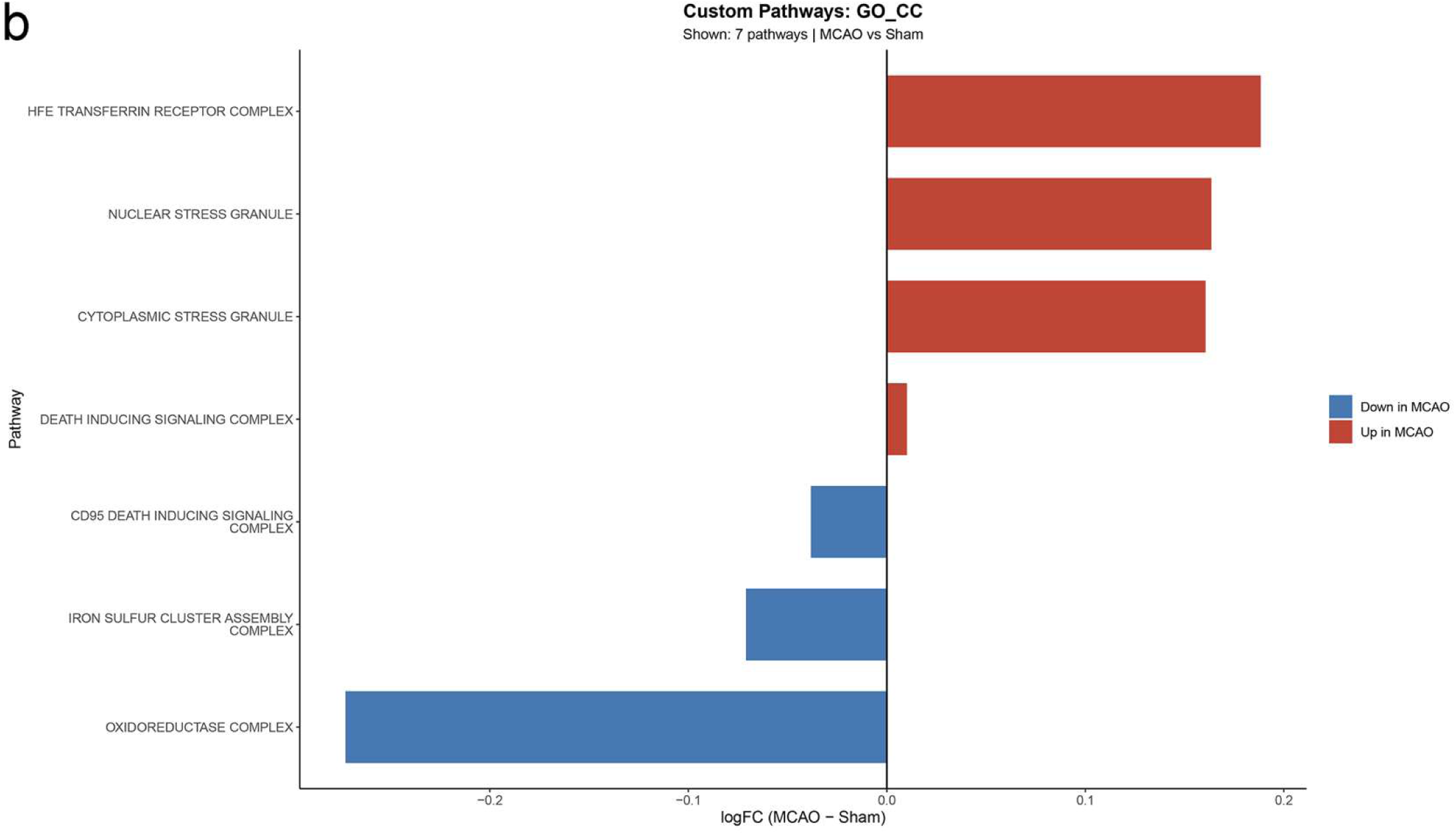
also includes volcano plots and bar plots of cellular components (CC) from GSVA analysis (a: volcano plot of CC; b: bar plot of CC). This figure reveals significant differences in cellular components between Stressed Neurons and Surviving Neurons, highlighting the activation and suppression of intracellular components associated with cell death.

**Figure 16.**
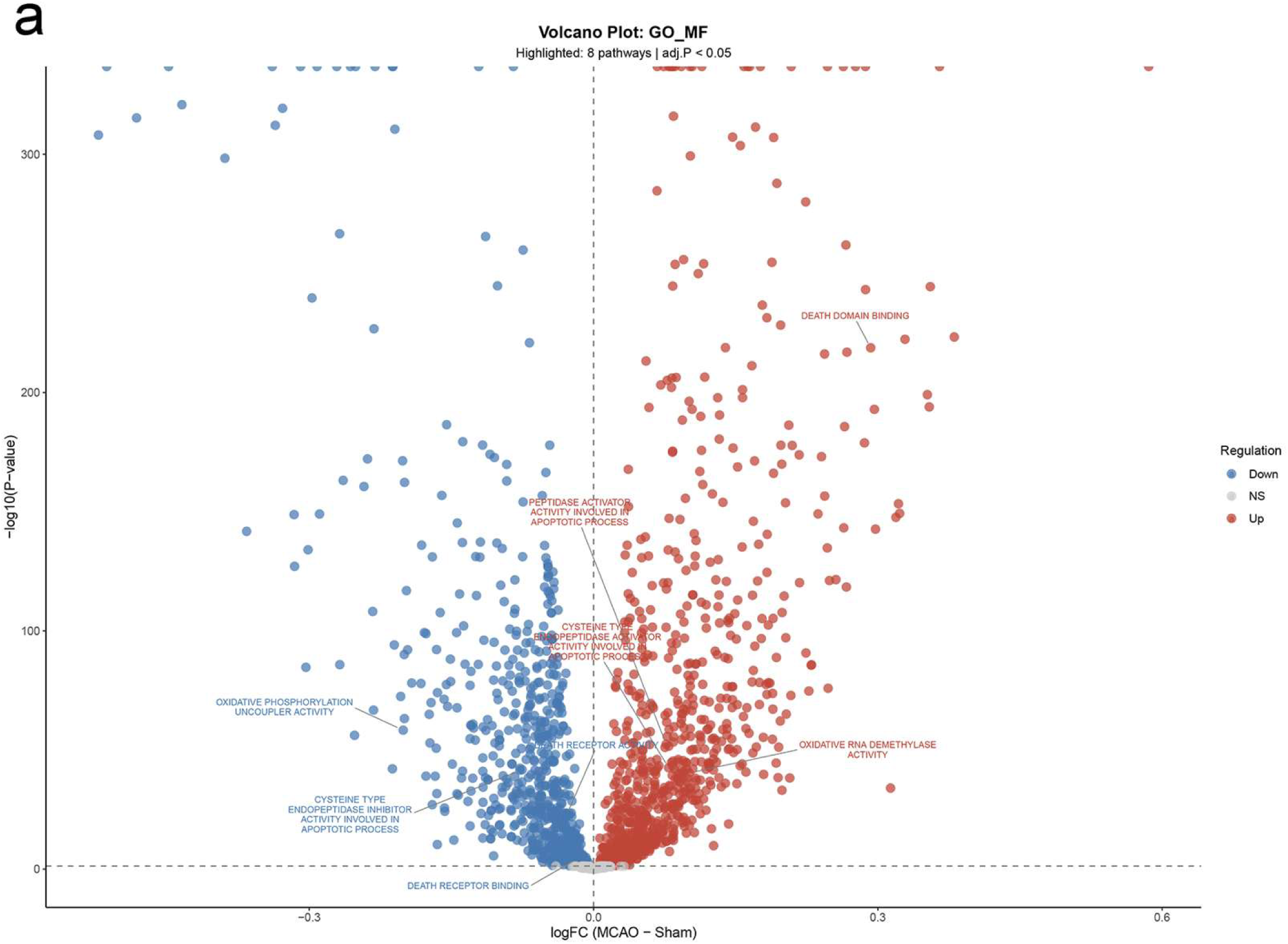
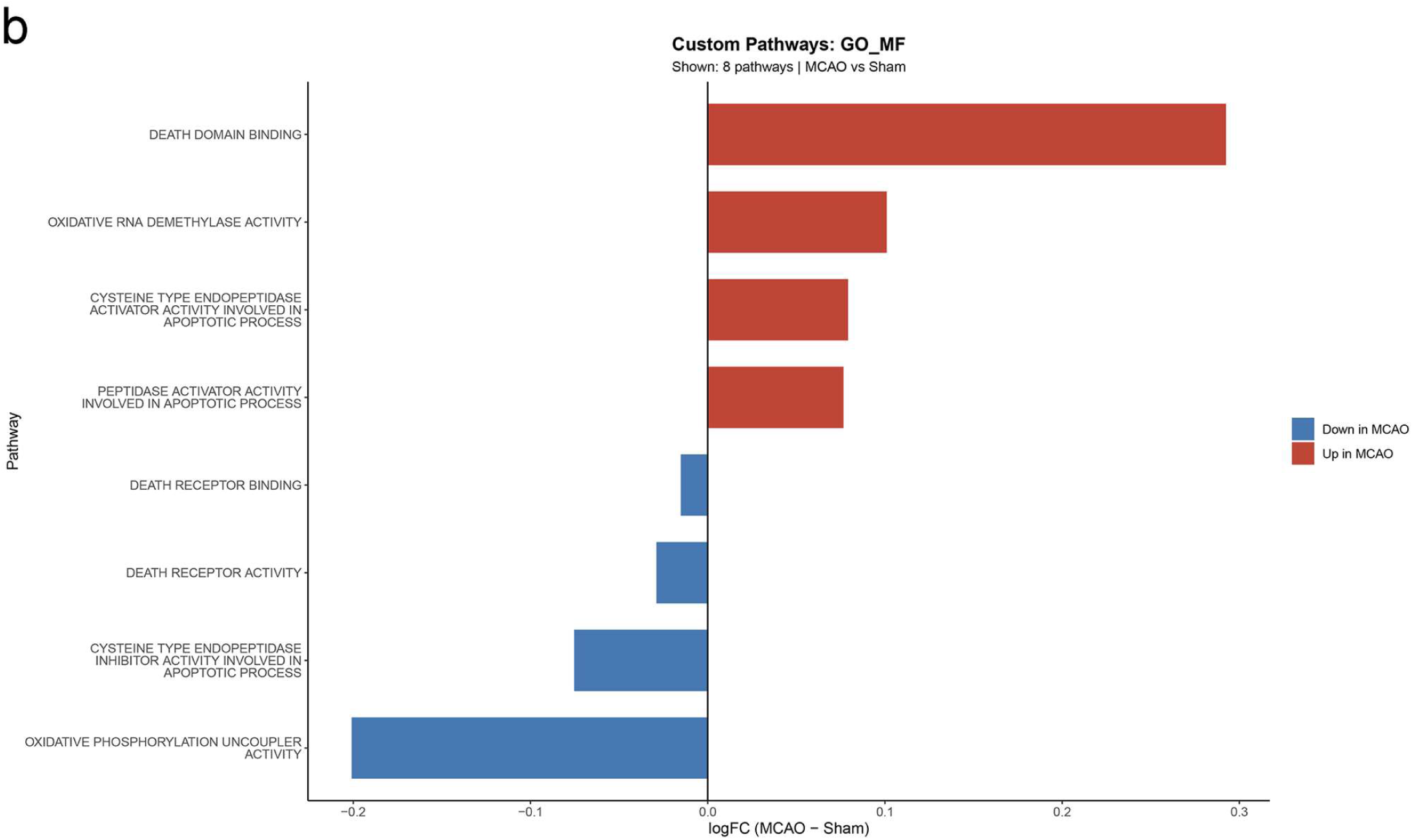
further presents volcano plots and bar plots of molecular functions (MF) from GSVA analysis (a: volcano plot of MF; b: bar plot of MF). This figure demonstrates significant differences in molecular functions between Stressed Neurons and Surviving Neurons, showing the relative activation and suppression of biomolecular functions related to cell death.

**Figure 17.**
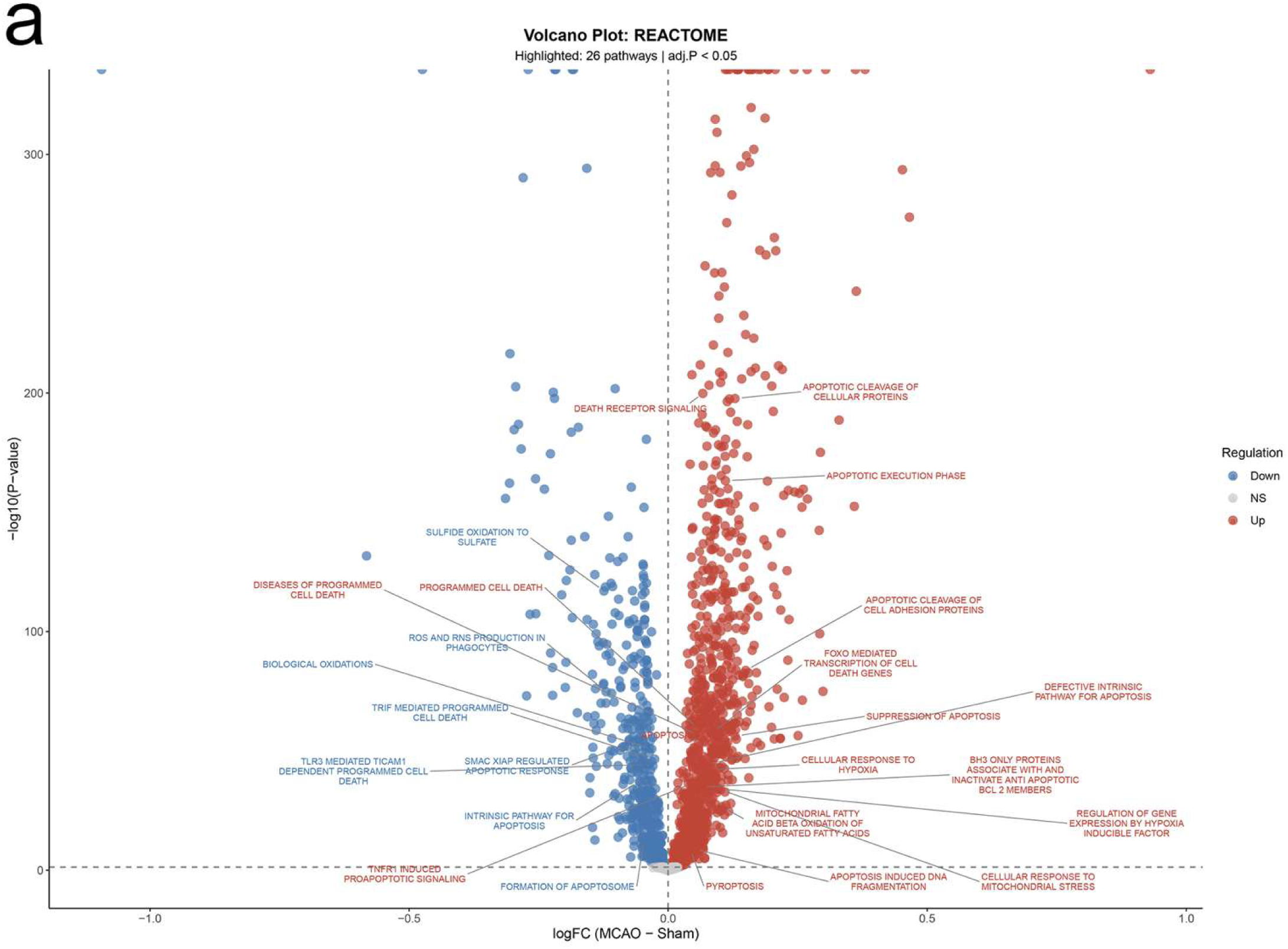
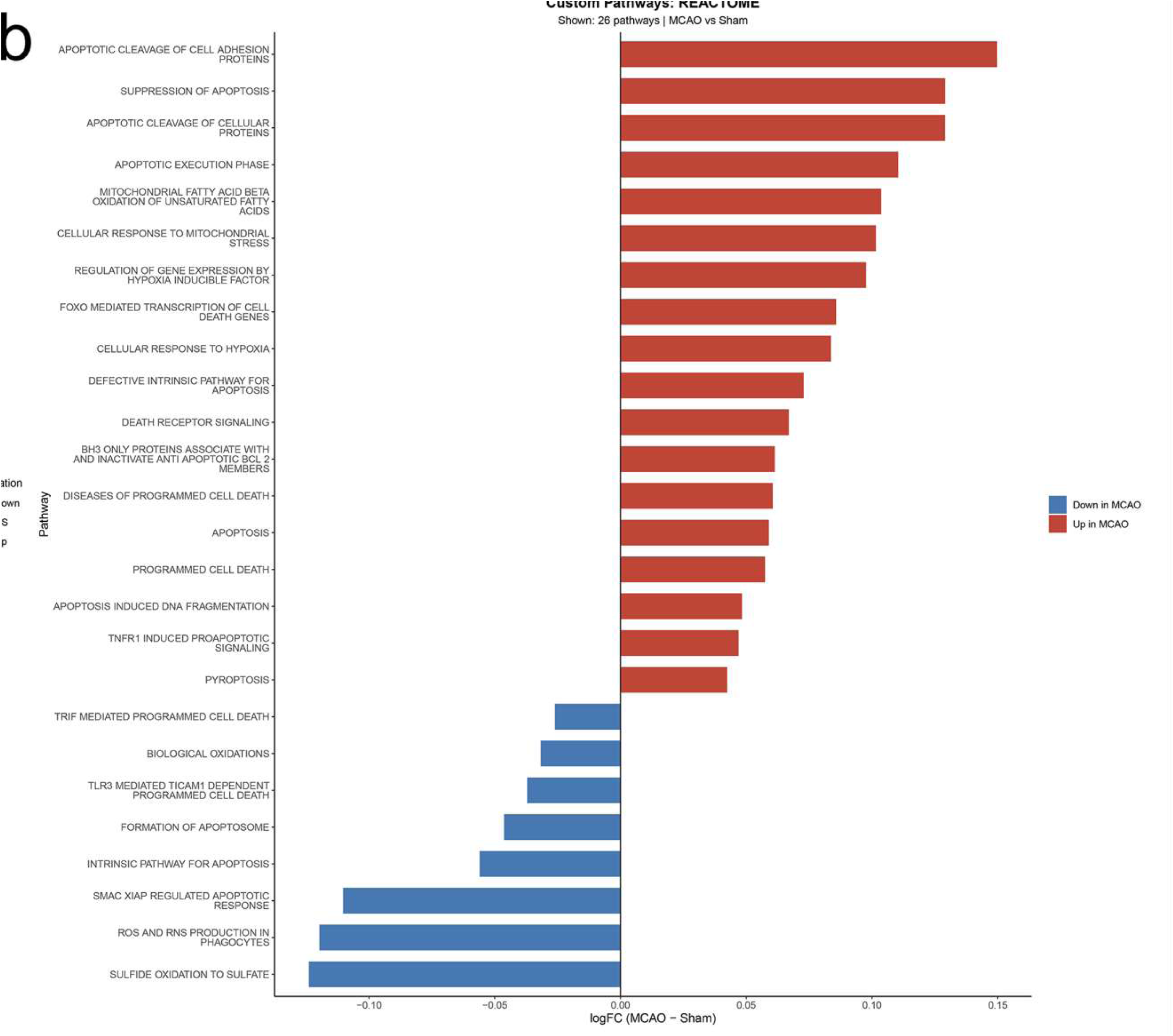
additionally displays volcano plots and bar plots of Reactome pathways from GSVA analysis (a: volcano plot of Reactome pathway; b: bar plot of Reactome pathway). This figure illustrates significant differences in Reactome pathways between Stressed Neurons and Surviving Neurons, revealing the activation and suppression of signaling pathways linked to neuronal apoptosis, iron metabolism, and other processes related to programmed cell death.

#### 3.2.4 Identification of ferroptosis-prone cells within the Stressed population

To isolate cells approaching the ferroptosis state within the Stressed neuron population, we extracted the gene expression matrix of Stressed neurons and applied TOGGLE for analysis.

TOGGLE partitioned the Stressed neurons into three distinct cell groups, encompassing a total of five states: Apoptosis (including Apoptosis-leaning and Apoptosis-prone states), Mix, and Ferroptosis (including Ferroptosis-leaning and Ferroptosis-prone states) (Figures 18 and 19).

**Figure 18.**
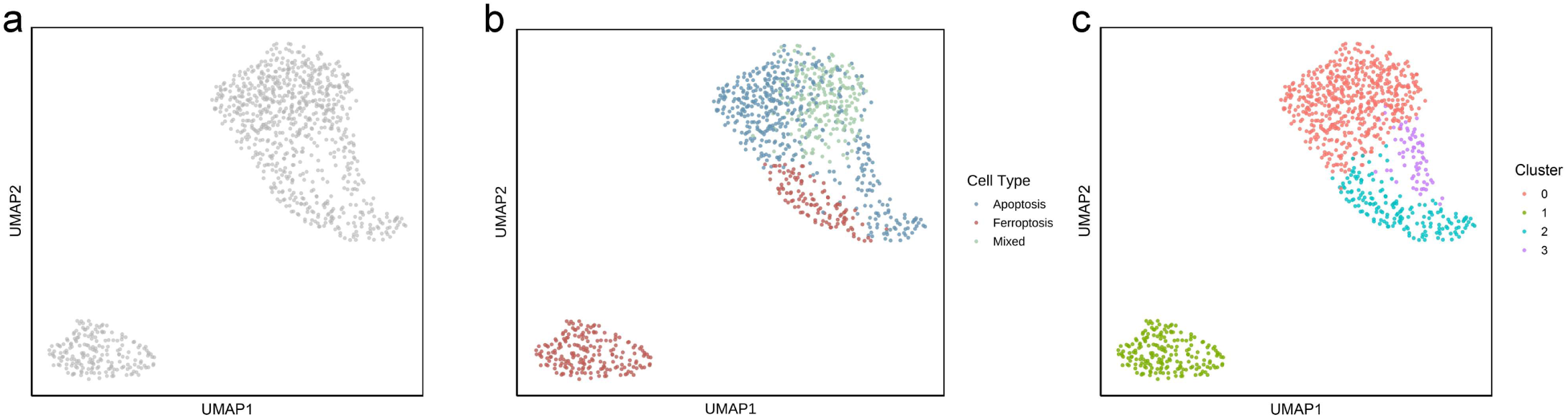
UMAP visualization of Stressed neurons (a: original UMAP; b: UMAP based on TOGGLE classification results; c: UMAP based on Seurat clustering). Seurat is one of the most widely used tools for cell classification, with its primary distinction levels based on prominent changes in gene expression patterns and differences in cell subtypes. This comparison is presented to highlight the differences between Seurat and TOGGLE, demonstrating that conventional classification levels cannot substitute for TOGGLE’s functional classification levels. The classification accuracy is validated in the sections that follow.

**Figure 19.**
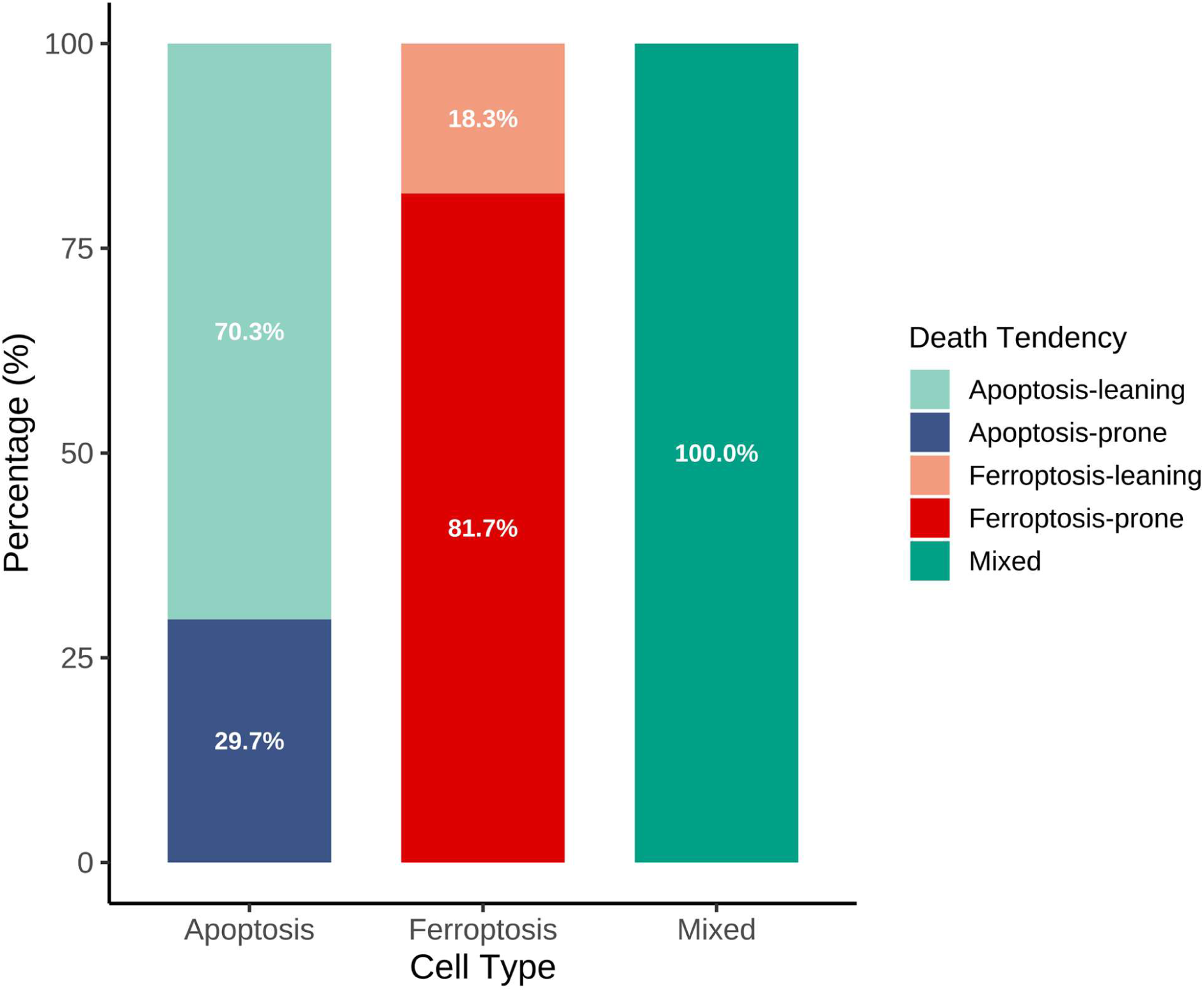
Death tendency in each cell group.

#### 3.2.5 Accuracy Inspection

In the Apoptosis group, 70.3% of the cells were in the Apoptosis-leaning state and 29.7% were in the Apoptosis-prone state. In the Ferroptosis group, 18.3% of the cells were Ferroptosis-leaning and 81.7% were Ferroptosis-prone.

#### 3.2.6 Biological interpretation of the ferroptosis grouping

GSVA analysis showed that, compared with Apoptosis cells, Ferroptosis cells exhibited greater activity in iron metabolism pathways closely associated with ferroptosis and in pathways that inhibit apoptosis, while pathways related to neuronal apoptosis were relatively suppressed (Figure 20).

**Figure 20.**
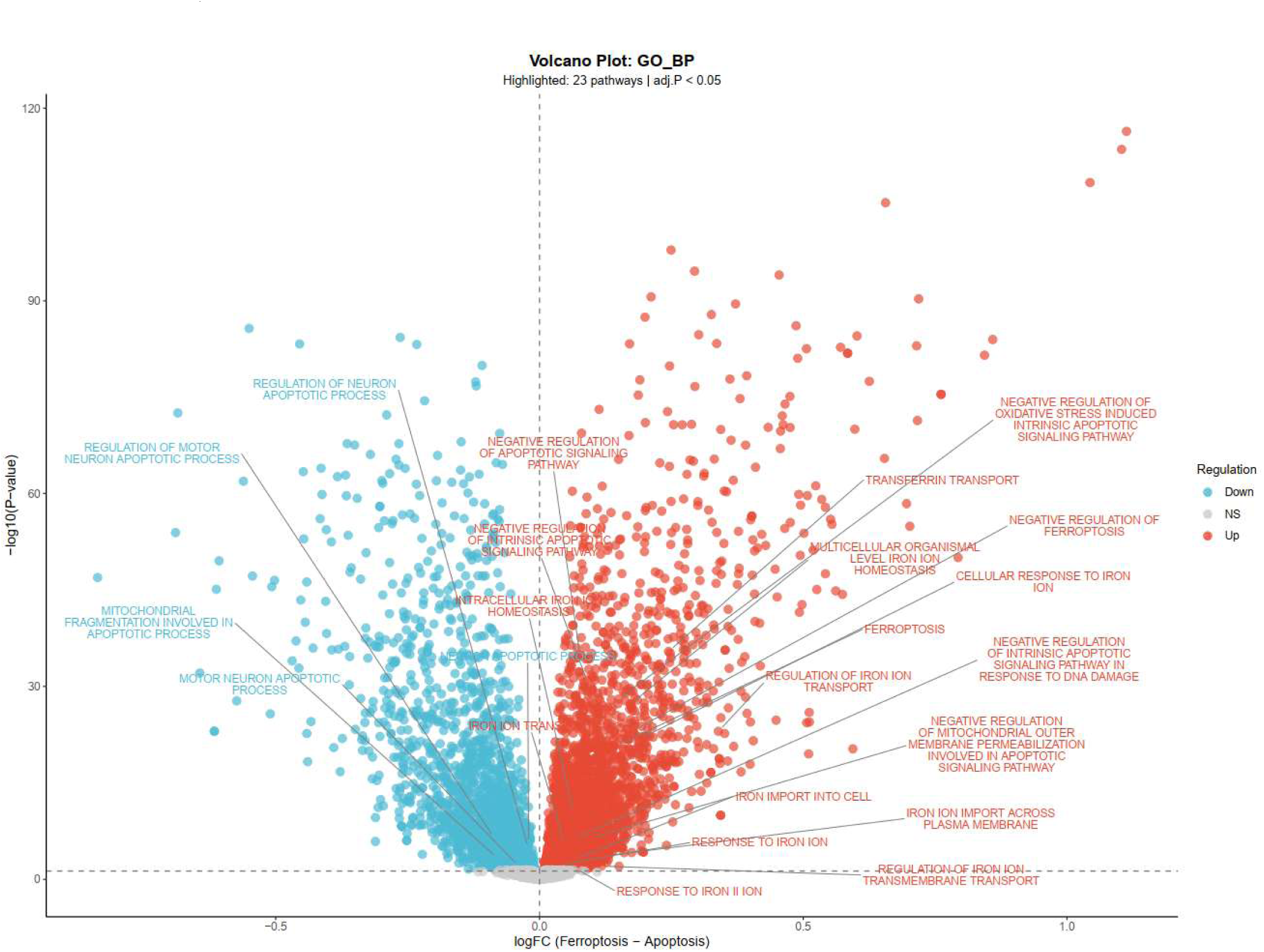
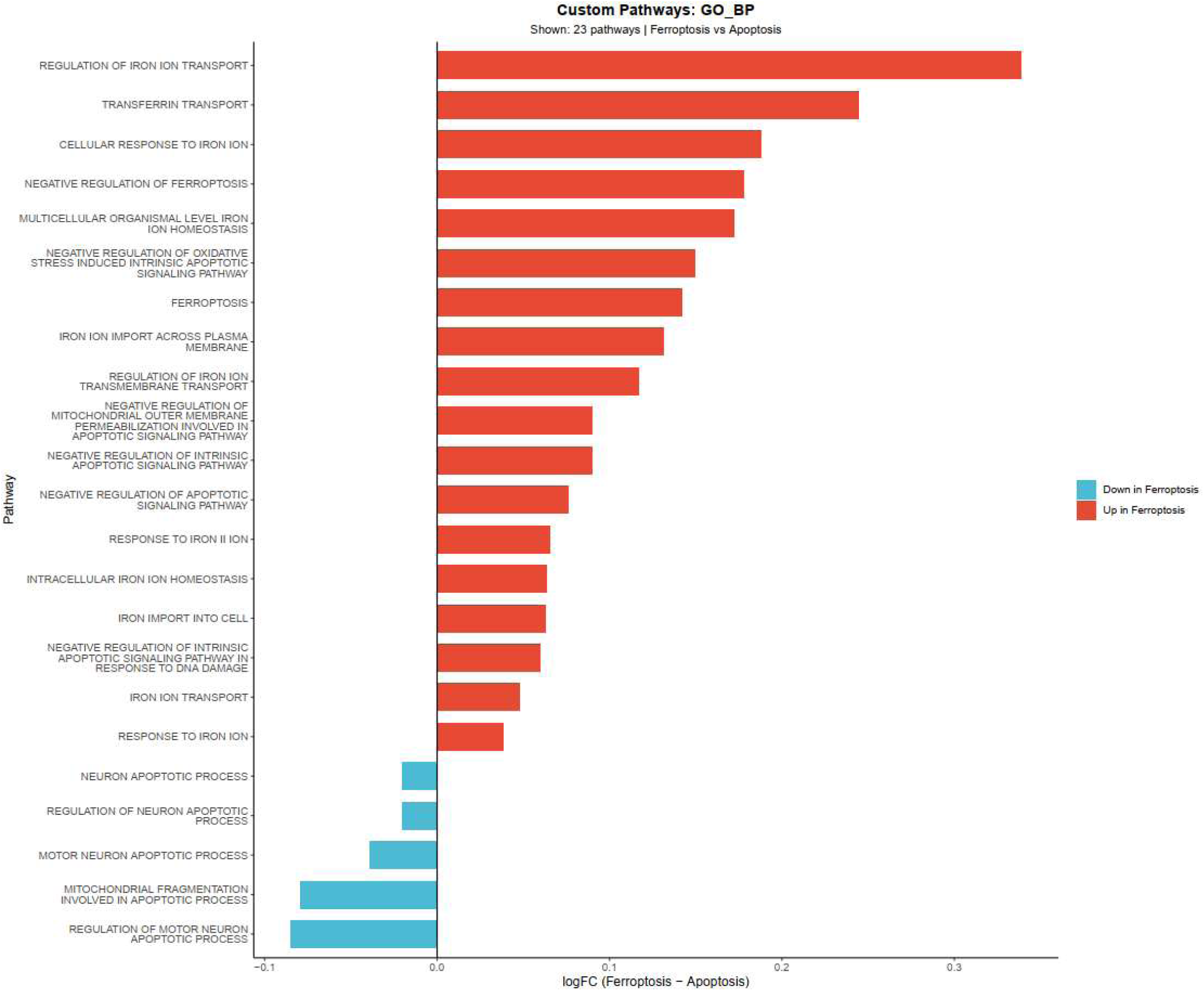
GSVA results comparing Ferroptosis and Apoptosis groups. The plot displays activated (red) and suppressed (blue) biological processes. In Ferroptosis cells, expression of apoptosis-related pathways is suppressed, whereas ferroptosis-related pathways show increased activity. These patterns confirm the success of the classification.

These findings indicate the successful classification by TOGGLE.

**Table 5.**
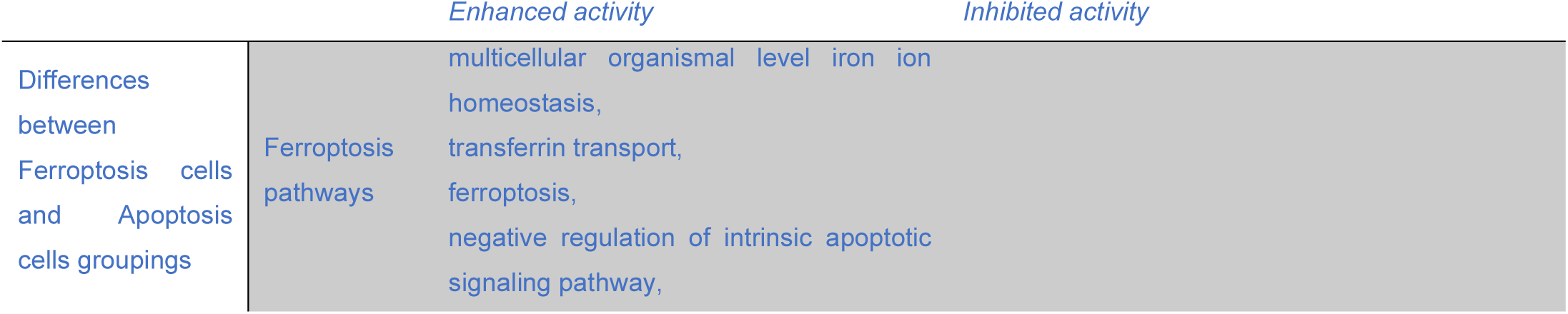
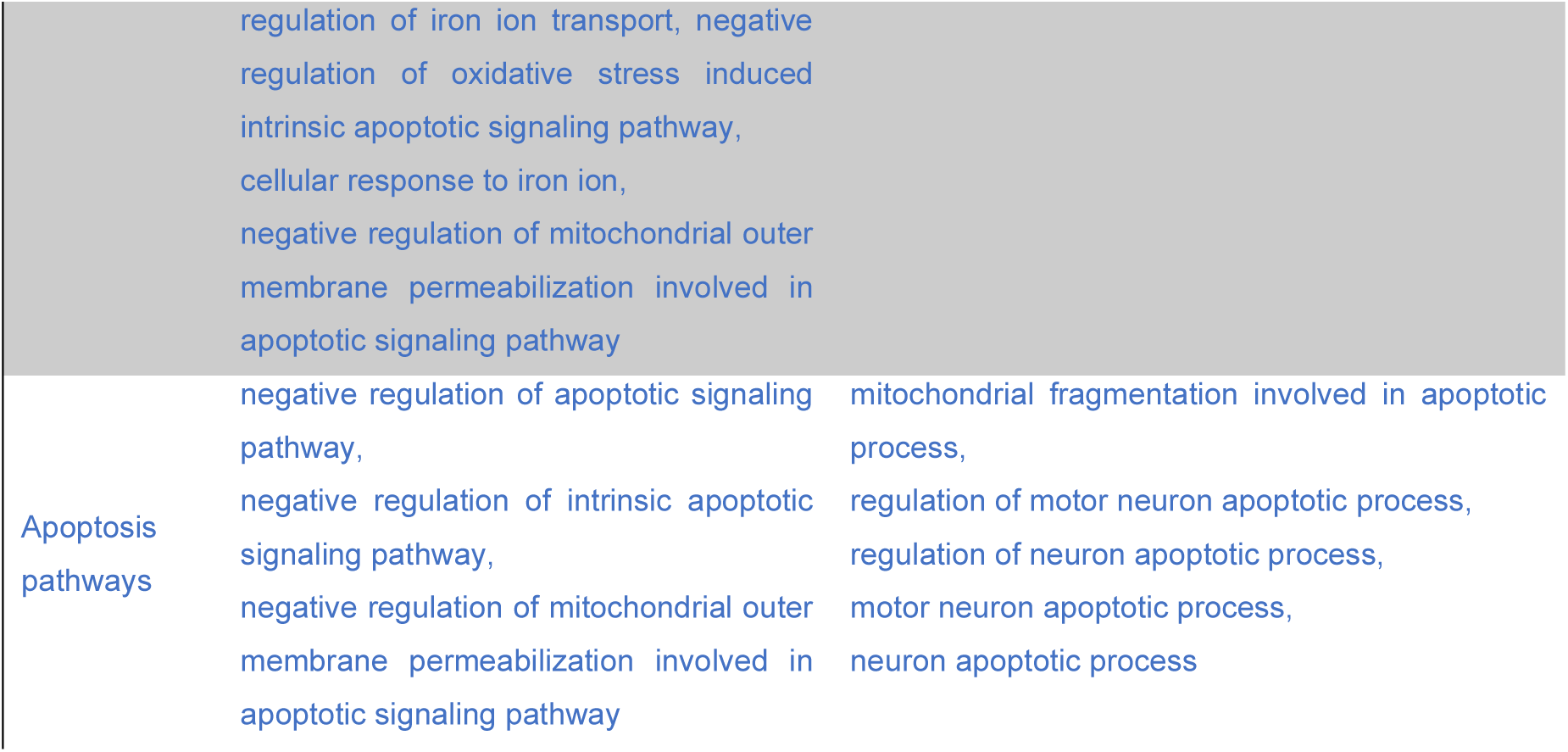
Differences between Ferroptosis cells and Apoptosis cells groupings.

#### 3.2.7 Ferroptosis- and Apoptosis-related genes expression in stressed neuron

To confirm the reliability of these findings, we examined the differentially expressed genes identified in section 3.2.1. For ferroptosis-related genes: (1) Pro-ferroptosis genes were upregulated, including Trf (Log2FC = 2.07, FDR < 0.05), Tfrc (also known as TFR; Log2FC = 0.60, FDR < 0.05), Smad1 (Log2FC = 1.05, FDR < 0.05), and Smad5 (Log2FC = 0.68, FDR < 0.05). (2) Anti-ferroptosis genes were downregulated, including Gpx4 (Log2FC = -1.04, FDR < 0.05).

Among apoptosis-related genes, pro-apoptotic genes were upregulated, including Rela (also known as p65; Log2FC = 1.00, FDR < 0.05), Bax (Log2FC = 0.77, FDR < 0.05), and Nfkbia (also known as IkBA; Log2FC = 1.89, FDR < 0.05). We subsequently performed experimental validation to further confirm the expression changes of these genes.

#### 3.2.8 Pathological changes of brain tissue

For methods of animal experiments, please refer to the Appendix 6.。In the Sham operation group, neurons exhibited normal morphology, with orderly arrangement, clear nuclear membranes, uniformly stained cytoplasm, and no evident pathological changes. In contrast, the MCAO group showed significant structural damage in the ischemic cortical area, with disorganized neuronal arrangement, widened intercellular spaces, and signs of cell swelling, rupture, and nuclear condensation, leading to vacuole formation. In the MCAO+deferiprone group, neuronal structural damage was mild, with minimal cell membrane rupture, slight nuclear condensation, and a significant reduction in vacuolar degeneration (Figure 21).

**Figure 21.**
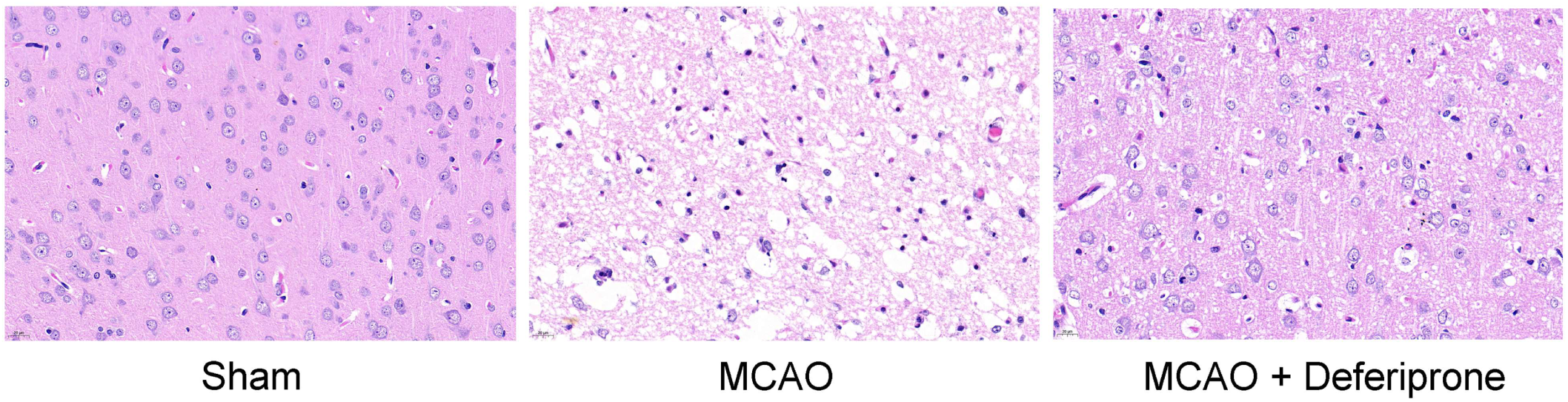
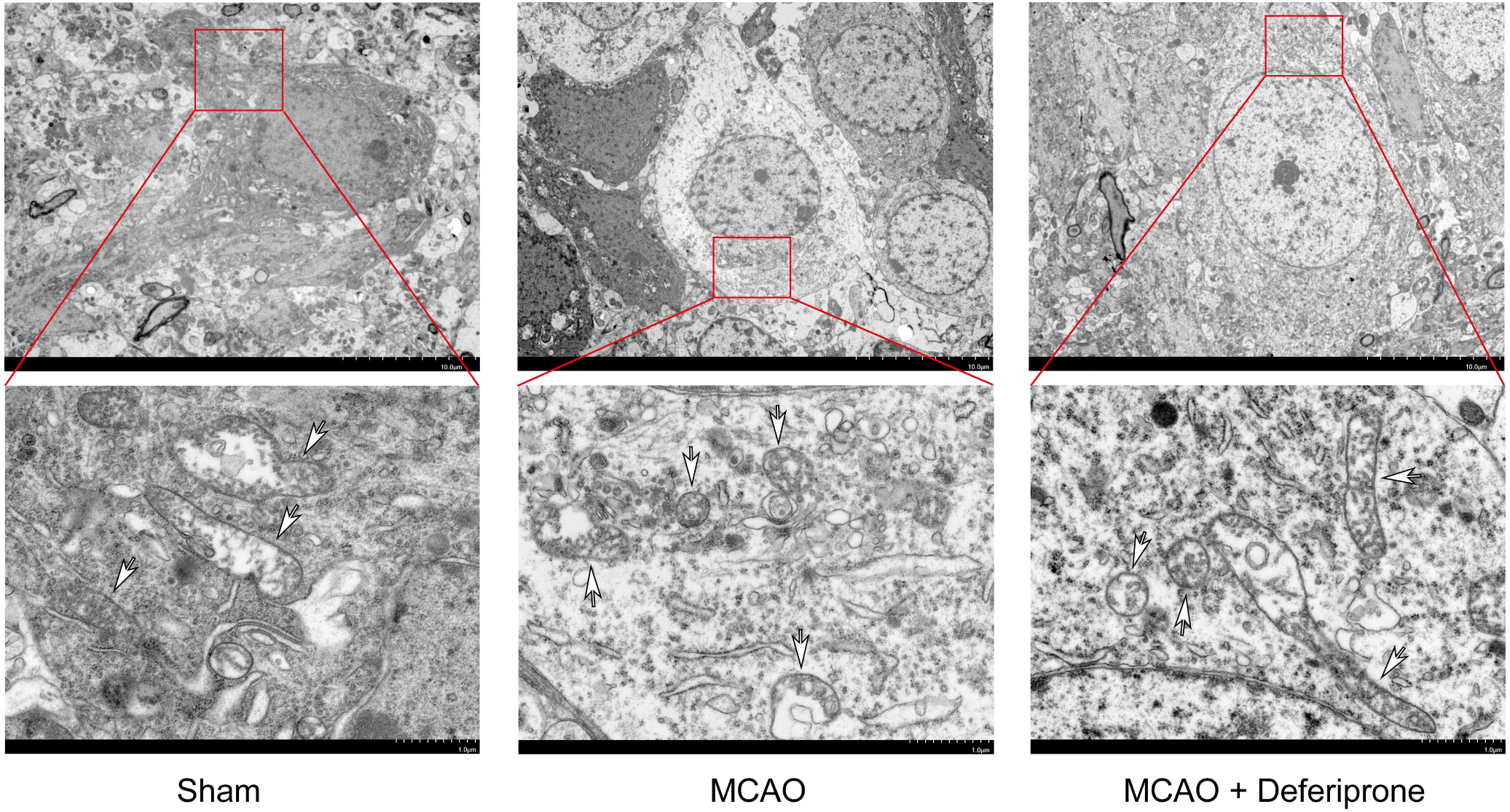
Pathological changes of brain tissue (Top). HE staining, 40× magnification. **Ultrastructural changes of brain tissue (Bottom).** Transmission electron microscope. The upper images are at 1500× magnification, and the lower images are at 6000× magnification. Mitochondria are marked by white arrows. **Summary:** Through these results, we observed characteristic cellular ultrastructural changes of ferroptosis.

#### 3.2.9 Ultrastructural changes of brain tissue

In the Sham group, mitochondria appeared structurally normal with clear features and intact membranes. In contrast, the MCAO group exhibited mitochondrial atrophy, reduced cristae, and increased membrane density, consistent with the morphological characteristics of ferroptosis. The MCAO+deferiprone group showed significant improvement in mitochondrial integrity compared to the MCAO group, with reduced damage (Figure21).

#### 3.2.10 Expression levels of *Acsl4, Gpx4* and *Fsp1* proteins detected by immunohistochemistry

Compared to the Sham operation group, the MCAO group showed an increase in Acsl4 levels (P<0.05) and a decrease in Gpx4 levels (P<0.05). These findings are consistent with the differential gene expression results from single-cell sequencing. Meanwhile, compared to the Sham operation group, the MCAO group showed an decrease in Fsp1 levels (P<0.05). Compared with the MCAO group, the MCAO+deferiprone group showed a decrease in Acsl4 levels and Fsp1 levels (P<0.05), and a increase in Gpx4 levels (P<0.05). (Figure 22).

**Figure 22.**
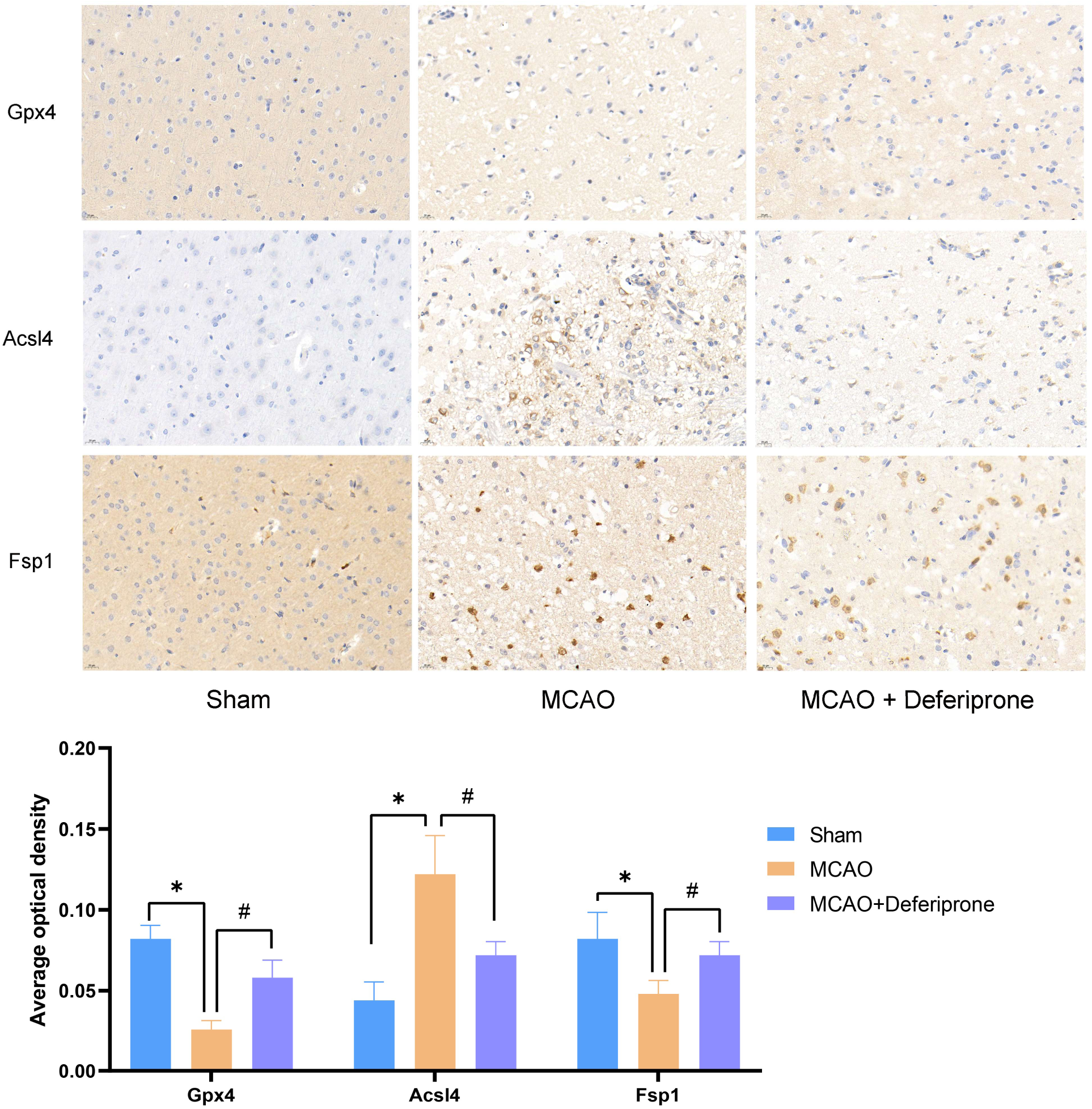
Expression levels of Acsl4 and Gpx4 proteins. Immunohistochemistry, 40× magnification. *compared with the sham group, P<0.05; #compared with MCAO group, #P<0.05. n=3 per group

#### 3.2.11 Expression levels of *Bax, IkBa, Casp3, p-Smad, p65(Rela), Tfrc, Trf and Sirt1* proteins detected by immunofluorescence staining

Compared to the Sham operation group, the MCAO group exhibited significantly elevated levels of Bax, IκBα, p-Smad, p65 (Rela), Tfrc, and Trf proteins (P<0.05). These results are consistent with the differential gene expression findings from the single-cell sequencing analysis. Meanwhile, Compared to the Sham operation group, the Casp3 protein level was increased, while the Sirt1 protein level was decreased in the MCAO group (P<0.05) (Figure 23).

**Figure 23.**
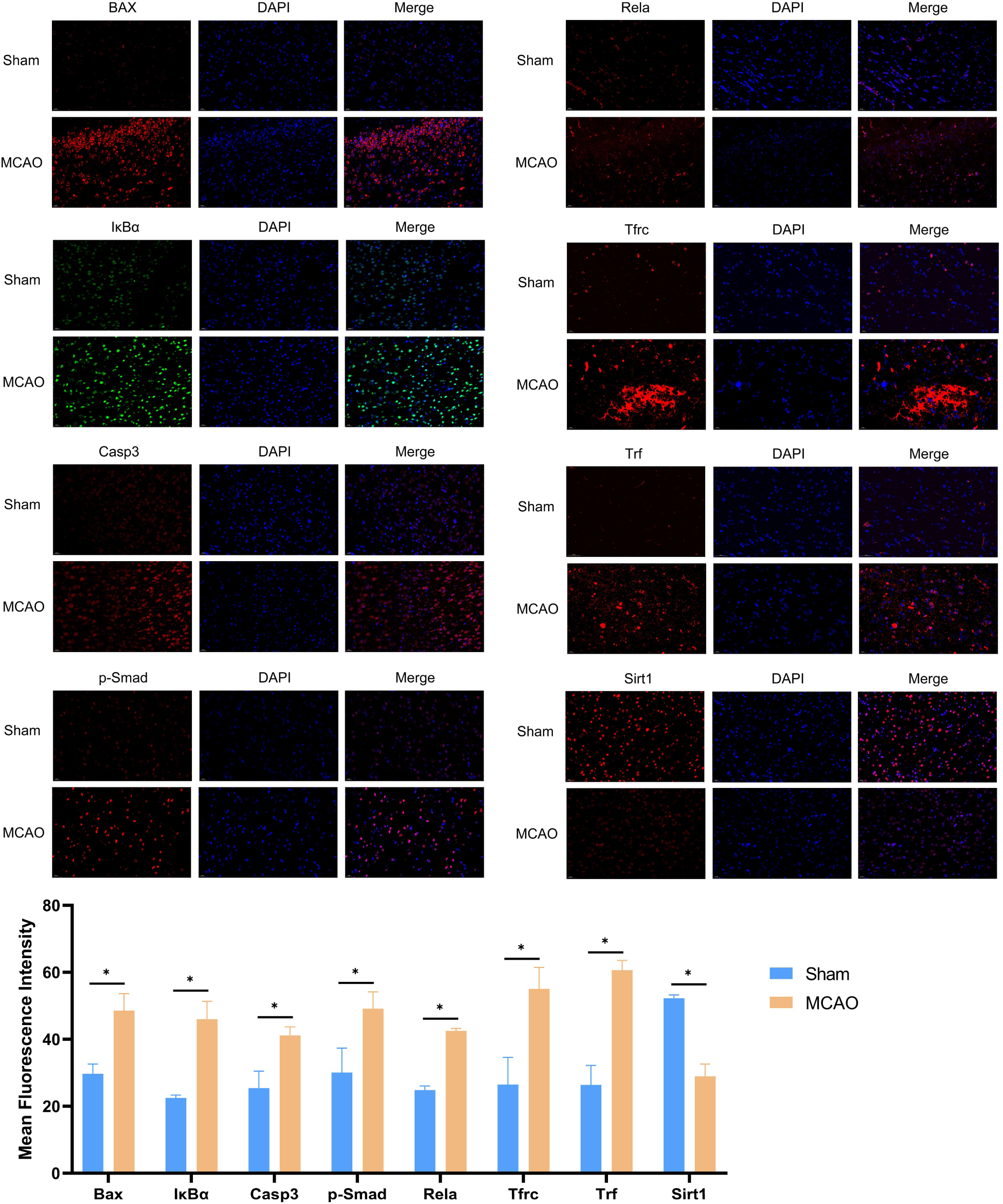
Expression levels of Bax, IkBa, p-Smad, p65, Tfrc, and Trf proteins. Immunofluorescence, 40× magnification. Compared with the sham group, *P<0.05. n=3 per group.

### 3.3 Test 3: Identifying Epigenetic Metabolic Regulation in NSCs Using Novel Functional Mapping

#### 3.3.1 Establish New RNA Functional Map (Graph Diffusion Functional Map)

We used Groups R3-4 and R3-5 from Section 3.2 to evaluate the effectiveness of RNA functional clustering. In the GSE209656 dataset (Figure 24), the top 1,000 most variable genes were selected from active neural stem cells (aNSCs) and quiescent neural stem cells (qNSCs). The results obtained using the method in Section 2.3 are shown in Figure 25 a-f and Table 6.

**Figure 24.**
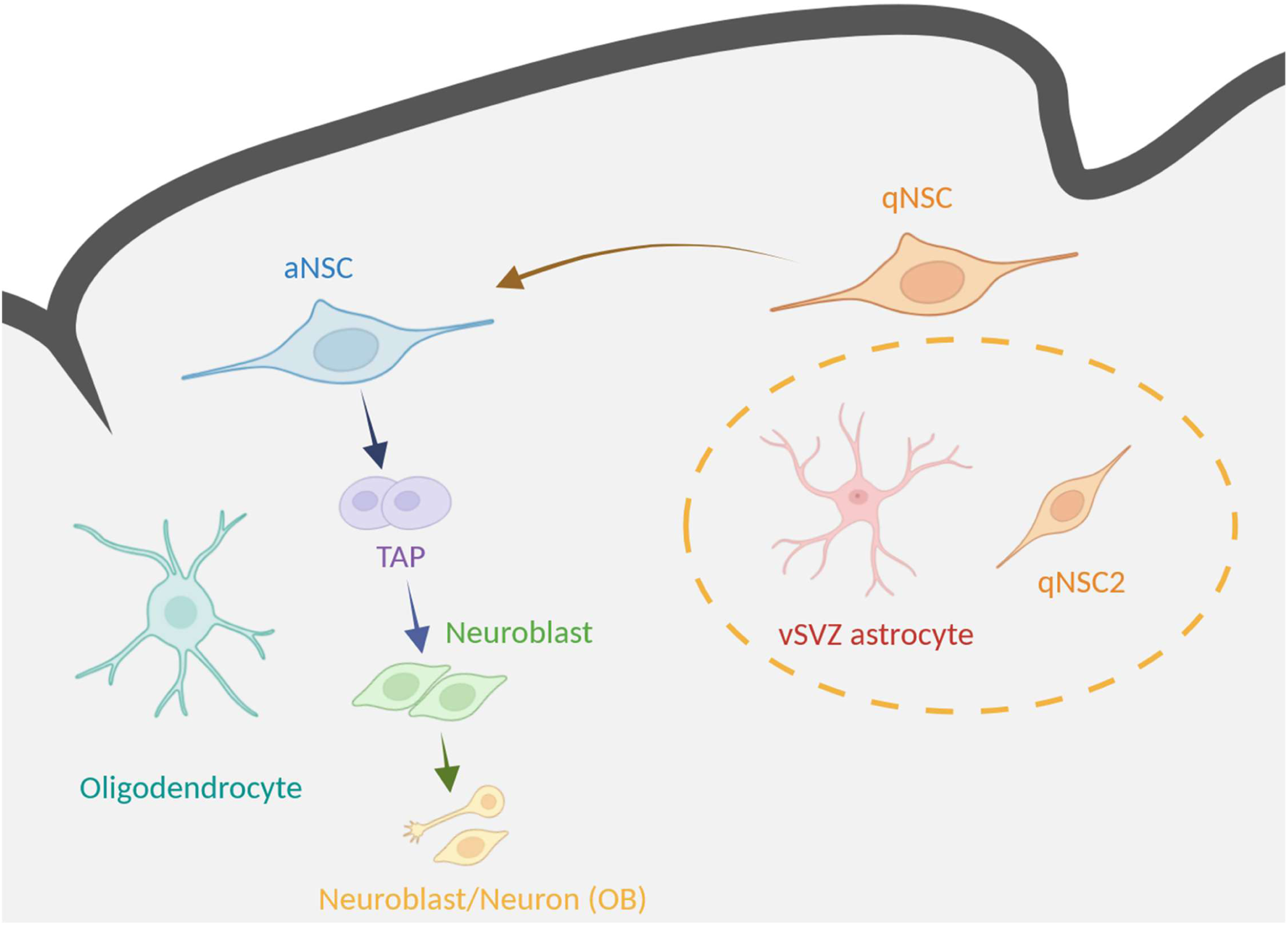
Brief background: Schematic diagram of the vSVZ adult NSC lineage. Referenced from Kremer et al. 2024^2^. qNSC: quiescent neural stem cell; qNSC2: Primed quiescent neural stem cells; TAP: transient amplifying progenitors; OB: olfactory bulb. qNSCs are classified into — qNSC2 and vSVZ. qNSCs can transition into aNSCs, and aNSCs can further differentiate into other types based on functional states. The complexity of these relationships renders most existing algorithms ineffective. Created by Biorender.

**Figure 25.**
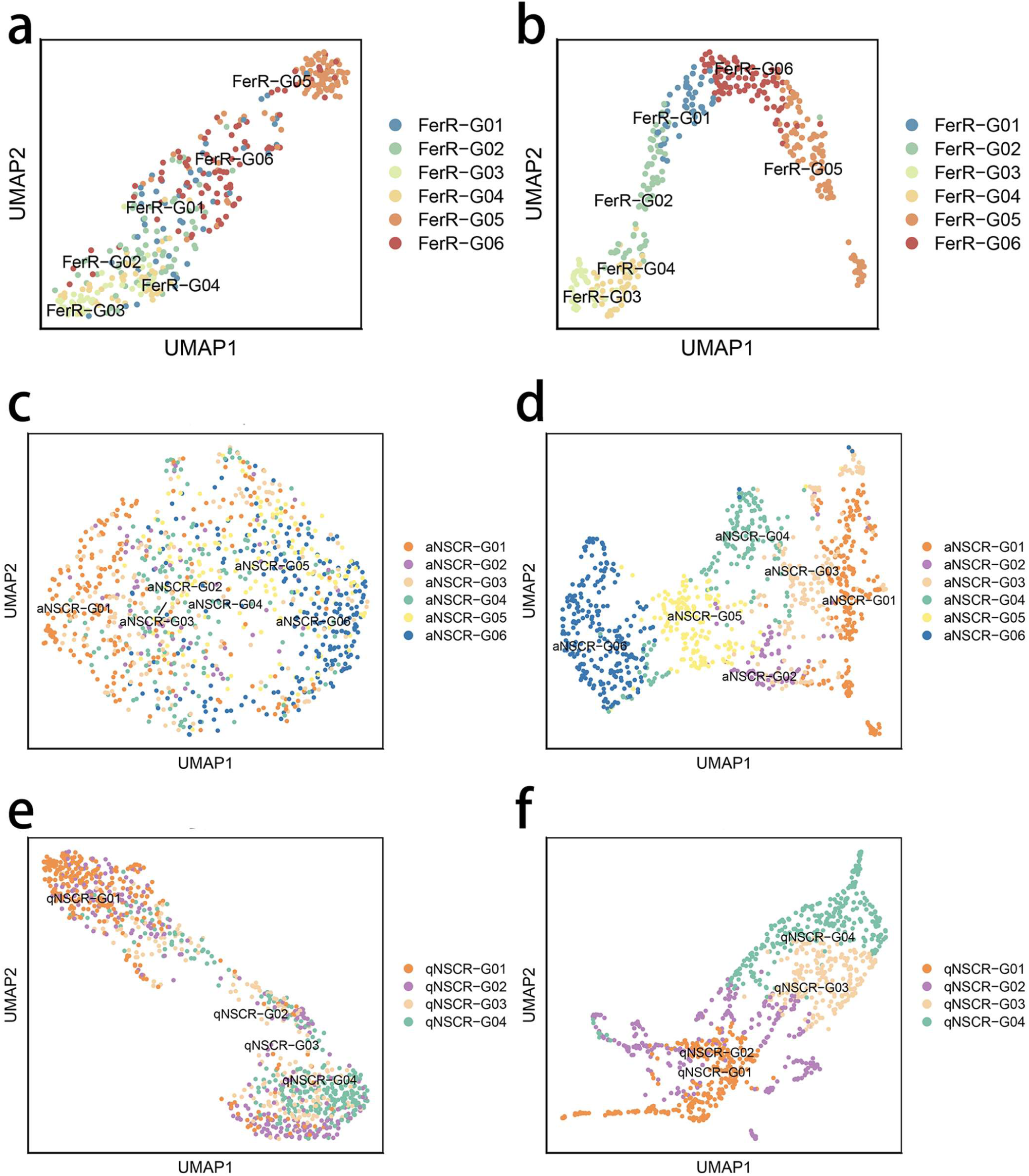
RNA functional profiles from different data. a: RNA map of ferroptosis-related genes in Group R3-4 and R3-5 cells made by using single-cell expression matrix; b: RNA map made using the new method for the data in a; c: RNA map of 1000 highly variable genes in aNSC cells made by using single-cell expression matrix; d: RNA map made using the new method for the data in c; e: RNA map of 1000 highly variable genes in qNSC cells made by using single-cell expression matrix; f: RNA map made using the new method for the data in e.

**Figure 26.**
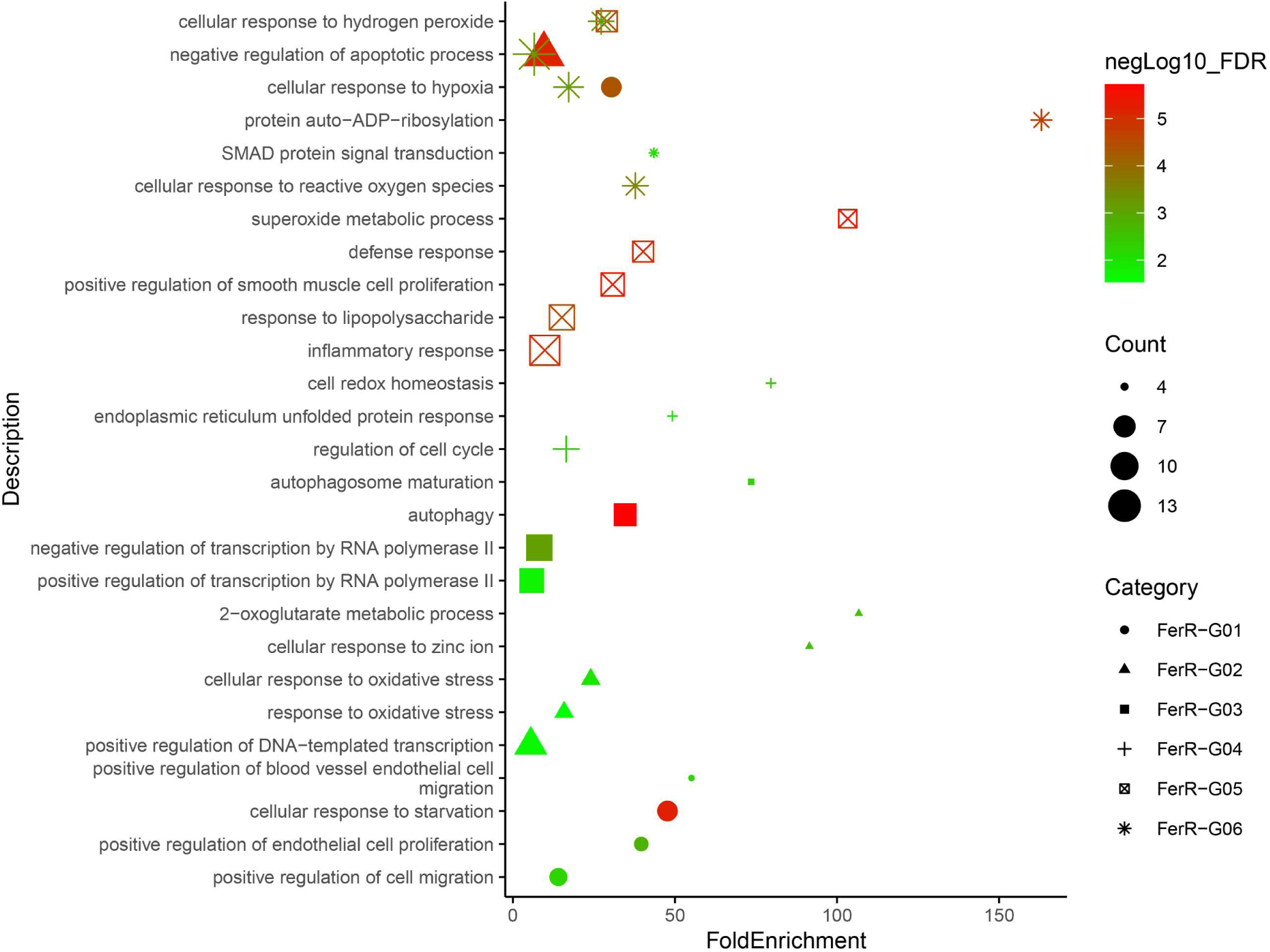
Bubble chart: The function of each RNA group in Figure 11(B), as determined by enrichment analysis. Genes not enriched in specific signaling pathways may still be functionally associated with their corresponding RNA groups, potentially revealing novel mechanisms involved in ferroptosis.

**Table 6:**
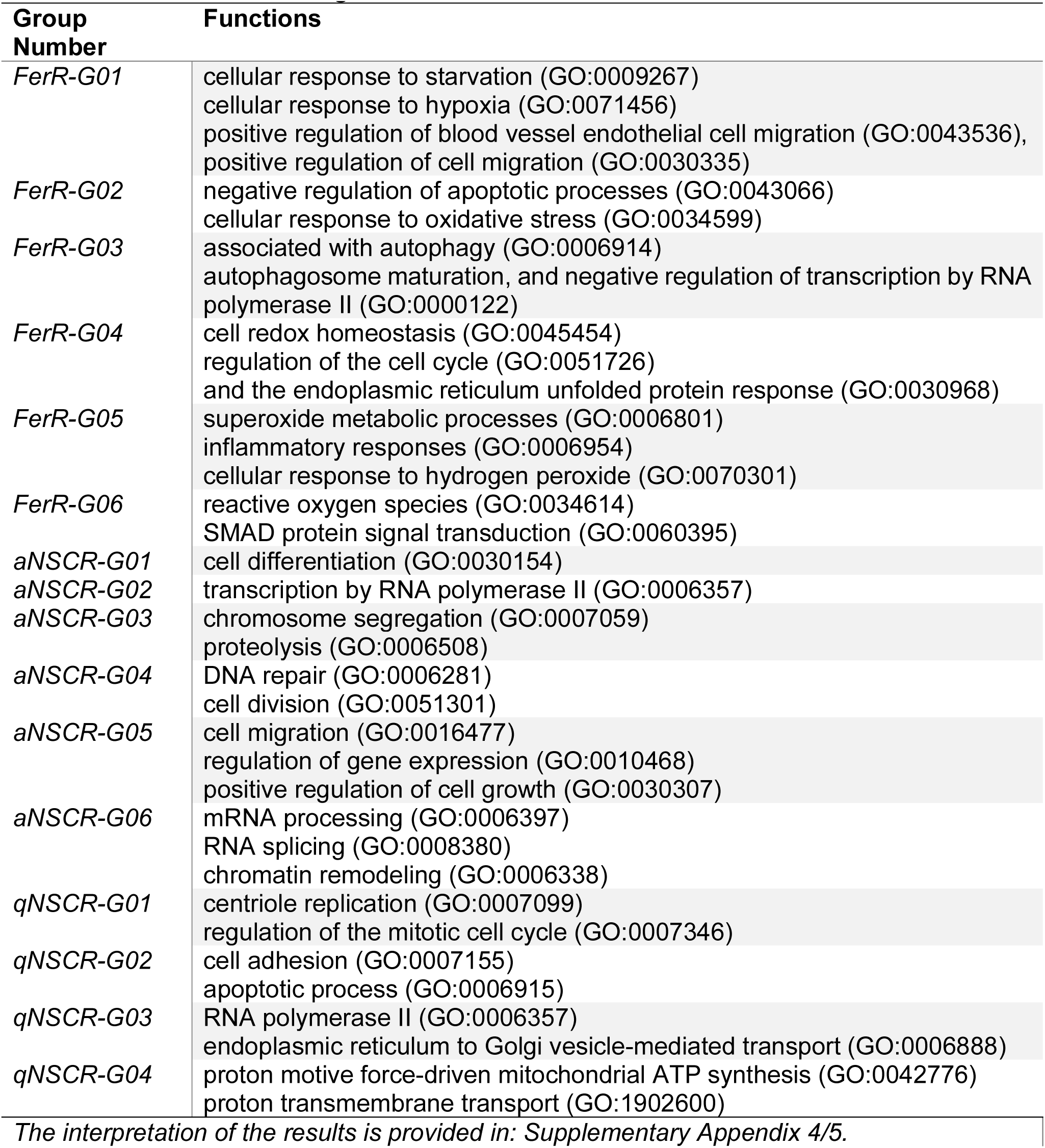
RNA Functions of figure 11a-f.

We performed functional enrichment analysis on RNA clusters derived from ferroptosis-related cell populations (FerR-G01 to G06) and neural stem cells (aNSCs and qNSCs), focusing on highly variable genes. Taking ferroptosis as an example (Figure 12), FerR-G01 was enriched in hypoxia response and pro-angiogenic functions. These enriched pathways suggest that this group may help alleviate hypoxia in ischemic brain regions by promoting neovascularization. Key genes such as Nfs1, Osbpl9, and Pgd may regulate iron metabolism, lipid signaling, and antioxidant capacity. Genes not enriched in specific pathways may still be functionally associated with their RNA group, potentially revealing novel mechanisms of ferroptosis. For example, in FerR-G01, Nfs1 plays a critical role in the biosynthesis of iron–sulfur (Fe–S) clusters, which are essential for mitochondrial respiratory chain complexes. Hypoxia caused by cerebral ischemia can impair mitochondrial function, reduce ATP production, and lead to cellular injury. Nfs1 may regulate ferroptosis by maintaining Fe–S cluster homeostasis, thereby influencing iron metabolism and antioxidant defense. Osbpl9 is a key regulator of cholesterol and phospholipid exchange, involved in intracellular lipid transport and signal transduction. It may promote endothelial cell migration and angiogenesis in ischemic regions by modulating the lipid microenvironment and associated signaling pathways. Pgd participates in the pentose phosphate pathway (PPP) and primarily functions by generating NADPH to maintain cellular antioxidant capacity. Under hypoxic conditions, elevated reactive oxygen species (ROS) levels induce oxidative stress. Pgd may counteract ferroptosis by boosting antioxidant defenses, protecting cells from oxidative damage.

#### 3.3.2 Identifying Epigenetic Metabolic Regulation in NSCs Using Novel Functional Mapping

Neural stemness refers to the ability of neural cells to self-renew and differentiate into other cell types. This state is not static but continuously evolving, which blurs the functional boundaries between different neuronal states and makes accurate classification of these states extremely challenging.

Kremer et al^2^. demonstrated that despite their distinct functional roles, astrocytes and quiescent neural stem cells (NSCs) exhibit highly similar transcriptomic profiles^2^. However, studies have shown that under specific conditions—such as injury or overexpression of key factors—astrocytes can reactivate their latent neurogenic potential. This discovery opens new avenues for regenerative medicine: by regulating cellular memory and energy metabolism, it may be possible to induce astrocytes to acquire neurogenic capacity, enabling regenerated tissues or organs to exhibit neuronal activity. Cellular memory involves multiple components, including protein, RNA, and DNA modifications, but the precise mechanisms sustaining cellular energy activity remain unclear. Improving cellular energy metabolism—such as enhancing mitochondrial function or supplying metabolic substrates—can significantly inhibit myofibroblast activation. This strategy holds promise for reversing heart failure by promoting the transdifferentiation of myofibroblasts into fibroblasts, thereby restoring cardiac function^34^.

To investigate the epigenetic regulation mechanisms of energy metabolism, we conducted a multimodal analysis of the single-cell RNA sequencing (scRNA-seq) and single-cell epigenomic data of NSCs provided by Kremer et al.² First, active neural stem cells (aNSCs) and quiescent neural stem cells type 2 (qNSC2) were extracted based on labels provided by Kremer et al.², and the scRNA-seq dataset (GSE209656) was analyzed using TOGGLE (Figure 27).

**Figure 27.**
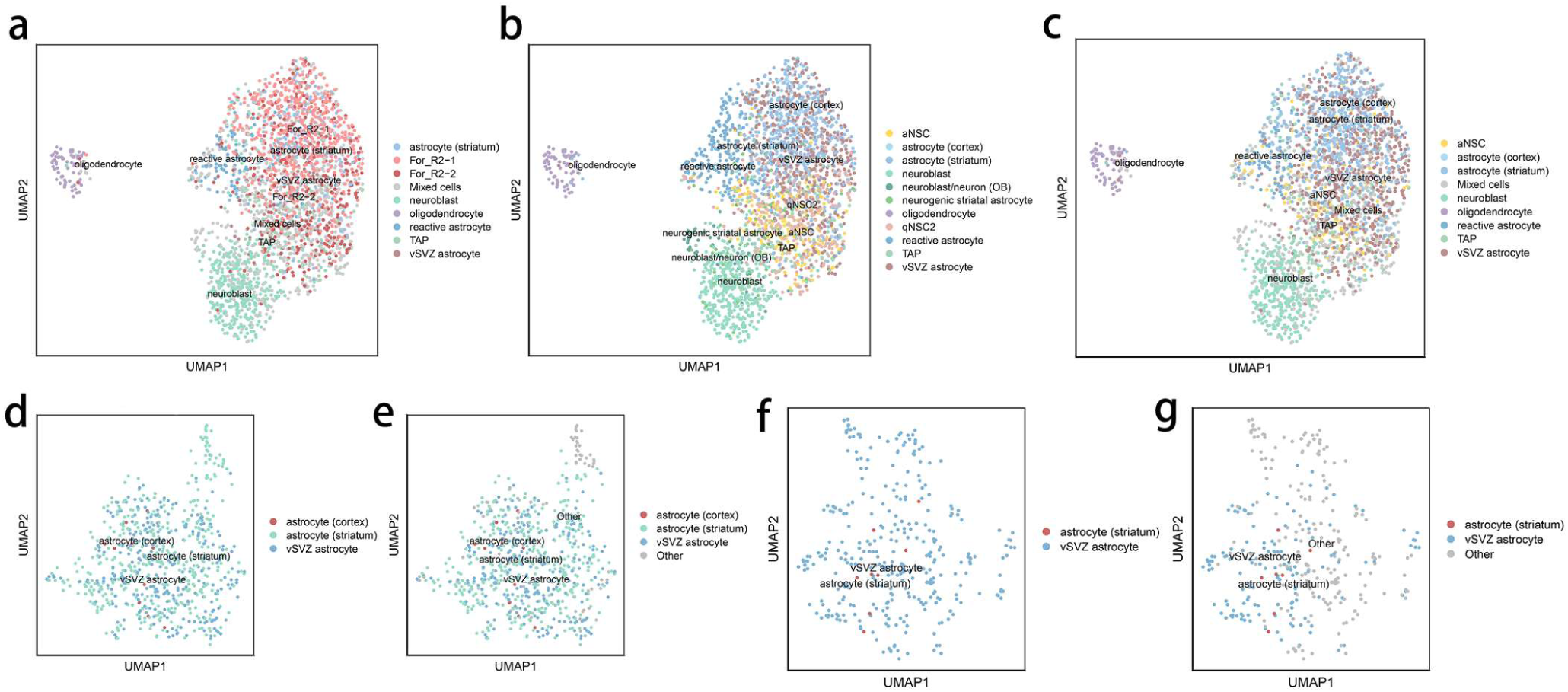
A: Preliminary prediction results indicating that For_R2-1 and For_R2-2 are highly similar and difficult-to-classify groups. B: True labels. C: Results after completing classifications D and F. D: Predicted labels for For_R2-1. E: True labels for For_R2-1. F: Predicted labels for For_R2-2. G: True labels for For_R2-2.

Next, we divided the cells (Figure 28a and 28b) into two groups based on the median mitochondrial RNA expression levels: a high mitochondrial RNA expression group (High-MT) and a low mitochondrial RNA expression group (Low-MT) (Figure 28d and 28e). The analysis revealed that NeuG04 predominantly consisted of metabolically active cells, mainly active neural stem cells (aNSCs), whereas NeuG01 was mainly composed of metabolically inactive cells, primarily quiescent neural stem cells (qNSCs) (Figure 28c and 28f). The remaining cell groups were distributed between these two categories, possibly representing transitional metabolic states. To identify differentially methylated regions between the two groups, we performed two-sided Wilcoxon rank-sum tests on VMRs covered by at least six cells in each group. Regions with a Benjamini–Hochberg corrected P-value less than 0.05 were labeled as low methylation regions (LMRs) or highly accessible chromatin regions (high ACC). LMRs overlapping gene body regions were annotated to their corresponding genes.

**Figure 28.**
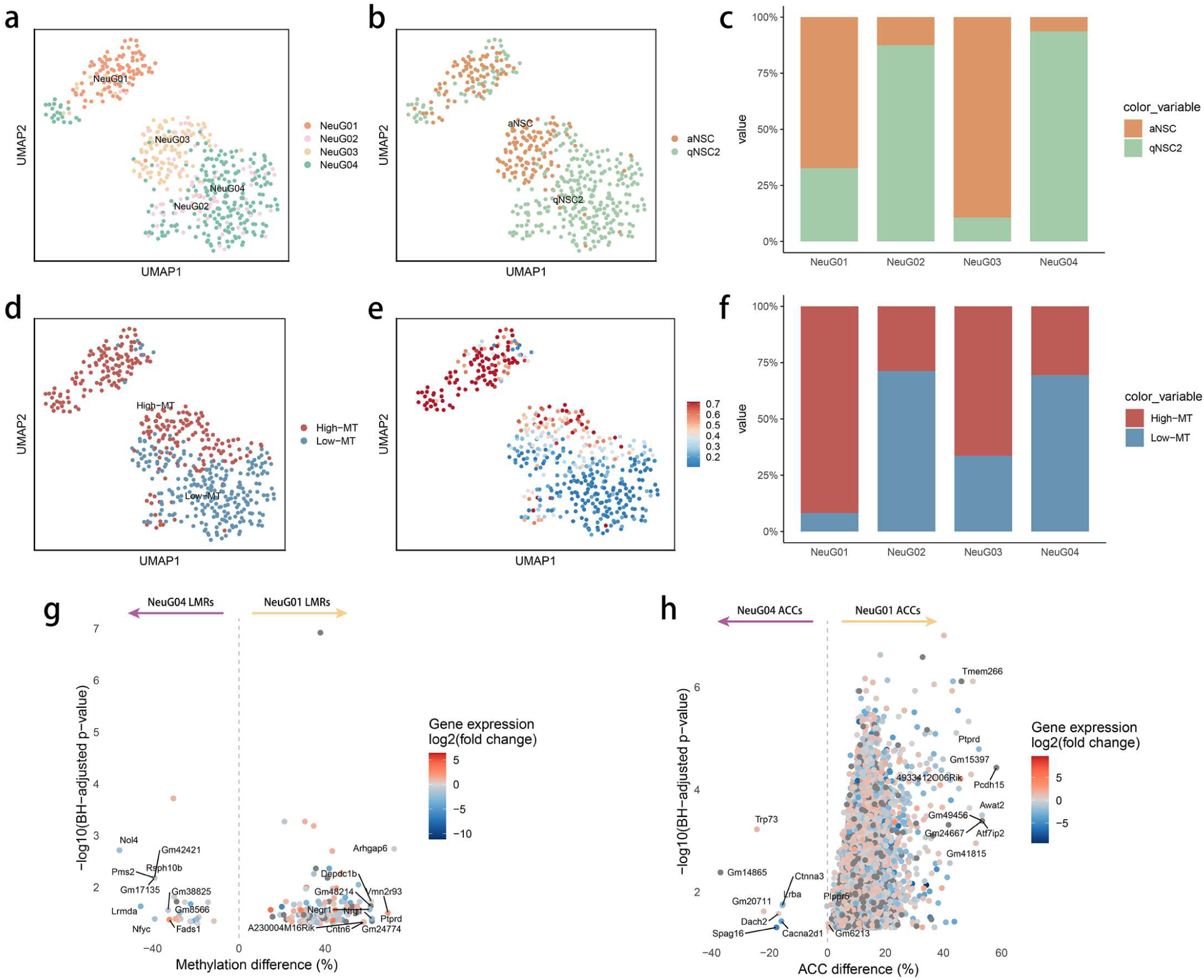
Results of epigenetic analysis of aNSCs and qNSCs. a: Cell populations after TOGGLE analysis; b: Labels provided by Kremer et al.; c: The proportion of aNSC and qNSC2 in each cell group; d: Cells are divided into High-MT and Low-MT according to the expression of mitochondrial genes; e: Mitochondrial gene expression in cells; f: The proportion of high-MT cells and low-MT cells in each cell group; g: Volcano plot of LMRs, tested for differential methylation between NeuG01 and NeuG04 (two-sided Wilcoxon test), LMRs are colored according to the differential expression of the nearest gene. h: Volcano plot of high ACC resgions, tested for differential ACC between NeuG01 and NeuG04 (two-sided Wilcoxon test), LMRs are colored according to the differential expression of the nearest gene

Since NeuG01 predominantly consists of aNSC cells and NeuG04 mainly comprises qNSC2 cells, we conclude the following: (1) During the transition from qNSC2 to aNSC, local DNA demethylation occurs in key genomic regions associated with cell proliferation and energy metabolism (Figure 28g), accompanied by significantly increased chromatin accessibility (Figure 28h). (2) The proportion of cells exhibiting high mitochondrial gene expression (High-MT) within the aNSC-dominated group is significantly higher than in the qNSC2 group (Figure 28c, 28d, and 28e), further indicating enhanced metabolic activity in aNSCs. In summary, we propose that during the transition from qNSC2 to aNSC, local DNA demethylation and chromatin opening activate the expression of genes related to cell proliferation and energy metabolism, gradually enhancing metabolic activity. These epigenetic modifications reflect a difference in “cellular memory” between the two cell types, potentially serving as the underlying mechanism driving their distinct mitochondrial metabolic states.

### 3.4 Other test results and findings

In this study, we applied the TOGGLE algorithm to the functional classification of cell-cell communication^35–39^ and RNA pathways. Due to space constraints, detailed methodological descriptions are provided in the appendix, and only a summary is presented here. In the e-cigarette aerosol exposure model (GSE183614), TOGGLE grouped differentially expressed genes and conducted enrichment analyses, identifying that Groups 2 through 5 exhibited significant immune and inflammatory responses across biological processes and KEGG pathways (e.g., TNF signaling and cell cycle). This effect was particularly pronounced in younger mice, indicating that e-cigarette exposure has more severe impacts on immature cardiac tissue. In the myocardial infarction (MI) model (GSE253768), TOGGLE integrated single-nucleus RNA sequencing (snRNA-seq) data with bulk RNA sequencing data to identify seven functional subpopulations of fibroblasts and myofibroblasts (FibR1-G1 to G7), with the MI group predominantly composed of FibR1-G5 through G7. Further analysis of cell-cell ligand-receptor interactions revealed differences in their communication patterns with immune cells. TOGGLE subsequently mapped bulk RNA-seq data onto the snRNA-seq space, discovering that the bulk RNA-seq signals primarily originated from FibR2-G6, a population enriched in pathways related to cell migration, adhesion, angiogenesis, and ECM-receptor interactions, suggesting its role as a key cell type driving tissue-level changes. In summary, TOGGLE not only precisely dissects cellular heterogeneity but also integrates single-nucleus and bulk RNA sequencing data across platforms to uncover critical signaling pathways and functional cellular populations. These results are detailed in the Supplementary Appendix 1&2.

## 4. Discussion

This study proposes TOGGLE, an unsupervised learning framework that does not require manual labeling. Combining graph diffusion, self-supervised mechanisms, and a BERT-like deep encoding network, TOGGLE achieves high-precision identification of cell functional states and fate stages from single-cell transcriptomic data. Additionally, TOGGLE supports integrated analysis of single-cell transcriptomic and epigenomic datasets. In neural stem cell (NSC) data, cells were divided into High-MT and Low-MT groups based on mitochondrial RNA expression levels. The analysis revealed that the transition from qNSC to aNSC is accompanied by increased chromatin accessibility and DNA demethylation, elucidating the epigenetic memory mechanism underlying metabolic activity differences. Compared to mainstream algorithms, TOGGLE has advantages in distinguishing cell functional states, tracking cell fate trajectories, integrating cross-modal data, eliminating the need for pre-training, and enabling multifunctional analyses within similar cell populations (see Table 7 for details). It thus holds broad applicability in regenerative medicine, immunotherapy, responses to viral infections, and cancer research.

**Table 7:** Research differences.

| Algorithms | Subtype | Development |  | Communication | Death | Functional RNA | Functional regulation distinction |  |  |
| --- | --- | --- | --- | --- | --- | --- | --- | --- | --- |
|  | Cell subtype differentiation | Developmental prediction | Distinguishing between progenitor and daughter cells | Functional classification of intercellular communication | Label-free identification of cells with promiscuous shared pathways | RNA functional cluster classification | Epigenetic memory distinction, taking methylation as an example | RNA transcription activity changes in epigenetic memory | Differentiate between the developmental stages of the same cell |
| Cell Rank | √ | × | √ | × | × | × | Part*1 | Part*1 | × |
| Deep Cell | √ | × | √ | × | Part*2 | × | × | × | × |
| scGen | √ | × | × | Part*1 | × | × | × | × | × |
| CellTypist | √ | × | √ | Part*1 | × | × | × | × | × |
| SCENIC | √ | × | × | × | × | Part*1 | × | × | × |
| pySCENIT | × | × | × | Part*4 | × | Part*4 | × | × | × |
| WOT | √ | √ | √ | × | × | × | × | Part*3 | × |
| Cospar | × | √ | × | × | × | × | × | × | × |
| MARGARET | √ | × | × | × | × | × | × | × | × |
| TOGGLE | √ | √ | √ | √ | √ | √ | √ | √ | √ |
| Pseudo-time type algorithm | × | × | × | × | × | × | × | × | × |

Part\*1 is limited to distinguishing between different cell types; Part\*2 can only identify cells that have undergone terminal cell death; Part\*3 requires a fully comprehensive reference dataset; Part\*4 is restricted to analyzing regulatory networks.

While TOGGLE demonstrates robust performance in integrating transcriptomic and epigenomic data and tracing cell fate transitions, it also has several limitations. Firstly, it does not yet incorporate post-transcriptional regulatory signals such as RNA methylation (e.g., m6A), which play critical roles in modulating gene expression. Secondly, the algorithm’s ability to resolve rare or minor cell subpopulations remains to be further improved, as these subtypes often carry important biological functions in complex tissues. Lastly, TOGGLE currently does not include the analysis of post-translational protein modifications (e.g., phosphorylation, ubiquitination), which are key regulators of signaling and cell functional states. Future work could aim to integrate these additional regulatory layers to provide a more comprehensive picture of cell state and functionality.

## Contribution

The methodology, bioanalysis, and test application of this paper were collaboratively designed by Junpeng Chen, Zhouweiyu Chen, Kailin Yang. Junpeng Chen organized and led the entire study, defining the manuscript’s topic and narrative, designed the experimental protocol and algorithmic framework, and wrote the entire manuscript. Junpeng Chen was the main contributors to the algorithm innovation, and the primary completer and contributor of this manuscript and project-code, while Kailin Yang was the main contributors to the bioanalysis and bio-innovation. Zhouweiyu Chen assisted in solving critical computational problems and completed the design and testing of auxiliary plans, making the method complete. Junpeng Chen proposed a complete set of biological problems and performed the analysis, with assistance from Kailin Yang, Kaiqing Liu, Tianda Sun, Yibing Nong, Tao Yuan, Charles C. Dai, Changping Li, Yexing Yan, Zixiang Su, You Li, Tian Song and Richa A. Singhal, etc. Jinwen Ge, Haihui Wu, Tong Yang, Shanshan Wang and Kailin Yang performed the animal experiments. Alex P. Carll played a crucial role in correspondence, providing experimental resources and research funding. Junpeng Chen, Alex P. Carll, and Lu Cai jointly organized the study. Alex P. Carll, Yibing Nong guided the analysis of cardiac pathway differences and cardiomyocytes. Lu Cai supervised the experimental work on ferroptosis.

## Acknowledgments

The University of Louisville provided computational equipment support for this research and assisted all researchers with equipment maintenance and updates. These hardworking and dedicated staff members share this honor with us, including the following staff: Harrison Simrall, Tomas Felipe Llano Rios, and Randy Vargas. The content is solely the responsibility of the authors and does not necessarily represent the official views of the NIH or FDA. 𝘛𝘩𝘦 𝘜𝘯𝘪𝘷𝘦𝘳𝘴𝘪𝘵𝘺 𝘰𝘧 𝘓𝘰𝘶𝘪𝘴𝘷𝘪𝘭𝘭𝘦 𝘱𝘳𝘰𝘷𝘪𝘥𝘦𝘥 𝘤𝘰𝘮𝘱𝘶𝘵𝘢𝘵𝘪𝘰𝘯𝘢𝘭 𝘦𝘲𝘶𝘪𝘱𝘮𝘦𝘯𝘵 𝘴𝘶𝘱𝘱𝘰𝘳𝘵 𝘧𝘰𝘳 𝘵𝘩𝘪𝘴 𝘳𝘦𝘴𝘦𝘢𝘳𝘤𝘩 𝘢𝘯𝘥 𝘢𝘴𝘴𝘪𝘴𝘵𝘦𝘥 𝘢𝘭𝘭 𝘳𝘦𝘴𝘦𝘢𝘳𝘤𝘩𝘦𝘳𝘴 𝘸𝘪𝘵𝘩 𝘦𝘲𝘶𝘪𝘱𝘮𝘦𝘯𝘵 𝘮𝘢𝘪𝘯𝘵𝘦𝘯𝘢𝘯𝘤𝘦 𝘢𝘯𝘥 𝘶𝘱𝘥𝘢𝘵𝘦s. 𝘛𝘩𝘦𝘴𝘦 𝘩𝘢𝘳𝘥𝘸𝘰𝘳𝘬𝘪𝘯𝘨 𝘢𝘯𝘥 𝘥𝘦𝘥𝘪𝘤𝘢𝘵𝘦𝘥 𝘴𝘵𝘢𝘧𝘧 𝘮𝘦𝘮𝘣𝘦𝘳𝘴 𝘴𝘩𝘢𝘳𝘦 𝘵𝘩𝘪𝘴 𝘩𝘰𝘯𝘰𝘳 𝘸𝘪𝘵𝘩 𝘶𝘴, 𝘪𝘯𝘤𝘭𝘶𝘥𝘪𝘯𝘨 𝘵𝘩𝘦 𝘧𝘰𝘭𝘭𝘰𝘸𝘪𝘯𝘨 𝘴𝘵𝘢𝘧𝘧: 𝙃𝙖𝙧𝙧𝙞𝙨𝙤𝙣 𝙎𝙞𝙢𝙧𝙖𝙡𝙡, 𝙏𝙤𝙢𝙖𝙨 𝙁𝙚𝙡𝙞𝙥𝙚 𝙇𝙡𝙖𝙣𝙤 𝙍𝙞𝙤𝙨, and 𝙍𝙖𝙣𝙙𝙮 𝙑𝙖𝙧𝙜𝙖𝙨. This study was supported by The Fundamental Research Funds for the Central Public Welfare Research Institutes (No. ZZ19-XRZ-113) and the CACMS Innovation Fund (No. ZB2025033)

## Supplementary Appendix 1 RNA Pathway Alterations in Air Pollution

### Label-Free Discovery of RNA Functional Pathways

TOGGLE was applied to analyze the effects of e-cigarette aerosol inhalation on mouse cardiac tissue using dataset GSE183614^1^. Taking the differentially expressed genes of adult mice exposed to e-cigarette smoke and fresh air as an example, they were divided into 6 groups. Except for Group 2, Group 3, Group 4 and Group 5, the remaining groups did not show significant enrichment in biological processes or KEGG signaling pathways and require further investigation (Figure S1).

Differentially expressed genes in adult mice exposed to e-cigarette smoke versus fresh air were categorized into 6 groups. Group 2 genes were primarily associated with glutamate secretion and regulation of cytoplasmic calcium ion concentration. Group 3 genes were linked to the regulation of cardiac contractility, while Group 4 genes were related to the Notch signaling pathway. Group 5 genes were associated with collagen fiber organization and cell division (Figure S1). The remaining groups did not show significant enrichment in biological processes or KEGG signaling pathways and require further investigation.

Similarly, differentially expressed genes in young mice exposed to e-cigarette smoke versus fresh air were categorized into 5 groups. Group 1 was primarily associated with immune and inflammatory pathways, including Lipid and Atherosclerosis, TNF signaling, and Cytokine-cytokine receptor interactions; Group 2 was linked to immune inflammation as well, with a focus on pathways related to vascular endothelial growth factor production, apoptosis, TNF signaling, and Cytokine-cytokine receptor interactions; Group 3 genes were mainly involved in the Calcium signaling pathway and cellular responses to interferon-β; Group 4 was associated with pathways such as p53 signaling, FoxO signaling, apoptosis, cell differentiation, and blood vessel maturation; Group 5 genes were related to mesenchymal cell proliferation, inflammation, and fibroblast proliferation.

Similarly, differentially expressed genes in young mice exposed to e-cigarette smoke versus adult mice exposed to e-cigarette smoke were categorized into 5 groups. Group 1 was primarily associated with immune and inflammatory pathways, including cytokine-cytokine receptor interactions, chemokine signaling, and leukocyte chemotaxis. Group 2 also involved immune inflammation, particularly cytokine-cytokine receptor interactions, TNF signaling, and inflammatory responses. Group 3 was primarily linked to the cell cycle and cell division pathways. Group 4 was associated with FoxO signaling, TNF signaling, and cell migration. Finally, Group 5 was involved in pathways related to the cell cycle, mitosis, p53 signaling, and NOD-like receptor signaling.

Similarly, differentially expressed genes in young mice exposed to fresh air versus adult mice exposed to fresh air were categorized into 4 groups. Group 1 was primarily associated with cytokine-cytokine receptor interactions and the cell cycle. Group 2 focused on cellular senescence and cell cycle-related pathways. Group 3 was linked to the cell cycle and cell division processes. Similarly, Group 4 was also related to cell cycle and cell division pathways.

The results indicate that exposure to e-cigarette aerosol significantly altered RNA pathways in the hearts of mice. These changes are reflected in the grouping of differentially expressed genes and their enrichment in biological functions and signaling pathways. More pronounced immune and inflammatory responses (e.g., TNF signaling) and cell cycle-related changes among young mice compared to mature adults suggested that the impact of e-cigarette use is greater at younger ages.

**Figure S1.**
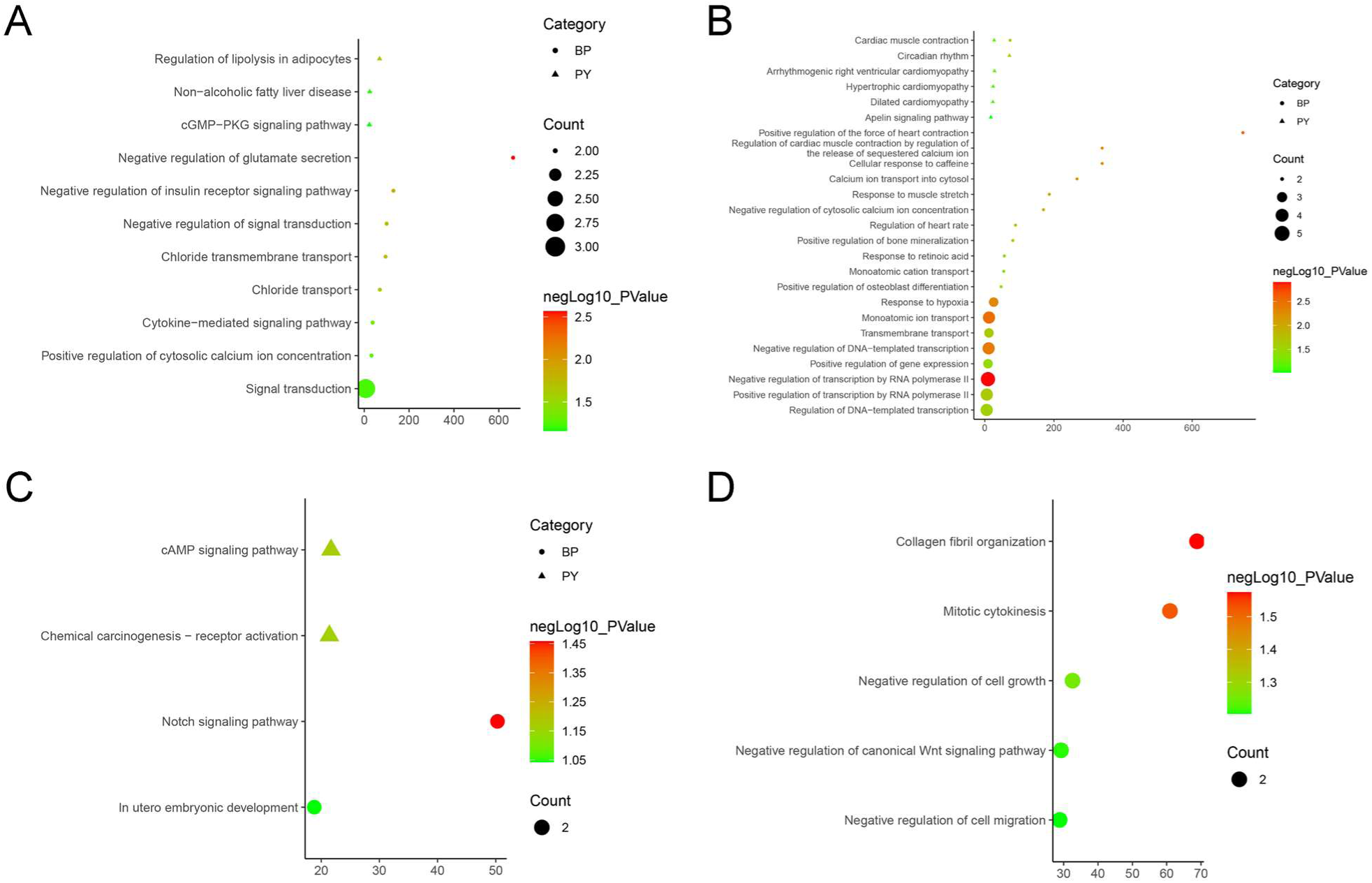
The Enrichment Results of Genes in Each Gene Group. A: Biological processes and pathways of Group 2; B: Biological processes and pathways of Group 3; C: Biological processes and pathways of Group 4; D: Biological processes of Group 5. (BP: biological processes; PY: signaling pathways. X-axis is fold enrichment)

## Supplementary Appendix 2 Cellular functional mapping

### Discovery of Signaling Pathways in the RNA Network

TOGGLE was applied to analyze single-nucleus RNA sequencing (snRNA-seq) and bulk RNA sequencing (bulk-RNA seq) data from a rat myocardial infarction (MI) model (dataset: GSE253768). After classifying the snRNA-seq data into various cell types, including fibroblasts, myofibroblasts, macrophages, T cells, and cardiomyocytes (Figure S2A), TOGGLE divided the fibroblasts and myofibroblasts into seven groups (Figure S2B). The MI group was predominantly represented by FibR1-G5, FibR1-G6, and FibR1-G7 (Figure S2C). The results of the inferred ligand-receptor interactions between these fibroblasts and T cells or macrophages (cell-cell communication) revealed differences in the strength and types of interactions, suggesting functional variations among these fibroblast subgroups (Figure S2D and S2E). Further analysis of the highly variable genes in the MI fibroblasts also identified eight distinct functional groups (Figure S2F). Additionally, TOGGLE was used to map bulk RNA-seq data onto the snRNA-seq dataset, revealing that the gene expression pattern of fibroblasts in the bulk RNA-seq MI group closely matched that of FibR2-G6 (highlighted in the red box in Figure S2G). This finding suggests that the bulk RNA-seq fibroblasts predominantly exhibit functions similar to those of FibR2-G6, which are associated with cell migration, adhesion, angiogenesis, and ECM-receptor interactions (Figure S2H).

Thus, TOGGLE can effectively combine bulk RNA-seq data with snRNA-seq to contextualize each, and which cell types and specific phenotypes undergo nuclear transcriptional changes that most closely represent overall tissue-level changes in transcription.

**Figure S2.**
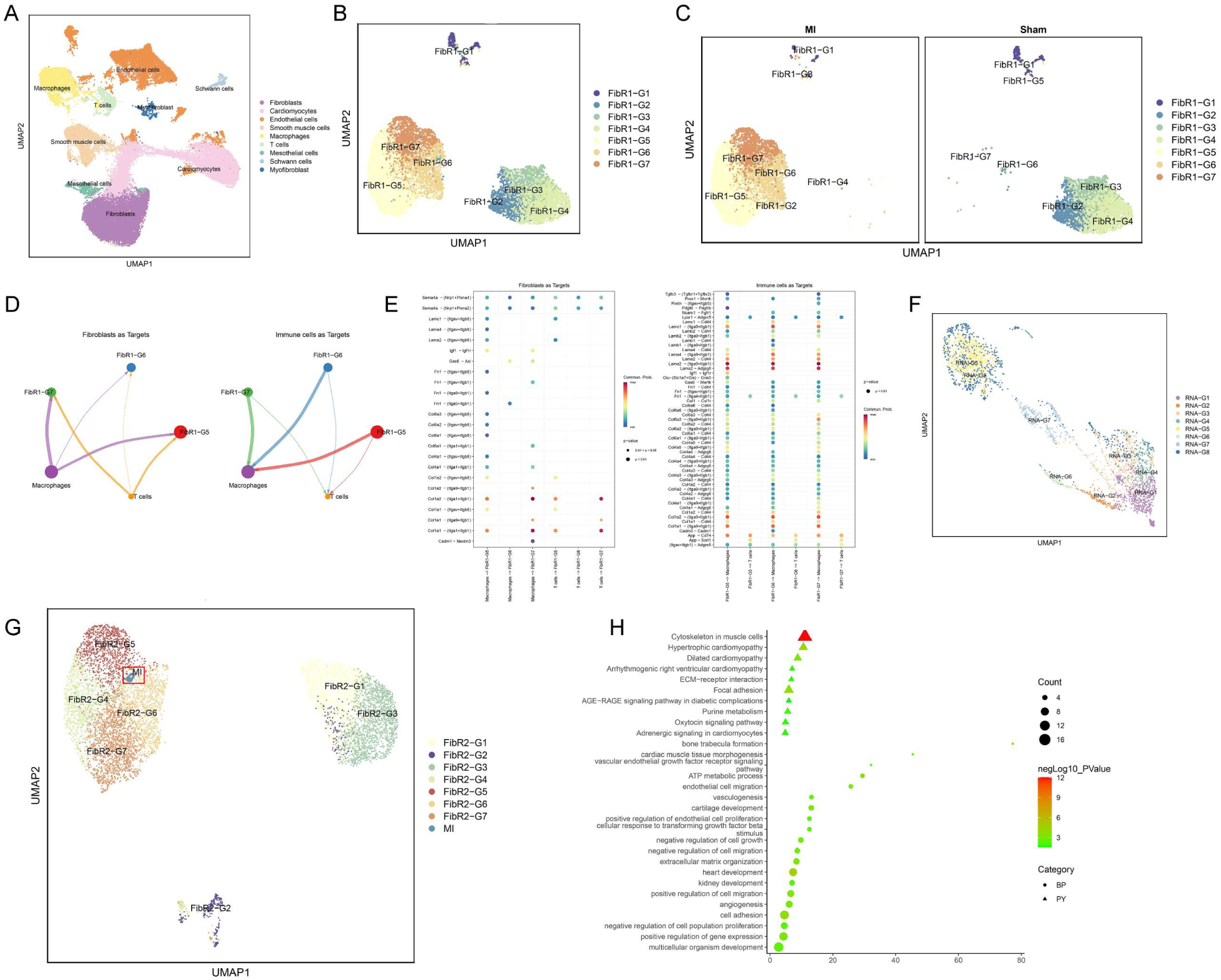
Analysis results of GSE253768 set. A: Cell types of snRNA-seq; B: Fibroblasts and myofibroblasts were divided into 7 groups after TOGGLE analysis; C: Distribution of 7 cell types in MI cohort and Sham cohort; D: Weight/strength diagram of cell communication between fibroblasts and immune cells; E: Bubble diagram of cell communication between fibroblasts and immune cells; F: RNA distribution of fibroblasts in the MI group; G: Mapping MI group bulk RNA-seq data onto UMAP of fibroblast snRNA-seq results; H: Enrichment results of FibR2-G6 (BP: biological processes; PY: signaling pathways. X-axis is fold enrichment).

## Supplementary Appendix 4 Functional details and site discovery of the ferroptosis RNA map

**Figure S7.**
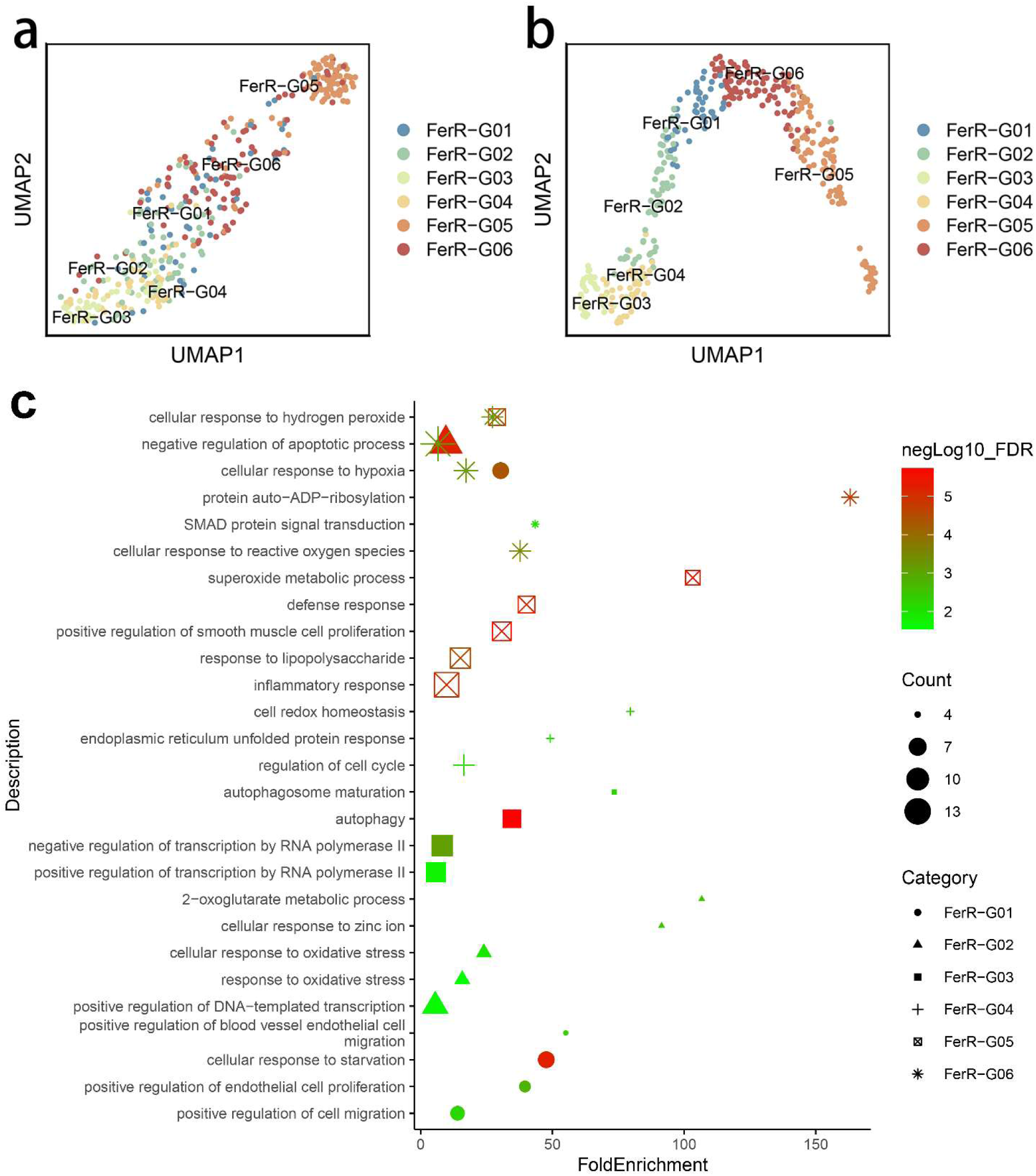
Up: RNA functional profiles from different data. a: RNA map of ferroptosis-related genes in Group R3-4 and R3-5 cells made by using single-cell expression matrix; b: RNA map made using the new method for the data in fig a. Down: c: Bubble chart of biological processes of ferroptosis-related genes in Group R3-4 and R3-5 cells

### Establish Graph Diffusion Functional Map, Functional discovery of RNA groups

We performed enrichment analysis on the RNA groupings presented in Figure S7a and S7b. The enrichment analysis results of FerR-G01 include cellular response to starvation (GO:0009267), cellular response to hypoxia (GO:0071456), positive regulation of blood vessel endothelial cell migration (GO:0043536), and positive regulation of cell migration (GO:0030335). These suggest that this RNA cluster may play a role in promoting angiogenesis in ischemic brain regions to counteract hypoxia caused by cerebral ischemia. Within this RNA group, we identified ferroptosis-related genes Nfs1, Osbpl9, and Pgd, which were not enriched in any of the related biological processes, implying their functions may be previously unrecognized. We hypothesize that these genes are functionally linked to the processes associated with FerR-G01.

FerR-G02 is associated with negative regulation of apoptotic processes (GO:0043066) and cellular response to oxidative stress (GO:0034599), indicating that this RNA cluster may be involved in both ferroptosis-related oxidative stress and apoptosis. Notably, several ferroptosis-related genes, Pvt1, Abhd12, Cbr1, Far1, and Sat1, were not enriched in the identified processes, suggesting their potential involvement in oxidative stress regulation.

FerR-G03 is associated with autophagy (GO:0006914), autophagosome maturation, and negative regulation of transcription by RNA polymerase II (GO:0000122), indicating a possible role in both ferroptosis and autophagy regulation. The genes Gstm1, Rpl8, Ubc, and Vdac2 within this group may also be functionally relevant to these processes.

FerR-G04 is linked to cell redox homeostasis (GO:0045454), regulation of the cell cycle (GO:0051726), and the endoplasmic reticulum unfolded protein response (GO:0030968), suggesting its involvement in ferroptosis-related oxidative stress. The genes Agpat3, Abcc5, Cd82, Emc2, Cs, and Psat1 within this cluster may also contribute to these functions.

FerR-G05 is associated with superoxide metabolic processes (GO:0006801), inflammatory responses (GO:0006954), and cellular response to hydrogen peroxide (GO:0070301), implying a connection to oxidative stress in ferroptosis. The genes Amn, Cdca3, Cltrn, Hcar1, Plin4, and Galnt14 may also play roles in these pathways.

FerR-G06 is associated with cellular response to reactive oxygen species (GO:0034614) and SMAD protein signal transduction (GO:0060395), suggesting a link between ferroptosis-related oxidative stress and the SMAD signaling pathway. Several genes within this cluster, including Cisd3, Hddc3, Lyrm1, Rbms1, Ahcy, Adam23, Aqp5, Ccdc6, Eno3, Gria3, Gpt2, Gstz1, Ints2, Klhdc3, Lpcat3, Mib2, Phkg2, Parp4, and Pdss2, may also be functionally relevant (Figure S7).

The genes not enriched in the identified signaling pathways may still be functionally relevant to their respective RNA clusters, potentially revealing new mechanisms in ferroptosis. For example, in FerR-G01, Nfs1 plays a crucial role in the biosynthesis of iron-sulfur (Fe-S) clusters, which are essential for the function of mitochondrial respiratory chain complexes^9^. Ischemia-induced hypoxia disrupts mitochondrial function, reducing ATP production and causing cellular damage. Nfs1 may regulate ferroptosis by maintaining Fe-S cluster homeostasis, thereby modulating iron metabolism and antioxidant capacity. Osbpl9 is a key regulator of cholesterol and phospholipid exchange, playing an important role in intracellular lipid transport and signaling^10^. It may promote endothelial cell migration and neovascularization in ischemic brain regions by modulating the lipid microenvironment and associated signaling pathways. Pgd participates in the pentose phosphate pathway (PPP), primarily generating NADPH to maintain cellular antioxidant capacity^11^. Under hypoxic conditions, elevated reactive oxygen species (ROS) levels contribute to oxidative stress. Pgd may counteract ferroptosis by enhancing antioxidant defenses, protecting cells from oxidative damage.

In FerR-G02, Pvt1 is a long non-coding RNA (lncRNA) known to regulate various physiological processes, including cell proliferation, apoptosis, and oxidative stress response^12^. In the context of ferroptosis, Pvt1 may influence iron accumulation and ROS generation by modulating stress-response genes. Abhd12, an enzyme involved in lipid metabolism, plays a key role in membrane lipid processing^13^. It may contribute to ferroptosis resistance by regulating fatty acid metabolism, stabilizing cellular membranes, and modulating intracellular lipid homeostasis. Cbr1 is an oxidoreductase enzyme that catalyzes the reduction of aldehyde compounds, protecting cells from oxidative damage ^14^. It may mitigate ferroptosis by reducing ROS levels and limiting oxidative injury. Far1 is involved in peroxisome biogenesis^15^, suggesting a potential role in oxidative stress regulation and ferroptosis modulation. Sat1, a key gene in the polyamine metabolism pathway, facilitates polyamine acetylation, which regulates intracellular polyamine levels^16^. Given that polyamines exhibit antioxidant properties, Sat1 may counteract oxidative stress, thereby influencing ferroptosis susceptibility.

## Supplementary Appendix 5 Functional details and site discovery of the aNSC/qNSC RNA map

### Functional discovery of RNA groups, Discovery of epigenetic control of energy metabolism

Additionally, we performed enrichment analysis on the RNA clusters derived from the top 1,000 highly variable genes in aNSCs, as shown in Figures S8a and S8b. Our findings indicate that aNSCR-G01 is primarily associated with cell differentiation (GO:0030154), while aNSCR-G02 is linked to the regulation of transcription by RNA polymerase II (GO:0006357). aNSCR-G03 is enriched in chromosome segregation (GO:0007059) and proteolysis (GO:0006508), whereas aNSCR-G04 is involved in DNA repair (GO:0006281) and cell division (GO:0051301). aNSCR-G05 is related to cell migration (GO:0016477), regulation of gene expression (GO:0010468), and positive regulation of cell growth (GO:0030307), while aNSCR-G06 is enriched in mRNA processing (GO:0006397), RNA splicing (GO:0008380), and chromatin remodeling (GO:0006338). These functional associations suggest that the identified RNA clusters are predominantly involved in cell proliferation, DNA replication, and RNA transcription, aligning well with the in vivo characteristics of aNSCs.

We also performed enrichment analysis on the RNA clusters derived from the top 1,000 highly variable genes in qNSCs, as shown in Figures S8c and S8d. Our findings indicate that qNSCR-G01 is primarily associated with centriole replication (GO:0007099) and regulation of the mitotic cell cycle (GO:0007346), while qNSCR-G02 is linked to cell adhesion (GO:0007155) and the apoptotic process (GO:0006915). qNSCR-G03 is enriched in regulation of transcription by RNA polymerase II (GO:0006357) and endoplasmic reticulum to Golgi vesicle-mediated transport (GO:0006888), whereas qNSCR-G04 is involved in proton motive force-driven mitochondrial ATP synthesis (GO:0042776) and proton transmembrane transport (GO:1902600). These functional associations suggest that the identified RNA clusters are involved in regulating the cell cycle, cell adhesion, intracellular protein synthesis, and cellular energy metabolism—characteristics typical of non-proliferative cells, aligning well with the in vivo behavior of qNSCs.

**Figure S8.**
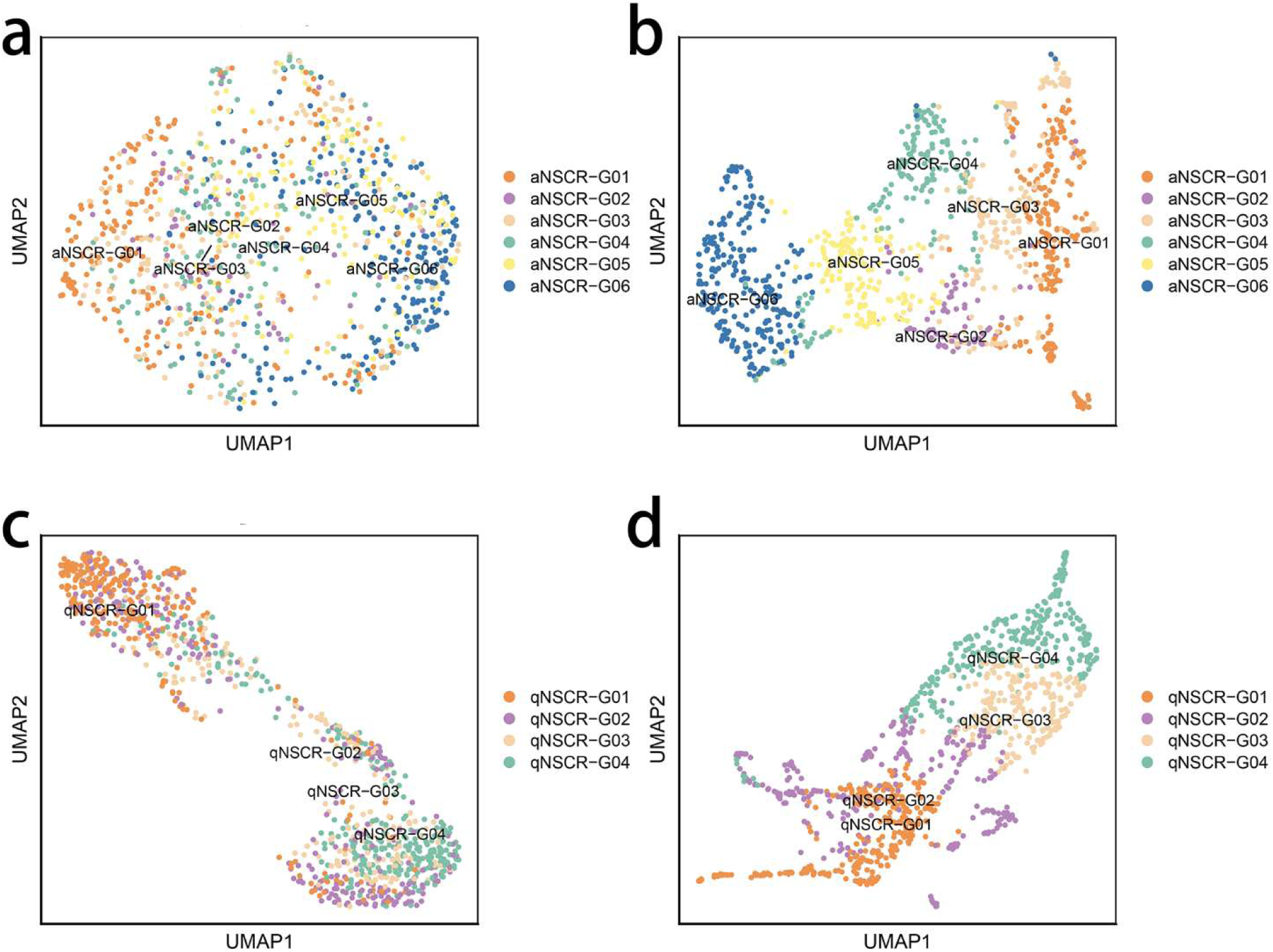
RNA functional profiles from different data. a: RNA map of 1000 highly variable genes in aNSC cells made by using single-cell expression matrix; b: RNA map made using the new method for the data in c; c: RNA map of 1000 highly variable genes in qNSC cells made by using single-cell expression matrix; d: RNA map made using the new method for the data in e

To investigate the epigenetic regulation of energy metabolism, we conducted a multimodal analysis of NSC scRNA-seq and single-cell epigenomic data from Kremer et al.^17^ First, we extracted aNSCs and qNSC2 based on the labels provided by Kremer et al.^17^ and analyzed the data of GSE209656 (scRNA-seq data) using TOGGLE (Figure S9a and S9b). Next, we classified the cells into two groups based on the median expression level of mitochondrial RNA: a high mitochondrial RNA expression group (High-MT) and a low mitochondrial RNA expression group (Low-MT) (Figure S9d and S9e). Our analysis revealed that NeuG04 primarily consists of metabolically active cells, predominantly aNSCs, while NeuG01 is composed mainly of metabolically inactive cells, predominantly qNSCs (Figure S9c and S9f). The remaining cell groups fall between these two cell groups and may represent transitional states.

**Figure S9.**
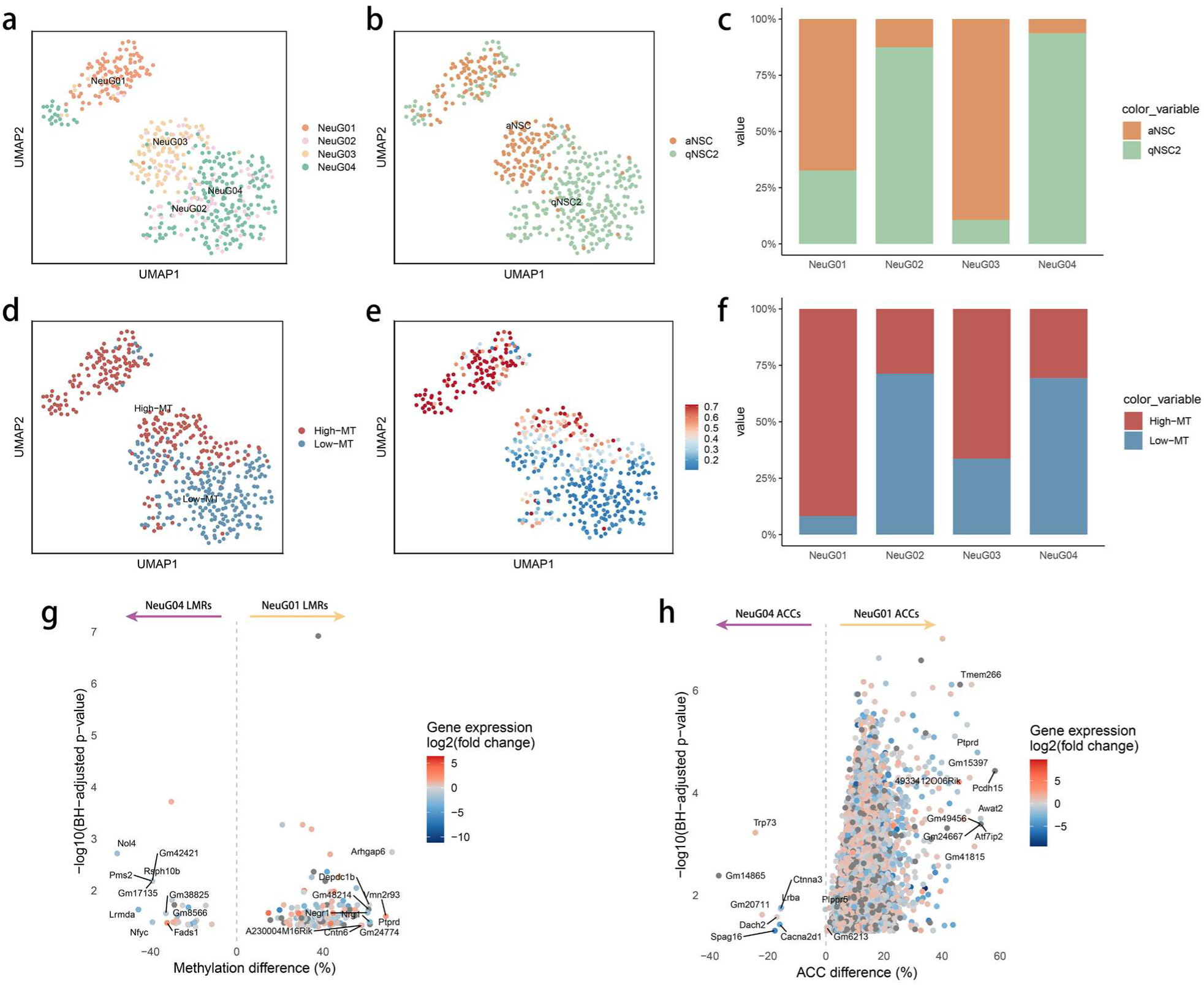
Results of epigenetic analysis of aNSCs and qNSCs. a: Cell populations after TOGGLE analysis; b: Labels provided by Kremer et al.; c: The proportion of aNSC and qNSC2 in each cell group; d: Cells are divided into High-MT and Low-MT according to the expression of mitochondrial genes; e: Mitochondrial gene expression in cells; f: The proportion of high-MT cells and low-MT cells in each cell group; g: Volcano plot of LMRs, tested for differential methylation between NeuG01 and NeuG04 (two-sided Wilcoxon test), LMRs are colored according to the differential expression of the nearest gene. h: Volcano plot of high ACC resgions, tested for differential ACC between NeuG01 and NeuG04 (two-sided Wilcoxon test), LMRs are colored according to the differential expression of the nearest gene

Then, we analyzed the single-cell epigenomes (GSE211786) dataset from Kremer et al.^17^ based on the cell grouping results to analyze the epigenetic differences between the high-MT-dominated cell population (NeuG01) and the low-MT-dominated cell population (NeuG04). First, we extracted the demethylation data for cells classified as NeuG01 and NeuG04 and analyzed the variably methylated regions (VMRs) using Kremer et al.’s approach. We then calculated average DNA methylation (meth) and chromatin accessibility (ACC).

To identify differentially methylated regions between the two groups, we performed a two-sided Wilcoxon rank sum test on the VMRs, considering only regions with genomic coverage in at least six cells per group. VMRs with a Benjamini–Hochberg-adjusted P-value < 0.05 were labeled as lowly methylated regions (LMRs) or high ACC regions. To annotate the genes, all LMRs overlapping a gene body were assigned to that gene.

The results revealed that compared to NeuG04, NeuG01 exhibited higher LMR and high ACC regions (Figure S9g and S9h). The LMRs in NeuG01 were enriched in genes associated with stem cell differentiation (GO:0048863) and positive regulation of transcription by RNA polymerase II (GO:0045944), while high ACC regions in NeuG01 were linked to genes involved in cell adhesion (GO:0007155), chromatin remodeling (GO:0006338), axon guidance (GO:0007411), positive regulation of transcription by RNA polymerase II (GO:0045944), actin cytoskeleton organization (GO:0030036), nervous system development (GO:0007399), cell migration (GO:0016477), positive regulation of DNA-templated transcription (GO:0045893), and cell differentiation (GO:0030154). These findings suggest that the epigenetic modifications in NeuG01 cells are primarily associated with cell proliferation, differentiation, and migration—processes that demand high energy consumption.

Conversely, the LMRs in NeuG04 were linked to genes involved in intracellular signal transduction (GO:0035556), nervous system development (GO:0007399), regulation of synapse assembly (GO:0051963), cell adhesion (GO:0007155), synaptic membrane adhesion (GO:0099560), signal transduction (GO:0007165), long-term memory (GO:0007616), negative regulation of the cell cycle (GO:0045786), negative regulation of cell migration (GO:0030336), and negative regulation of DNA-templated transcription (GO:0045892). These functions are negatively correlated with cell proliferation and division, indicating that NeuG04 cells are transitioning into a quiescent state. A detailed list of LMRs, high ACC regions, and their corresponding genes for NeuG01 and NeuG04 is provided in Table S3.

Since NeuG01 is primarily composed of aNSC cells, while NeuG04 consists mainly of qNSC2 cells, we can conclude the following: (1) During the transition from qNSC2 to aNSC, key genomic regions associated with cell proliferation and energy metabolism undergo significant demethylation (Figure S9g), accompanied by a notable increase in chromatin accessibility (Figure S9h). (2) In cell populations dominated by aNSCs, the proportion of high mitochondrial gene expression (High-MT) cells is significantly higher than in qNSC2 (Figures S9c, S9d, and S9e), further indicating that aNSCs exhibit higher metabolic activity.

In summary, we propose that during the qNSC2-to-aNSC transition, localized DNA demethylation and chromatin opening activate the expression of genes involved in cell proliferation and energy metabolism, leading to a gradual increase in metabolic activity. This epigenetic modification reflects differences in “cellular memory” between these two cell types and may be a key mechanism underlying their distinct mitochondrial energy metabolism states.

## Supplementary Appendix 6 Animal Processing

### 1 scRNA-seq Data Processing

Cells from the GSE232429 dataset^2^ with 500 < nFeature_RNA < 8000, mitochondrial gene expression percentage (mt_percent) < 30%, red blood cell gene proportion < 5%, and nCount_RNA < 98th percentile were retained for further analysis. FindMarker Function was used to perform a preliminary analysis of this dataset to identify differentially expressed genes. Differentially expressed genes were screened out using the criteria of |Log2FC|≥0.25 and FDR<0.05. Cellchat was used for cell-cell communication analysis^18^.

### 2 Experimental Materials and Methods

#### 2.1 Experimental Materials

##### (1) Experimental Animal

Fifty male specific pathogen free Sprague-Dawley (SD) rats, aged 6 to 8 weeks and weighing 250-280 g, were provided by the Experimental Animal Center of Hunan University of Chinese Medicine (Animal Quality Certificate No.: 430727211100405318). The rats were purchased from Hunan Sileike Jingda Experimental Animal Co., Ltd. [License No.: SCXK (Xiang) 2019-0004]. They were housed in the Experimental Animal Center of Hunan University of Chinese Medicine [License No.: SYXK (Xiang) 2019-0009] under standard conditions, with a room temperature of (24±1)°C, humidity of approximately 55%, and a 12-hour light/dark cycle. After a one-week acclimation period, the animals were used for the experiments. All animals’ care and experimental procedures were approved by the Animal Ethics Committee of Hunan University of Chinese Medicine (Approve number: LL2019092009) and were in accordance with the National Institute of Health’s Guide for the Care and Use of Laboratory Animals.

##### (2) Reagents

Paraformaldehyde (Batch No.: G1101) was purchased from Servicebio, and anhydrous ethanol (Batch No.: 10009259) from China National Pharmaceutical Group Chemical Reagent Co., Ltd. Electron microscopy fixative (Cat. No. G1102) was purchased from Wuhan Guge Biotechnology Co., Ltd. Transferrin (Trf) polyclonal antibody (Cat. No.: 17435-1-AP), Bax polyclonal antibody (Cat. No.: 50599-2-Ig) and IκBα polyclonal antibody (Cat. No.: 10268-1-AP) were obtained from Proteintech. Casp-3 antibody (Cat. No.: 9661L) was purchased from Cell Signaling. Sirt1 antibody (Cat. No. ab189494) was purchased from Abcam. Transferrin receptor (Tfrc) polyclonal antibody (Cat. No.: A5865), and Rela (p65) polyclonal antibody (Cat. No.: A2547) were sourced from ABclonal. P-smad (Cat. No.: AF8313) was acquired from Affinity Bioscience. Gpx4 antibody (Cat. No.: AF301385), Acsl4 polyclonal antibody (Cat. No.: AFW6826) and Fsp1 polyclonal antibody (Cat. No.: AFW21808) were purchased from AiFangbio. DAPI (Cat. No.: ab228549) was obtained from Abcam. Deferiprone tablets (specification: 50 mg/tablet, batch number: H20140379) was purchased form Apotex Inc.

##### (3) Instruments

The optical microscope (Axioscope-5) was purchased from Zeiss, Germany. The pathology microtome (RM2016) and Leica blades (LEICA-819) were both obtained from Leica Instruments Co., Ltd. Transmission electron microscope (HT7700) was purchased from Hitachi Inc.

#### 2.2 Animal Grouping, Modeling and Intervention

Fifty SD rats were randomly assigned into three groups: the Sham operation group, the MCAO group and the MCAO+Deferiprone group, with five rats in each group. The MCAO model was established using a modified reversible middle cerebral artery suture method^19^ for the rats in the MCAO group and the MCAO+Deferiprone group. Rats were fasted for 12 hours prior to surgery. During surgery, the right common carotid artery and external carotid artery were exposed and clamped, and a suture was inserted into the internal carotid artery via the common carotid artery to block the origin of the right middle cerebral artery. The suture was advanced approximately 18.5±0.5 mm until resistance was felt, indicating the correct position. The suture was then fixed to both the common and internal carotid arteries, and the incision was closed. Throughout the procedure, care was taken to maintain body temperature, and after surgery, the animals were returned to normal feeding conditions. In the Sham group, no suture was inserted, and the animals were kept under normal conditions.

Successful model establishment was confirmed when the rats exhibited left-sided limb paralysis, instability when standing, or circling behavior when lifted by the tail. Neurological function was scored on a scale of 1 to 3, with non-successful models excluded. The neurological scoring system was based on the Zea Longa scale^19^ as shown in Table S4.

Deferiprone tablets were dissolved in a 0.9% sodium chloride solution. Following model establishment, rats in the deferiprone group received oral administration of deferiprone at a dose of 125 mg/kg per day. Rats in the sham operation group and the MCAO group were given an equivalent volume of 0.9% sodium chloride solution. Treatments were administered once daily for three consecutive days.

#### 2.3 Neuronal morphological changes observation by hematoxylin-eosin (HE) staining

The brain tissue was perfused with pre-cooled 0.9% sodium chloride solution, followed by fixation in 4% paraformaldehyde at 4°C. The tissue was then coronally sectioned into 4 μm thick slices. The sections were deparaffinized, rehydrated, and stained with hematoxylin for 5 minutes. After differentiation and rinsing with water, the sections were blued with distilled water, dehydrated through a graded ethanol series, and counterstained with eosin for 5 minutes. Following dehydration with ethanol and clearing with xylene, coverslips were applied and sealed with neutral balsam. Morphological observations and image acquisition were performed under a light microscope.

#### 2.4 Transmission electron microscopy

The affected brain tissue was sectioned into 1 mm-thick slices. Samples were immersed in electron microscopy fixative at 4°C overnight, followed by fixation in 1% osmium tetroxide at 4°C for 2 hours. The samples were stained with 1% uranyl acetate for 1 hour, dehydrated, and embedded in epoxy resin. Ultrathin sections were prepared using an ultramicrotome, stained with uranyl acetate and lead citrate, and examined under a transmission electron microscope.

#### 2.5 Immunohistochemical Staining for Acsl4, Gpx4 and Fsp1 Protein Expression Levels

Paraffin-embedded tissue sections (5 μm) were deparaffinized and rehydrated. Endogenous enzymes were inactivated by incubating the sections in 3% hydrogen peroxide at room temperature for 10 minutes, followed by two washes with distilled water. Antigen retrieval was performed by immersing the sections in 0.01 mol/L citrate buffer (pH 6.0) and heating in a microwave until boiling, then turning off the power and repeating the process after a 5-minute interval. After cooling, the sections were washed twice with PBS buffer (pH 7.2-7.6) for 3 minutes each. Blocking solution was added and incubated at room temperature for 20 minutes, with excess liquid removed without washing. Primary antibodies were applied and incubated at 37°C for 2 hours, followed by three washes with PBS buffer (pH 7.2-7.6) for 3 minutes each. Biotinylated secondary antibody solution was added and incubated at 20-37°C for 20 minutes, followed by another round of PBS washing. Streptavidin-HRP was applied and incubated at 20-37°C for 20 minutes, followed by washing in PBS as described earlier. DAB was used for color development, with reaction time controlled under a microscope (5-30 minutes). Sections were lightly counterstained with hematoxylin, dehydrated, cleared, and mounted with neutral balsam for microscopic observation of Acsl4 and Gpx4 protein expression. Images from five randomly selected fields of view from different samples were captured, and average optical density was analyzed using ImageJ software.

#### 2.6 Immunofluorescence Staining for Bax, IκBα, Casp3, p-Smad, p65 (Rela), Tfrc, Trf, and Sirt1 Protein Expression Levels

Paraffin sections were deparaffinized and rehydrated, followed by membrane permeabilization with 0.3% Triton X-100 for 5 minutes. The sections were then blocked with 5% BSA in PBST at room temperature for 1 hour. Primary antibodies, diluted in PBST, were applied, and the sections were incubated overnight at 4°C. After washing with PBST, fluorescent secondary antibodies were added, and the sections were incubated in the dark at room temperature for 1 hour. The secondary antibodies were washed off, and the slides were mounted with anti-fade mounting medium containing DAPI. The next day, images were captured using a panoramic tissue cell quantification system at 40x magnification. Images from five randomly selected fields of view from different samples were captured and analyzed using ImageJ software to assess the average fluorescence intensity.

#### 3 Statistical Analysis

All data are presented as Mean±SD, and statistical analysis and graphing were performed using GraphPad Prism 9.0. Non-parametric tests were used to analyze average fluorescence intensity and average optical density. A P-value of less than 0.05 was considered statistically significant.

